# Negative selection may cause grossly altered but broadly stable karyotypes in metastatic colorectal cancer

**DOI:** 10.1101/2020.03.26.007138

**Authors:** William Cross, Salpie Nowinski, George D Cresswell, Maximilian Mossner, Abhirup Banerjee, Bingxin Lu, Marc Williams, Georgios Vlachogiannis, Laura Gay, Ann-Marie Baker, Christopher Kimberley, Freddie Whiting, Hayley Belnoue-Davis, Pierre Martinez, Maria Traki, Viola Walther, Kane Smith, Javier Fernandez-Mateos, Erika Yara, Erica Oliveira, Salvatore Milite, Giulio Caravagna, Chela James, George Elia, Alison Berner, Ryan Changho Choi, Pradeep Ramagiri, Ritika Chauhan, Nik Matthews, Jamie Murphy, Anthony Antoniou, Susan Clark, Miriam Mitchison, Jo-Anne Chin Aleong, Enric Domingo, Inmaculada Spiteri, Stuart AC McDonald, Darryl Shibata, Miangela M Lacle, Lai Mun Wang, Morgan Moorghen, Ian PM Tomlinson, Marco Novelli, Marnix Jansen, Alan Watson, Nicholas A Wright, John Bridgewater, Manuel Rodriguez-Justo, Chris P Barnes, Hemant Kocher, Simon J Leedham, Andrea Sottoriva, Trevor A Graham

## Abstract

Aneuploidy, the loss and gain of whole and part chromosomes, is near-ubiquitous in cancer genomes and likely defines cancer cell biology. However, the temporal evolutionary dynamics that select for aneuploidy remain uncharacterised. Here we perform longitudinal genomic analysis of 755 samples from a total of 167 patients with colorectal-derived neoplastic lesions that represent distinct stages of tumour evolution through metastasis and treatment. Adenomas typically had few copy number alterations (CNAs) and most were subclonal, whereas cancers had many clonal CNAs, suggesting that progression goes through a CNA bottleneck. Individual CRC glands from the same tumour typically had very similar karyotypes, despite evidence of ongoing instability at the cell level in patient tumours, cell lines and organoids. CNAs in metastatic lesions sampled from liver and other organs, after chemotherapy or targeted therapies, and in late recurrences were typically similar to the primary tumour. Mathematical modelling and statistical inference indicated that these data are consistent with the action of negative selection on CNAs that ‘traps’ cancer cell genomes on a fitness peak defined by the specific pattern of chromosomal aberrations. These data suggest that the initial progression of colorectal cancer requires the traversal of a rugged fitness landscape and subsequent CNA evolution, including metastatic dissemination and therapeutic resistance, is constrained by negative selection.

## Introduction

Most cancer genomes are aneuploid^1^. This is the result of gains and losses of whole or part chromosomes which are attained through a variety of mechanisms that include chromosome mis-segregation, aberrant double strand break repair, and genome doubling^2^. In colorectal cancer (CRC), approximately 85% of cancers are classified as chromosomally unstable (CIN) by virtue of having an abnormal karyotype and aneuploidy^3^. The remaining ∼15% of cancers exhibit genetic instability at the mutational level and are classified as hypermutant, but exhibit limited chromosome aberrations.

Aneuploidy likely has a causal role in cancer evolution. First, the observed ubiquity of aneuploidy in cancers and the prevalence of recurrent chromosome copy number alterations (CNAs) is suggestive that some CNAs are positively selected^4,5^. Second, aneuploidy is predictive of progression to cancer in premalignant diseases such as Barrett oesophagus^6^ and ulcerative colitis^7^. Third, in established cancers, individual CNAs have prognostic value over-and-above single nucleotide alterations in key cancer driver genes^8^. Fourth, ploidy has been shown to have prognostic value in many types of established cancer^9^, and notably within-tumour heterogeneity of CNAs, as principally measured by the number of lineages with distinct karyotypes, is prognostic pan-cancer^10^. In lung cancer, the diversity of CNAs but not single nucleotide variants (SNVs) has prognostic value^11^.

Aneuploidy should also be subject to negative selection because the loss or gain of large genomic regions will affect the expression of thousands of genes and regulatory elements, some of which are likely to adversely affect tumour cell viability^12^. Moreover, it is likely that some chromosomal conformations prevent correct chromosomes segregation, resulting in cell death. Chromosomal-scale alterations are therefore expected to result in trade-offs between the dosages of positively- and negatively-selected elements^13^, potentially leading to an apparently stable aneuploid genome. Chromosomal changes may also have a buffering effect against deleterious point mutations^14^. In such an evolutionary regimen, most alterations are either neutral or negatively selected^15^. Indeed, studies show that the burden of CNAs is non-monotonically related with prognosis across cancer types^10^ (first observed in breast cancer^16^). As such, cancers carrying an intermediate level of alterations (‘just right’ aneuploidy) are associated with worse prognosis than cancers with lower or higher levels of aneuploidy. Observing negative selection may be complicated by the fact that less fit variants are effectively removed from the population and do not come to clinical prominence.

The spatial-temporal evolutionary dynamics that produce the aneuploidy observed in cancer genomes remain poorly understood, in part because longitudinal measurement is often not clinically-feasible and is compounded by the challenge of reconstructing ancestral history of chromosome gains and losses^17^. CRC presents a unique opportunity to track clonal evolution over space and time: pre-cancerous lesions (colorectal adenomas, CRAs) are occasionally ‘caught in the act’ of transformation and are found to contain a small focus of cancer (early CRCs). CRC frequently metastasizes to the liver, and repeated hepatic metastatectomy can provide a source of longitudinally sampled tumour material amenable for molecular analysis.

## Results

### Benign-to-malignant transformation through a CNA bottleneck

Here we investigated the evolution of aneuploidy in a large cross-sectional and longitudinally followed cohort of adenomas and cancers. Previously we observed that the degree of aneuploidy was much higher in a small cohort of (malignant) colorectal cancers than (benign) adenomas^18^. To validate this observation, we performed multi-region spatial sampling and genome-wide copy number alteration analysis (by shallow whole genome sequencing [sWGS] or SNP-array) in n=139 archival colorectal lesions (19 adenomas and 81 stage II/III CRCs, termed “regular CRCs”), and 39 early CRCs of which 23/39 (59%) contained residual adenoma (Fig 1A). In early CRCs, adenoma and carcinoma regions were separately subsampled (minimum 2 regions per neoplasm). The percentage of genome altered (PGA) by a CNA was lowest in adenomas and highest in regular CRCs (8.5% and 27.2% respectively, p=8.4×10^−14^) and early CRCs (25.3%; Fig 1B). Gains were more prevalent than losses in adenomas (p=0.0052; Fig S1A) whereas they occurred at similar frequency relative to baseline ploidy in cancers (p=0.75, p=0.32, respectively; Fig S1A). Which CNA events tended to be clonal or subclonal depended on tumour stage (Fig S1B).

**Figure 1:**
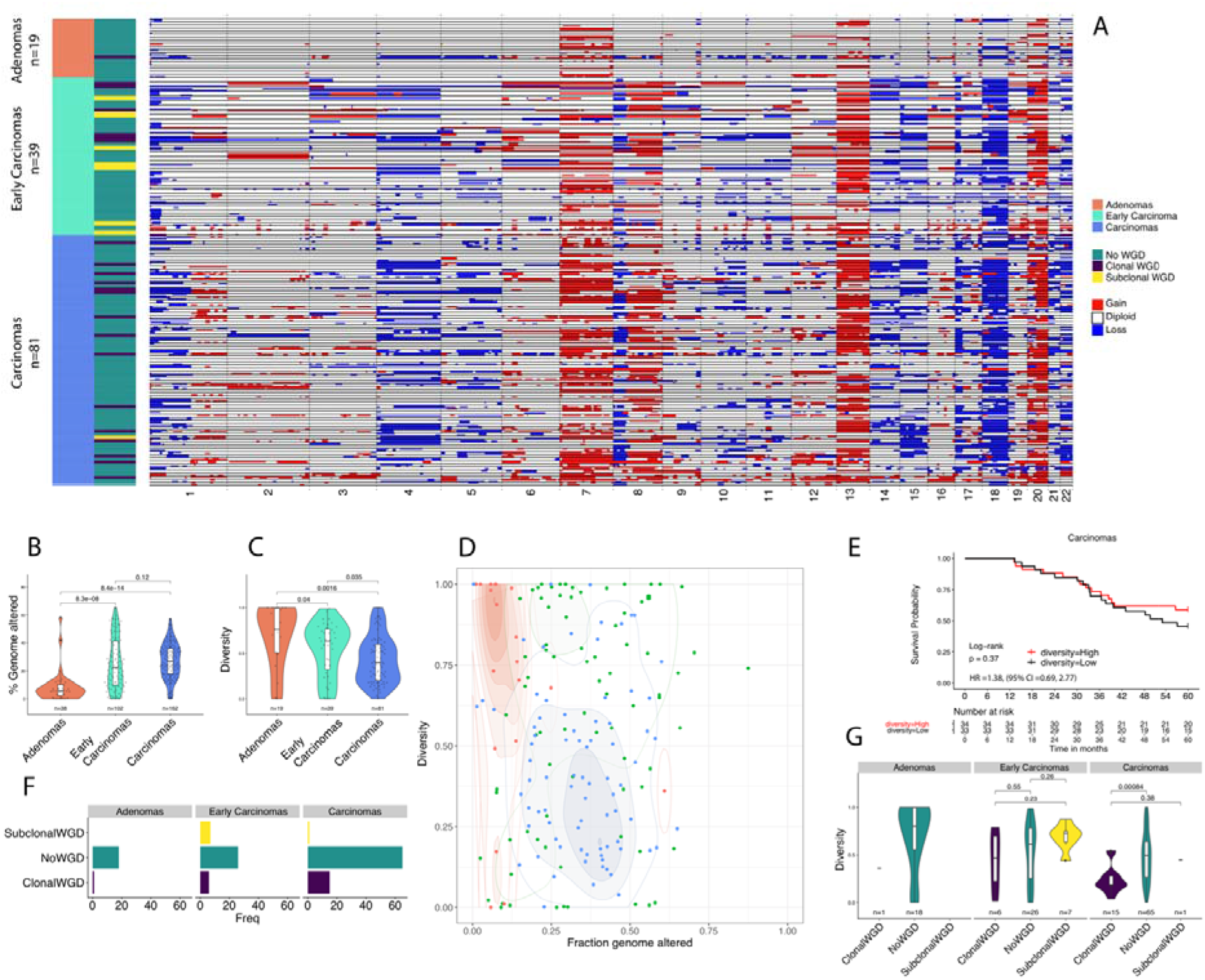
Aneuploidy increases through colorectal cancer progression, and clonal diversity decreases. **(A)** Heatmap showing copy number alterations (CNAs) in adenomas (orange; top), early cancers (green; middle) and established cancers (blue; bottom). Thin black lines delineate cases, within black lines are multiple regions of each case. Genome doubling (GD) status is shown on the left of the heatmap. Red=gain, blue=loss. **(B)** Percentage genome altered (PGA) by adenomas, early cancers and then later stage cancers. PGA increases through progression. **(C)** The proportion of CNAs that are subclonal decreases through progression from adenomas to carcinomas. **(D)** Phase space of PGA versus proportion of subclonal CNAs that are subclonal. Adenomas (orange) have low PGA but high CNA subclonality, whereas CRCs (blue) have high PGA but low CNA subclonality. Early cancers (green) are more disparate in PGA and CNA subclonality. **(E)** The proportion of CNAs that are subclonal is not significantly associated with overall survival in CRCs (Kaplan Meier analysis, groups split on the median proportion of subclonal CNAs across the cohort). (F) Frequency of genome doubling (GD) by tumour stage. GD is very rare in adenomas, more common in later stage cancers where it is usually found to be clonal. **(G)** The proportion of CNAs that are subclonal by tumour stage and genome doubling status. Clonal genome doubling is associated with a lower proportion of subclonal CNAs.

CNA subclonal diversity (the proportion of total altered genome not altered in all samples from a case; Methods) was greatest in adenomas and low in regular CRCs (p=0.0016; Fig 1C; Table S1; Table S2). Different measures to quantify CNA subclonality were highly correlated (Fig S2A). This suggests that progression from benign to malignant tumours goes through an aneuploidy-divergence ‘phase space’ from low-aneuploidy/high-diversity to high-aneuploidy/low-diversity (Fig 1D), indicating a selective genetic bottleneck acting on CNAs. Consistently, within the early CRCs, the fraction of subclonal CNAs was higher between the adenoma components than between cancer components (p=1.2×10^−6^; Fig S3A). Moreover, subclonal CNA fraction in cancer components of early CRCs was significantly lower than in regular CRCs (p=0.046; Fig S3A; Fig S3B).

We trained a classifier to infer the presence of genome doubling (GD) from shallow WGS data (Methods; Fig S4). GD was present in only a single adenoma and was clonal (Fig 1F). In regular CRCs, GD was more common (16/81, 18%) and was almost always clonal (15/16, 94%; Fig 1F). Within early CRCs, no adenoma component had GD, whereas a third of the early cancers (13/39) had evidence of GD (n=13/39; Fisher’s test p=0.0011) and typically this was clonal (11/13, 85%). These observations suggest that GD occurs at, or shortly prior to, the time of invasion. The proportion of subclonal CNAs was lower in GD than non-GD CRCs (p=0.00084; Fig 1F) and divergent CNAs were significantly shorter in length (p=0.015; Fig S5A). Therefore, following GD the aneuploid genome accrues little further alteration within primary CRCs, suggesting the process of genome doubling produces a fit genotype which drives passage through the CNA bottleneck.

We investigated the relationship between CNA diversity and clinical outcome in regular CRCs. The diversity of pre-existing adaptive phenotypes in a population should correlate with population evolvability, and consequently more diverse populations are more likely to harbour an individual pre-adapted to a new selective pressure. Accordingly, intra-tumour clonal diversity has been observed as a pan-cancer prognostic biomarker^10^ and it is prognostic in the premalignant lesion Barrett’s Oesophagus^19^. However, in our cohort, the proportion of subclonal CNAs was not associated with overall survival (comparison of cases above and below median proportion of subclonal CNAs ; p=0.37; Fig 1E; Table S3) Consequently, these data suggested that intra-tumour CNA diversity did not strongly reflect the heterogeneity of adaptive phenotypes in regular CRC, and was consistent with prior observations of a lack of positive selection experienced by CRC^20,21^. The presence of GD was also not associated with overall survival (p=0.56, Fig S5C).

### Limited CNA diversity at single gland resolution in CRCs

Previous reports using fluorescence in situ hybridisation (FISH) to measure CNAs at single cell resolution found widespread chromosomal instability in colorectal cancer cell lines^22^ and primary tumours^20^. To investigate CNA heterogeneity at higher resolution we exploited the fact that well-to-moderately differentiated CRCs are made up of a collection of glands, each composed of a few thousand cells recently derived from a common ancestor^20,23^. Glands can reasonably be considered as units of selection in CRC, as gland fission likely expands the cancer cell population^20^. We extracted a total of 307 individual colorectal cancer glands from the notional ‘left’ and ‘right’ opposite sides of an additional 6 regular CRCs (Fig 2A, Table S3) and performed high-depth exome sequencing (59 glands) or shallow whole genome sequencing (sWGS; 248 glands). In addition, we also analysed at least two bulk samples composed of numerous glands from each tumour.

**Figure 2:**
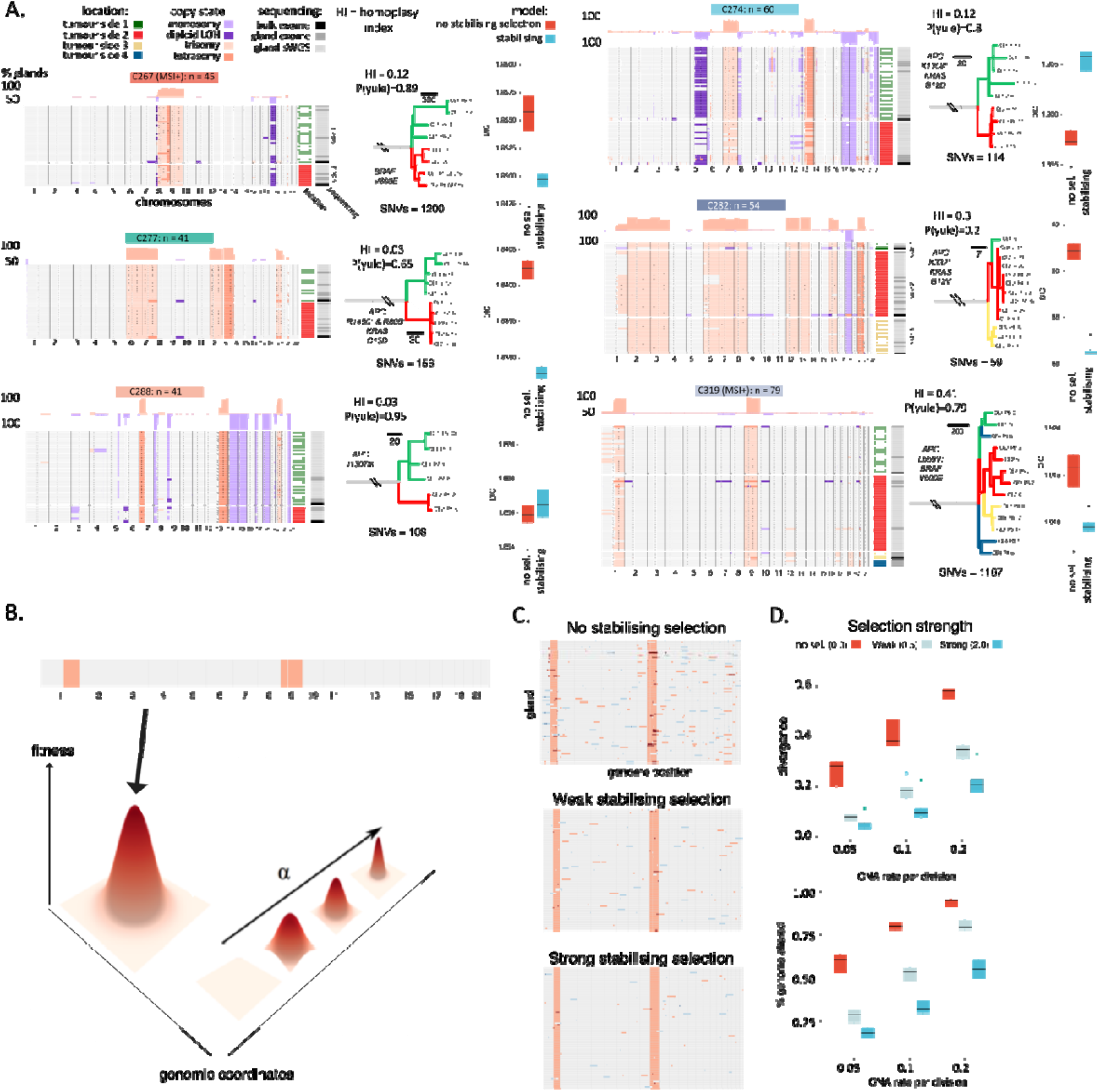
Lack of CNA heterogeneity at the level of individual colorectal cancer glands. **(A)** (left column) Heatmaps showing CNAs in individual glands in each of six CRCs. Glands were sequenced from arbitrary opposite sides of each CRC as indicated by the heatmap on the right-hand side. Red hue colours indicate gains, blue losses. (centre column) Phylogenetic trees reconstructed from individual gland single nucleotide alteration calls from exome sequenced glands. Trees are balanced (comparison with Yule model). Note the trunk length is not shown to scale, number above trunk represent number of clonal SNVs. Branches are coloured by notional tumour side where the gland was collected. HI=homoplasy index. (right column) Results of Bayesian inference to determine whether the observed CNA diversity is best supported by a model with or without stabilising selection. Model selection by deviance information criteria (DIC) suggests that the stabilising selection model is better supported for four patients (10 simulation replicates used for each boxplot). **(B)** Schematic showing shape of fitness landscape used in the computational simulations of stabilising selection. The model parameter determines the steepness of the single fitness peak in the landscape, at position defined by the above heatmap illustrating the optimal CNA pattern. **(C)** Heatmaps showing CNAs of the randomly sampled glands in one simulation as function of selective strength when CNA rate is fixed (0.2). The simulations were based on data from patient C319, and 77 glands were sampled when the population size reached 100000. **(D)** Genetic divergence (top) and percentage of the genome altered (bottom) as a function of CNA rate and selection strength (10 replicates for each case, the same simulation settings as in **C)**. Strong selection on an initially optimal clone supresses diversity and CNA accumulation and higher mutation rates lead to more diversity and more CNA accumulation. The box plots show the median (centre), 1st (lower hinge), and 3rd (upper hinge) quartiles of the data; the whiskers extend to 1.5× of the interquartile range (distance between the 1st and 3rd quartiles); data beyond the interquartile range are plotted individually.

At gland level (individual clone) resolution the pattern of CNAs was quite similar within cancers (Fig 2B, Table S4-5). In all cases, a ‘core karyotype’ was evident across the glands from that case, with a small numbers of glands showing additional subclonal CNAs (mean CNAs per gland = 7, mean clonal CNAs=6.5). We note that intra-tumour CNA heterogeneity was detected, but we emphasise the clonality of most CNAs. Zooming in further, single cell analysis by FISH in four additional cases (Fig S6) and single cell sequencing in case C274 (reported in Bollen *et al*.^24^) confirmed ongoing instability at the level of individual cancer cells within glands.

Single nucleotide variants (SNVs) were called from individual gland exome sequencing data and were used to reconstruct phylogenetic trees (Methods; Fig 2C, Table S6). Trees were balanced (ancestral nodes have equal numbers of offspring; Yule Model; p>0.05 for all cases) suggestive of a lack of stringent subclonal positive selection, in accordance with previous reports^20,21^. Individual glands from MSS tumours contained on average 156 SNVs (range: 68-232) which equated to 40 more SNVs than the most recent common ancestor (MRCA) cell of the cancer (371 SNVs for hypermutant cases). We calculated the SNV accumulation rate per year using only C>T mutations in the four trinucleotide contexts associated with COSMIC mutational signature 1 (ACG, CCG, GCG, TCG) which are known to accrue at a constant rate in CRC^25^. This analysis produced an estimate of the time to the MRCA of 24 years (range: 11-30 years). Despite this elapsed time, we observed little subclonal accrual of CNAs within the cancers.

### Mathematical modelling negative selection on CNAs

We hypothesised that the distinct pattern of CNAs observed in each CRC represented a (local) optimum in the fitness landscape, with negative (purifying) selection suppressing diversity at gland and bulk levels. To test this hypothesis, we developed a mathematical model of CNA accrual during cancer growth and the evolutionary dynamics due to the action of negative selection on the CNAs. Individuals in the model (‘glands’) had variable fitness 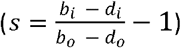 defined by the gland’s karyotype according to the relation:

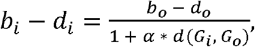

where *b*_*i*_ (*d*_*i*_) was the birth (death) rate of gland *i, G*_*i*_ was the karyotype of gland *i, G*_*o*_ was a pre-specified ‘optimal’ karyotype that maximised fitness, *b*_*o*_ (*d*_*o*_) was the “optimal” birth (death) rate, and the function *d*(*G*_*i*_, *G*_*o*_) measured the genetic distance between the two karyotypes. A gland’s karyotype was represented by a vector of length *C*, where *C* was the number of bins used in our empirical data analysis, with each entry representing the integer copy number of each locus in the gland’s (abstracted) genome. The number of CNAs followed Poisson distribution at mutation rate *µ* per gland division, where the size of each CNA (in the unit of bins) was sampled from the sizes of observed CNAs in our data. A gland’s fitness was then inversely proportional to the absolute distance between a gland’s karyotype and the maximum fitness karyotype (*G*_*o*_); the parameter *α* scaled the strength of stabilising selection experienced by non-optimal karyotypes, with *α* = 0 defining a “flat” landscape where all karyotypes have the same fitness, *s = s*_*max*_ (Fig 2B).

Stochastic simulations showed that ongoing mutation within a growing tumour in the absence of negative selection (*α* = 0) generated high CNA heterogeneity (Fig 2C&D). A growing tumour that initially occupied the fitness maxima (karyotype of the first cancer gland = *G*_*o*_) experienced negative selection which suppressed CNA diversity, with stronger negative selection (greater *α*) more severely limiting diversity (Fig 2C&D), and higher mutation rates generating more diversity (Fig 2D). Thus, modelling suggested that negative selection for an optimal CNA karyotype was a plausible explanation of the apparent stability of the aneuploid genome observed in CRC glands.

Our previous single cell sequencing data, derived from cells extracted from colorectal glands from the same tumours analysed in this study^24^, provided an empirical measurement of the lower limit of the CNA rate at *µ* ≈ 0.1 new CNAs per gland division. The dynamics of cells within glands affect the scaling of per-cell to per-gland mutation rates in a complex fashion dependent on the mechanism of selection upon cell/gland karyotypes; for simplicity we neglected to consider these dynamics and took prior distributions that set per-gland mutation rates the same as cellular rates. We used Bayesian model selection to compare the fit of models with negative selection versus no selection (purely neutral evolution) on the 6 cancers subjected to gland-by-gland analysis. In 4/6 cancers the model with stabilising selection better represented the data than the model without stabilising selection (Fig 2A), and it was not possible to distinguish between the models in 1/6 cases. The model rests on a number of assumptions, and the goodness of these assumptions (see Discussion) determines the strength of the conclusion that can be drawn from these analyses. Nevertheless, modelling provided some quantitative evidence that the low levels of CNAs diversity in primary CRCs was frequently due to the acting of stabilising selection upon a core karyotype.

### Ongoing CNA accrural in CRC model systems

Our model of negative selection presumes that CNA events were constantly generated during tumour development and were purged from the population by negative selection, leading to minimal CNA heterogeneity. An alternative hypothesis is that CNA alteration rates are very low most of the time, possibly with occasional bursts of high CNA rates that produced the observed highly-altered genomes. We note this is not necessarily the same as *punctuated evolution*, which does not require bursts of mutation rates because phenotypes develop in peripherally isolated populations in separated environmental niches^26^. High ongoing CNA rates in cancer and specifically in colorectal malignancies, rather than a burst of CAN events, has been demonstrated previously^24^, and was supported by single cell assays in our dataset (Fig S6 and ref ^24^).

We sought to provide further evidence of an ongoing high CNA rate in CRC. We generated single cell DNA sequencing data from SW620 colon cancer cells (Fig S7A, Table S7). Using 91 high quality cells (see Methods) we performed CNA calling and phylogenetic analyses. This highlighted that individual cells had ongoing chromosomal instability because the cells had generated multiple new CNAs since their MRCA, in addition to a number of shared (clonal) CNAs in the population (Fig 3A). The new CNAs were spread across the genome, and phylogenetic analysis showed that cell-unique (tip) CNAs were enriched for chromosome 11 losses. This may indicate that the copy number gain state (CN = 3) in chromosome 11 is unstable and frequently reversed in SW620 cells, or could indicate selection for this loss in our cell culture conditions.

**Figure 3:**
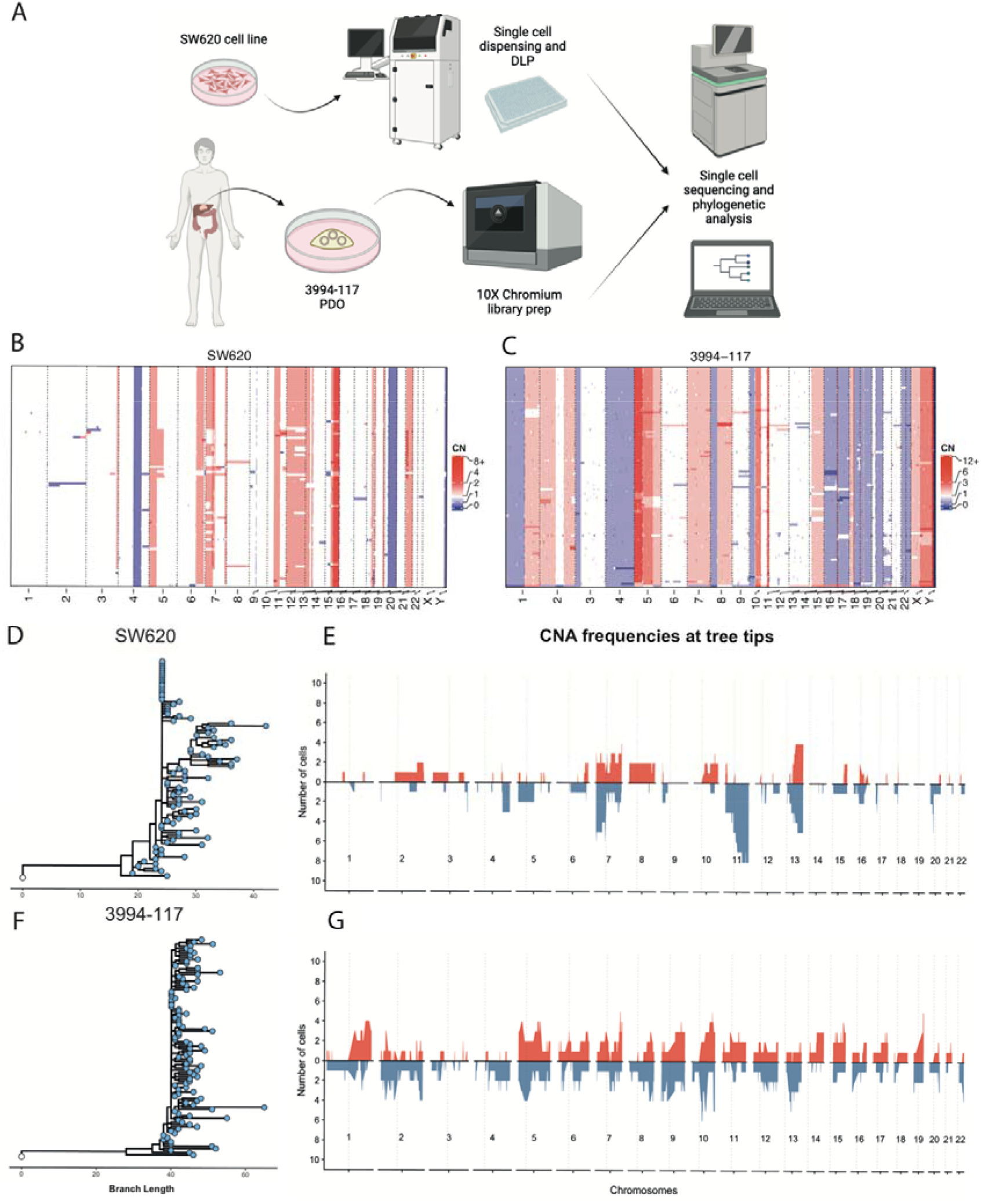
Ongoing copy number alterations in model systems. **(A)** Schematic of the method used to obtain and sequence single-cells from cultured colorectal cell-line SW620 and patient derived organoid 3994-1117. Cell line SW620 underwent single-cell dispensing using the cellenONE system, followed by direct library prep (DLP) while the 10x Genomics Chromium was used to prepare single cell libraries from the patient derived organoid S994-1117. **(B)** Heatmap showing copy number alterations (CNAs) in single-cells from colorectal cell-line SW620 and **(C)** from patient derived organoid 3994-1117. Red=gain, blue=loss. **(D)** Phylogenetic tree constructed using CNAs in the CRC cell line SW620. **(E)** Frequency of copy number gains and losses at the tips of the SW620 phylogenetic tree as in 3G. **(F)** Phylogenetic tree constructed using CNAs in the CRC organoid 3994-117. **(G)** Frequency of copy number gains and losses at the tips of the 3994-117 phylogenetic tree.

We next performed single cell DNA sequencing in a patient-derived colorectal cancer organoid^27^, which also showed ongoing CNA generation in individual cells that were spread across the entire genome (Fig 3C-D, Fig S7B, Table S7).

### Negative selection on CNAs continues through metastatic dissemination and treatment

We then investigated how CNAs evolved during metastatic spread and treatment. Colonisation of a new tissue and the application of therapy represent new selective pressures that could change the core karyotype observed in primary CRCs. Even in the absence of positive selection, bottlenecks during metastatic colonisation combined with the strong effects of genetic drift in an initial small population, could lead to gross differences between primary tumour and metastatic deposits (i.e. founder effect). CRC frequently metastasises to the liver and less frequently to other organs, including the lung. We collected tissue from 23 primary sites and 68 metastatic lesions from 22 patients, with the material collected longitudinally from a mode of 2 time-points (range 1-5 time points). Five patients remained untreated throughout their studied time course (an additional three patients received no treatment between sampled timepoints), while other patients had experienced a diverse range of treatment regimens (Table S8). Using sWGS, we measured inferred absolute copy number of CNAs for each tumour deposit (total of 168 distinct regions analysed) and called GD as described above. Comparing metastatic lesions to our previous primary CRC cohort (Fig 1), we observed that CNA diversity only fractionally increased in the metastatic cohort (Fig S8A). The frequency of GD was significantly higher in metastatic cases (18/22 [82%] had GD) versus regular CRCs (16/81 [20%] cases GD; p<0.0001; Fig S8B), and was comparable to previous reports of GD in metastatic CRC^28^. GD was clonal in two-thirds of metastatic cases with GD (12/18; 66%).

We examined how CNAs evolved over space and time by comparing patterns of CNAs within and between metastatic lesions and to the primary tumour. Additionally, we performed phylogenetic analysis per patient in the cohort based on CNAs using MEDICC2^29^ (Fig S9). We observed several examples of the maintenance of a core abnormal karyotype in the primary tumour wherein the pattern of CNAs did not greatly change in metastatic lesions through space, time, metastasis to other tissues and through chemotherapy and/or targeted treatment (Fig 3A-D, Fig S10-12, Table S9). In nearly all cases there were some CNA differences between lesions in the same patient, and we do not discount that these differences may be important for tumour cell biology.

We measured the proportion of the aberrant genome that was subclonal in pairwise sample comparisons within each patient, the mean of which we term divergence, in addition to counting phylogenetic events. Patient LM0027 displayed little divergence in karyotype between the primary tumour and a metastasis taken a year later in the liver, with no treatment in-between (Fig 3A; divergence of 0.26 [0.18-0.31], mean absolute PGA difference 5.4 percentage points, mean number of phylogenetic events 11 [4-17]). In patient LM0071, despite 23 months between the primary colon and metastatic liver samples 3-months of oxaliplatin and capecitabine chemotherapy, maintenance of a core karyotype was observed (Fig 3A; divergence 0.17 [0.03-0.22], mean absolute PGA difference 1.6pp, mean number of events 5.1 [2-7]). LM0029 was followed for 28 months from primary diagnosis, through hepatic metastatectomies, one pulmonary metastatectomy, and three challenges of chemotherapy in combination with anti-epidermal growth factor monoclonal antibody cetuximab. Again, the core karyotype was broadly maintained throughout, with the largest difference seen in the lung metastases which lacked a gain of chromosome 2 (Fig 4A, divergence between consecutive sampling points of 0.19 [0.13-0.25], 0.34 [0.19-0.60], 0.26 [0.12-0.52], 0.38 respectively, mean absolute PGA differences 4.6pp, 6.1pp, 6.2pp and 18pp respectively, mean number of events, 33 [30-36], 27 [14-38], 25 [13-40], 25 [25-25], respectively). Similarly, in patient LM0139, resection of the primary CRC at baseline was followed by hepatic metastatectomy at 23 months and pulmonary metastatectomy at 72 months. Despite 6 years of elapsed evolution and metastasis to two separate organs, the final karyotype sampled in the lung had a very similar pattern of CNAs to the initial karyotype observed in the primary, demonstrating marked stability across time and space (divergence between first and last timepoint 0.28 [0.18-0.38], mean absolute PGA difference 12pp, mean number of events 36 [35-36]; Fig S13A).

**Figure 4:**
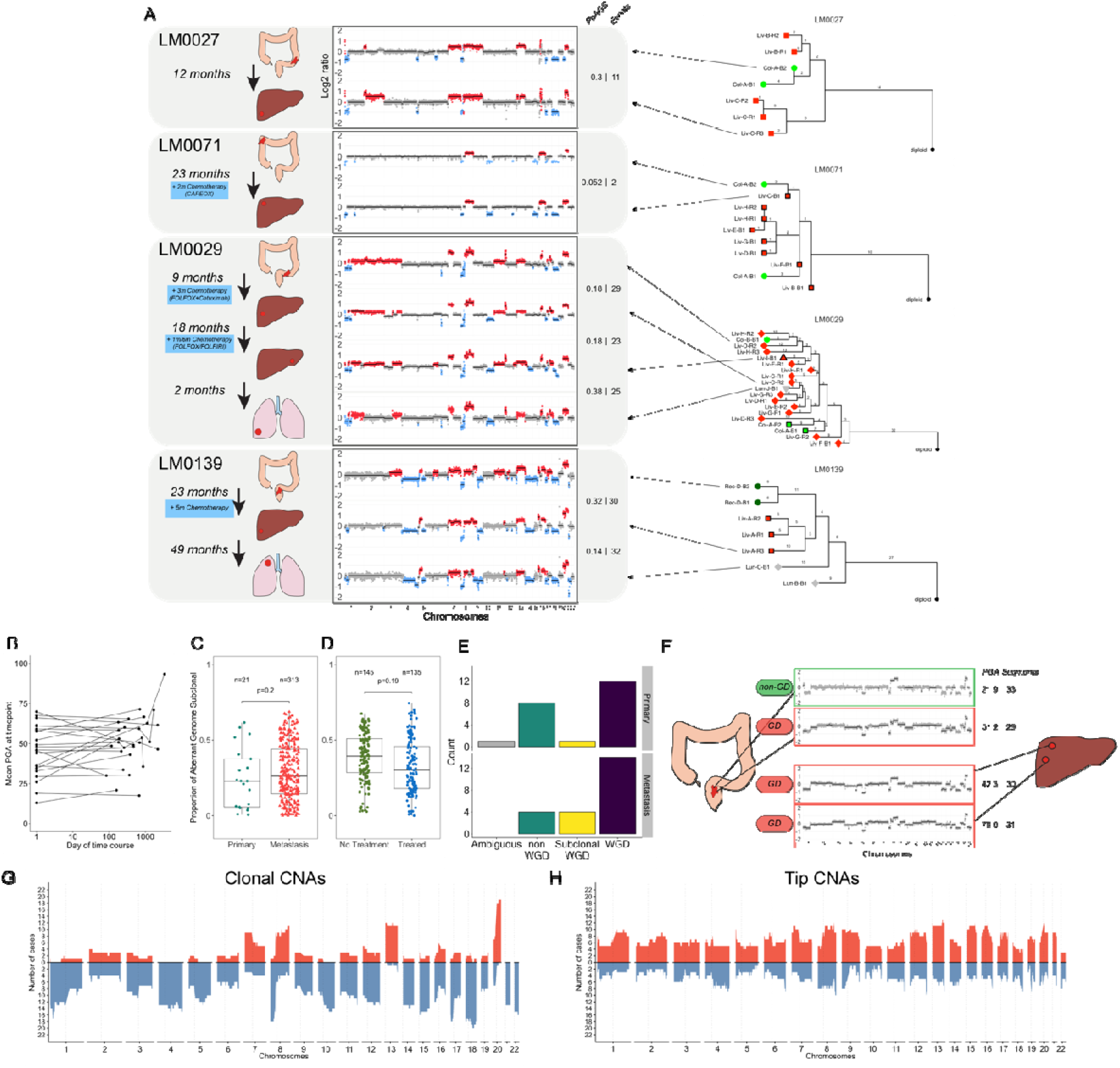
Stability of CNAs across space, time, tissue and treatment in colorectal cancer. **(A)** Longitudinal assessment of CNAs in primary colorectal cancers and metachronous metastatic lesions in liver and lung showed similar CNA profiles were observed over space time, tissue and through various treatment courses. Timelines of four representative metastatic patients are shown. Schematics show organ where cancer deposit was sampled. Genome-wide view of CNAs with gains (red) and losses (blue) relative to baseline ploidy. Proportion of Aberrant Genome Subclonal (PoAGS) and number of events are displayed for each comparison presented sequentially. To the right of each timeline is the corresponding phylogenetic tree as described in Fig S9. **(B)** Mean percent genome altered (PGA) averaged across all lesions at the sampling time by time since baseline (removal of the primary CRC). The PGA remained broadly constant over time in each patient (x-axis is days and is on a log scale). **(C)** The pairwise difference in CNAs between regions of the primary tumours was not significantly different to the pairwise difference between regions of paired metastatic lesions. **(D)** Across timepoint genetic divergence was not significantly different after treatment with various chemotherapy and/or targeted agents. **(E)** GD was common in metastatic primary tumours and metastatic deposits and was sometimes present in some but not all of the metastatic lesions from a patient (called subclonal GD). **(F)** An example case (LM0032) where GD was subclonal in the primary but clonal in the metastasis. The greater burden of CNAs is evident in the GD samples. **(G)** Frequency of copy number gains (red) and losses (losses) present on the clonal branches in the set of phylogenetic trees created from the cohort. Over-representation of some CNAs is evident. **(H)** Frequency of copy number gains and losses at the tips of the phylogenetic trees (i.e. unique events). CNAs are uniformly spread across the genome.

Despite these examples of relative stability, we observed some cases where CNA patterns diverged markedly between lesions. For example, the metastatic lesions in the ovary and omentum sampled at the second timepoint in LM0018, taken only 7 months after the sampling of the primary, showed the highest between timepoint divergence in the cohort (divergence 0.62 [0.55-0.7], mean absolute PGA difference 27pp, mean number of events 42 [32-52], Fig S10). The patient received FOLFOX in between sampling points. These samples were our only examples of ovary and omentum spread but raise the possibility that CNAs may change substantially to enable colonisation of these organs. We also highlight case LM0011 which also showed a large increase in divergence between the two timepoints in the colon and liver taken 16 months apart despite receiving no treatment in the interval (divergence 0.53 [0.49-0.62], mean absolute PGA difference 21pp, mean number of events 52 [40-67], Fig S10).

Alterations in the proportion of genome altered (PGA) were generally low in sequential timepoints indicating aneuploidy stability (mean PGA increased by 3.4pp on average for consecutive timepoints in the cohort; Fig 4B). Within the same time point, CNA differences between metastatic regions was not significantly different from CNA differences between primary regions (Fig 4C; p=0.2, linear mixed model, Fig S13B) and divergence within regions of primary colorectal samples was consistent with our analysis of regular CRCs (divergence 0.24). Overall, CNA divergence was not significantly different in on-treatment intervals versus off-treatment intervals (p=0.19, linear mixed model, Fig 3D).

Subclonal GD was inferred in six patients (LM0018, LM0019, LM0029, LM0032, LM0056, LM0124) and in these cases GD was enriched in metastatic samples versus primaries (Fig 4E). For example, the metastatic samples in LM0032 were GD whereas one of the two primary samples was not, suggesting GD was subclonal at the primary site and potentially led to the metastatic clone (Fig 4F). In LM0056 and LM0124 all primary samples were classified as non-GD, yet all bar one metastasis sample in each patient was considered GD, indicating the GD likely originated after metastasis. Only a single liver metastasis region was called as GD in LM0019. In primary CRCs, GD tumours had less diverse CNAs. The reason for this difference could be related to elapsed time: in GD primaries could be fit and grow rapidly with little elapsed time for new CNA evolution despite an intrinsically higher mutation rate, whereas in metastasis sufficient time elapses for new CNAs to accrue.

Based on the phylogenetic structure inferred for all 22 patients, we separated CNA events into three categories: clonal (events preceding the MRCA of the tumour samples), intermediate (events shared between samples but not preceding the MRCA) and tip (events unique to each sample). Clonal CNAs had occurred early in the cancer’s development, and so selection had had time to operate on these events. In comparison, tip CNAs had occurred more recently and so may not yet have been assorted by selection. The distribution of clonal events across the genome showed some CNAs were far more common than others consistent with the typical pattern for CRC^30^. Events such as 8q, 13 and 20 gain, as well as 8p and 17p loss were enriched only in clonal alterations (Fig 4G). Contrastingly, the events at the tips of the phylogenetic trees (Fig 4H) showed a strikingly uniform distribution across the genome suggesting an ongoing accrual of CNAs at a constant rate across the genome, prior to assortment by selection.

## Discussion

Our study reveals the dynamics of CNA during the evolution of colorectal cancer. We observe large changes in CNAs during the progression from benign to malignant disease, but thereafter the cancer karyotype appears quite stable, and typically remains relatively unchanged over many years, through metastasis to distant organs, and despite treatment. Taken together, these data suggested a model whereby strong positive selection for an abnormal karyotype occurs during transformation from benign to malignant disease, and that thereafter the karyotype evolution is constrained by negative selection acting at the level of individual cells including in metastasis and during treatment. This is a distinct mechanism to transient bursts in the CNA rate as has been previously suggested to occur in breast cancer evolution^31^. We note that our finding of pervasive negative selection at CNA level is congruent with our previous observation of “Big Bang” effectively-neutral evolution in CRCs^20,21^, because negative selection acts to remove deleterious variants before they can expand to detectable levels and so only neutrally evolving lineages remain. Positively selected variants, both SNVs and CNAs, are clonal in the tumour and part of the ‘trunk’ of the phylogenetic tree but do not drive subclone expansions.

We propose an altered “core karyotype” is necessary for the initiation of malignancy, and that the same karyotype is also sufficient for subsequent metastatic spread. These evolutionary dynamics are consistent with the observation of early metastasis in CRC^32^, whereby the genome of the initial primary tumour is competent for metastasis. Given the inter-tumour heterogeneity in patterns of CNAs, we suggest that the fitness landscape traversed by CNAs is ‘rugged’, with local fitness peaks located at normal-diploid and at multiple aberrant-genome positions, and that fitness of any individual karyotype is likely to be somewhat patient-specific due to the background of germline and somatic variants and tumour microenvironment composition. There could be multiple local maxima close together in genotype space – or similarly new CNAs could be deleterious or neutral – our analysis is insensitive to detect these dynamics. Our data do suggest that fitness peaks are likely sufficiently distant in karyotype space to prevent easy transition between peaks, and potentially that the selective landscape of CNAs differs between benign and malignant disease.

Given the potentially defining role of CNAs in CRC evolution, we hypothesise that the fixed pattern of CNAs in any individual CRC likely defines the biology of that tumour, and consequently is a major determinant of patient prognosis and treatment response. That the karyotype is relatively fixed through treatment is surprising, as presumably the effectiveness of therapy is dependent on it acting as a novel and strong selective pressure. We had expected this new selective pressure to change the fitness landscape and as a result cause wholesale change in the CNAs observed post-therapy. It may be that the fitness peak of maximal treatment-resistance is too far from that occupied by an untreated tumour and so is highly unlikely to evolve. However, we emphasise that our analysis has taken a genome-wide average view, and adaption could be driven by individual and/or small CNAs^8,33^ for which our analysis is insensitive to detect. Nevertheless, an alternative explanation is that chemotherapy response in CRC may be entirely plastic^34^, and so does not involve the clonal selection of individual lineages – a phenomenon that may be common across cancer types^35^. In CRC, previous work is suggestive that plasticity is established very early in CRC^35^ and is consistent with reports of early metastatic spread^32^.

We reiterate that there are typically clear genetic differences between matched primaries and mets, and we do not rule out that these differences could contribute importantly to tumour biology: we recognise that our analysis is insensitive to detect a driving role for such events. We acknowledge that our study is limited by its principal reliance upon bulk sampling, including our use of individual tumour glands, and stronger evidence for or against negative selection on CNAs will derive from large-scale single cell sequencing analyses. Our modelling relies on a number of critical but untested assumptions, such as simple scaling of mutation rates between tumour cells and glands, steady division rates of disseminated tumour cells, uncertain estimates of the age of metastases, and monoclonal origins of metastases. All these assumptions can alter the expectation of the degree of genetic divergence between and within lesions. Appropriately addressing these assumptions is a major undertaking and should be a priority for future work.

Collectively, out study demonstrates a plausible role for negative selection acting on the karyotype to define the genomes of colorectal cancers.

## Methods

Full methods are provided as supplementary material.

## Supporting information

Supplementary note on modeling of glands

## Acknowledgments

MJ, AS and TG acknowledge funding from Cancer Research UK (A22745, A22909 and A19771 respectively). TG, AS and SL are funded by the Wellcome Trust (202778/Z/16/Z, 202778/B/16/Z and 206314/Z/17/Z respectively) and via a Wellcome Trust award to the Centre for Evolution and Cancer at the ICR (105104/Z/14/Z). SL and LMW acknowledge funding from the National Institute for Health Research (NIHR) Oxford Biomedical Research Centre. DS, AS and TG received funding for this project from the NIH via the Cancer Systems Biology Consortium U54 scheme (CA217376). TG received pilot funding from the Barts Charity. AB was supported by Barts Health NHS Trust and Barts Charity grant. ED is supported by the S:CORT consortium which is funded by a grant from the Medical Research Council and Cancer Research UK. JB is in part funded by the UCLH/UCL Biomedical Research Centre. MJ, MN and MRJ receive funding support from Cancer Research UK UCL Experimental Cancer Medicine Centre and the Department of Health’s National Institute for Health Research Biomedical Research Centres funding scheme (Reference: C12125/ A15576). The research leading to these results has received funding from the People Programme (Marie Curie Actions) of the European Union’s Seventh Framework Programme (FP7/2007-2013) under REA grant agreement n° 608765 (MM). This research was supported by the National Institute for Health Research (NIHR) Biomedical Research Centre based at Guy’s and St Thomas’ NHS Foundation Trust and King’s College London who performed some of the genome sequencing, and we note that the views expressed are those of the author(s) and not necessarily those of the NHS, the NIHR or the Department of Health. We are grateful for the support of the QMUL Genome Centre (Charles Mein) and Institute of Cancer Research Tumour Profiling Unit for support with genomic assays. A small project grant from the UK Bowel and Cancer Research (BACR) Charity to TG and SL pump primed this research. Organ likeness in Fig 3A, 3F and Fig S6, S10-12 were based on images in Servier Medical Art (licensed under a Creative Commons Attribution 3.0 Unported License).

## Conflicts of interest statement

TG and AMB are named as coinventors on a patent application that describe a method for TCR sequencing (GB2305655.9), and TG is named on a method to measure evolutionary dynamics in cancers using DNA methylation (GB2317139.0). TG has received an honorarium from Genentech.

## Notes

### Summary of Updates

The revision includes additional analyses that help strengthen our conclusions. Primarily (1) improved copy number calling for the metastasis cohort that takes into account different ploidy solutions, (2) a whole genome duplication (WGD) classifier to identify samples in low coverage whole genome sequencing data that have experienced a WGD event, (3) the inclusion of single cell DNA sequencing (scDNA-seq) from two colorectal cancer models to investigate single cell level diversity, (4) phylogenetic analysis of both longitudinal metastasis samples and scDNA-seq to time copy number alterations and (5) improved computational modelling of copy number instability with direct parameter inference per patient.

