## Supplementary Figures for "Negative selection may cause grossly altered but broadly stable karyotypes in metastatic colorectal cancer"

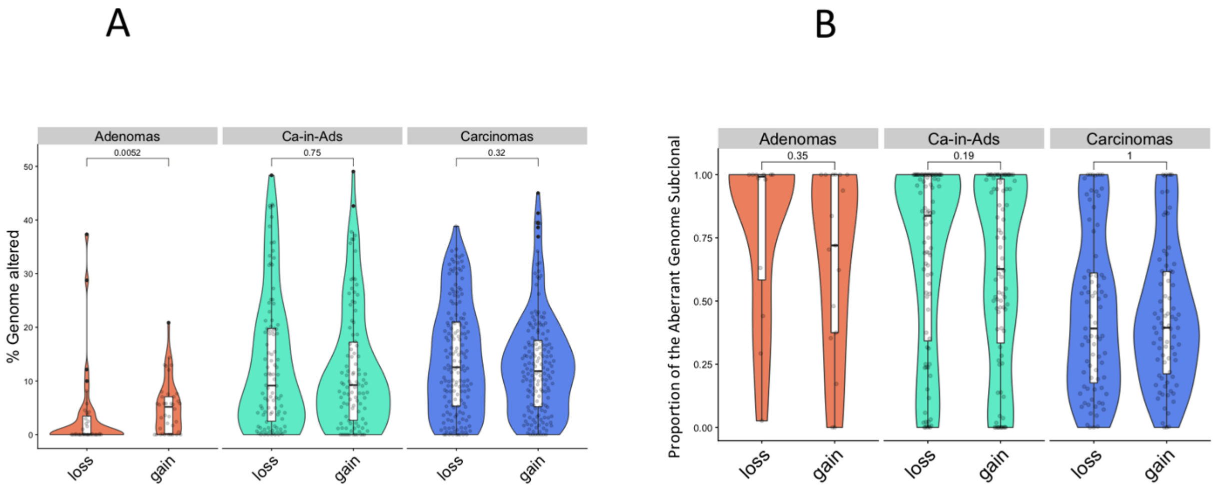


**Supplemental Figure 1: CNA subclonality decreases with progression within early colorectal cancers. (A)** % of the genome with losses and gains in CRAs (orange), ca-in-ads (green) and CRCs (blue). Gains were significantly more common than losses in CRAs, but gains and losses were equally common in more advanced lesions. **(B)** The proportion of CNAs that were subclonal was similar for losses and gains across tumour stages.

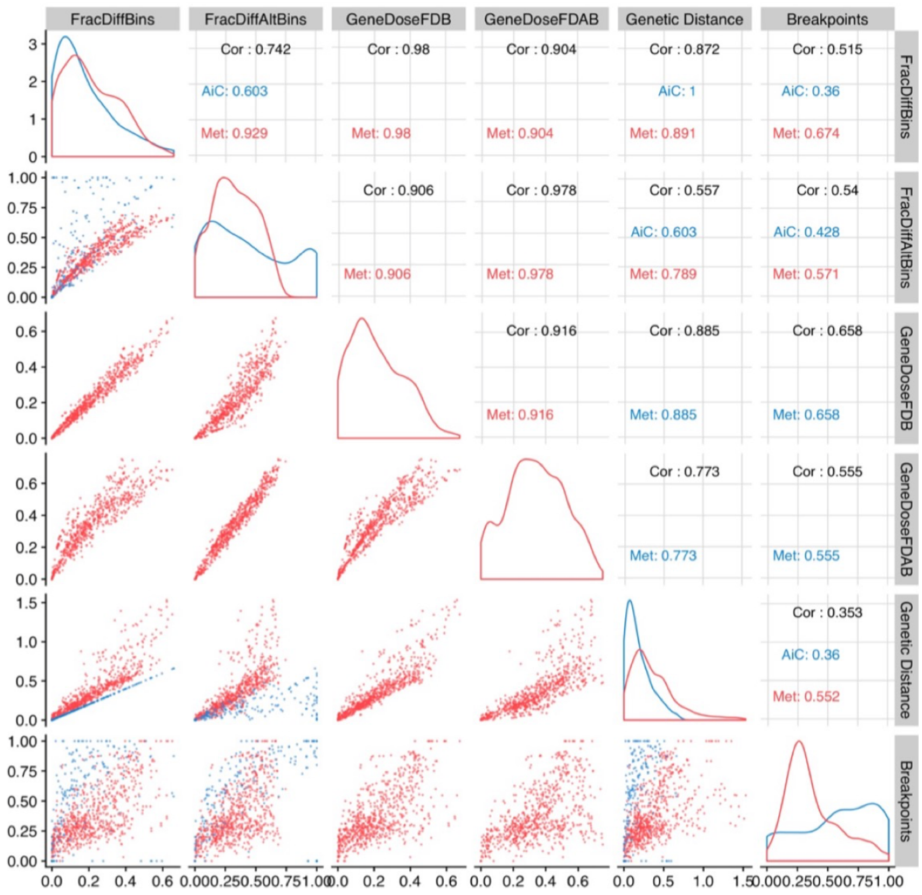

**Supplemental Figure 2: CNA subclonality assessed by different metrics**. We computed 6 different statistics to describe the burden of subclonal CNAs in pairwise comparison between samples. Statistics are defined in the supplementary methods in section “Assessment of CNA divergence”. We observed a high degree of correlation between all statistics examined.

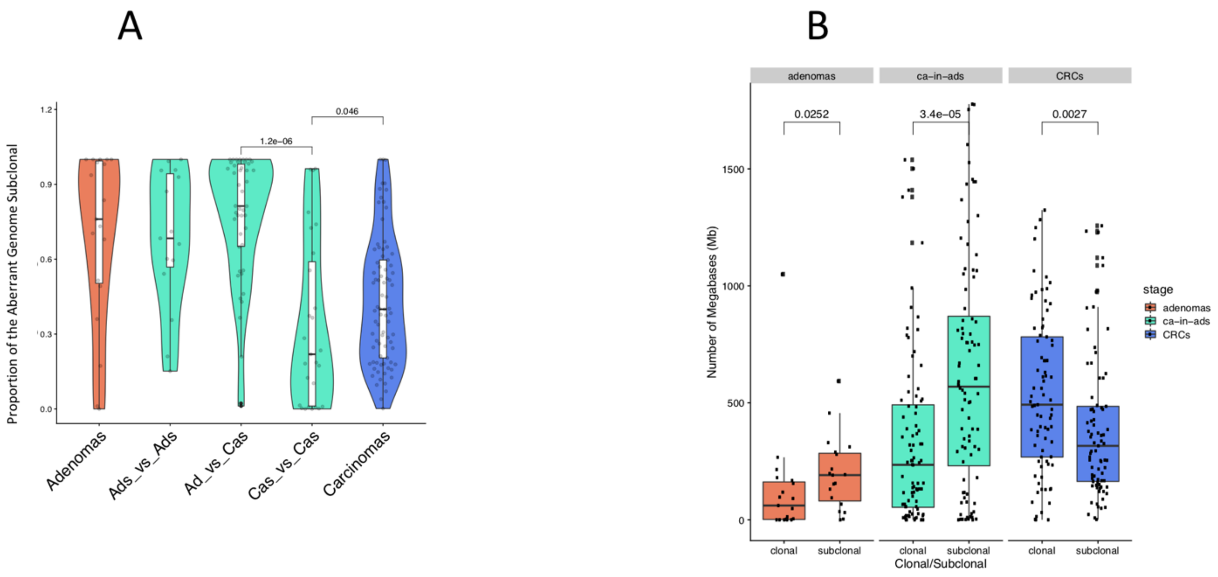


**Supplemental Figure 3: Divergence decreases with progression within early colorectal.** (A) Proportion of genome that is divergent across different lesion and sampling types; CRAs (orange), early cancers (green) and CRCs (blue). Ads vs Ads signifies comparison between adenomatous regions within early cancers. Ads vs Cas compares adenoma vs cancer components in early cancers. Cas vs Cas compares cancer components of early cancers. (B) The number of megabases (Mb) within the genome that were either subclonal or clonal within the CRAs (orange), ca-in-ads (green) and CRCs (blue). (C) The number of divergent Mbs within CRAs (orange), ca-in-ads (green) and CRCs (blue).

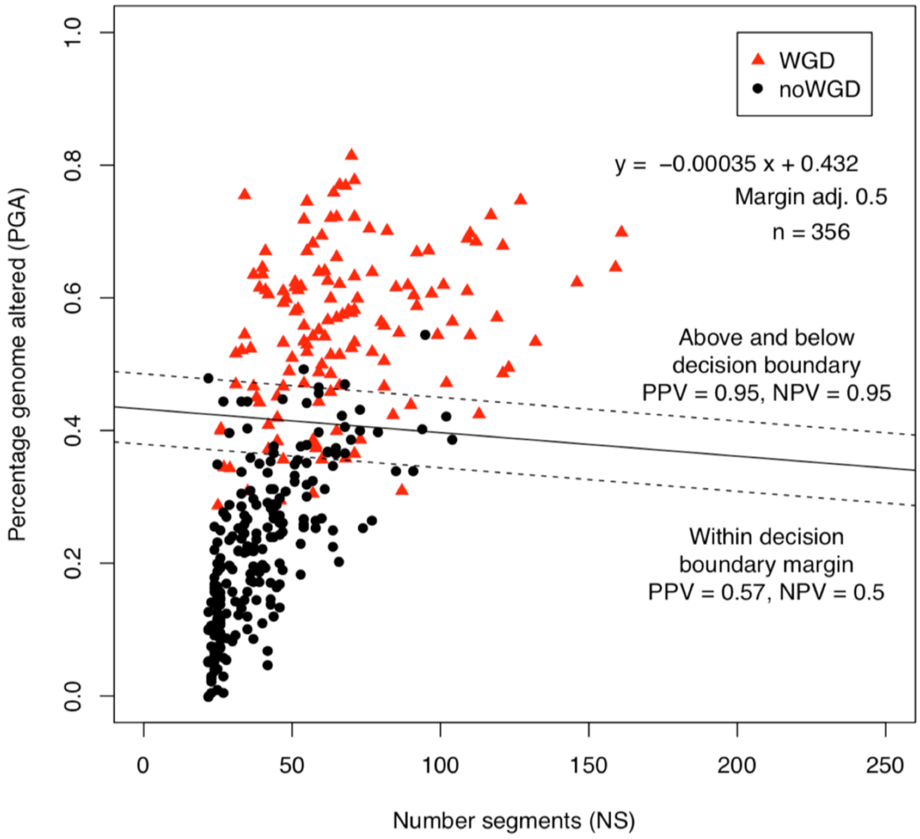


**Supplemental Figure 4: Derivation of a simple genome doubling classifier.** Genome doubling calls from a subset of PCAWG patients (red triangles= GD, black circles= not GD) by number of breakpoints and percentage genome with non-baseline copy number in each sample. A linear classifier was fitted to the data (solid line) and dashed lines demarcate the “uncertain region” where samples are called approximately equally as GD and non-GD.

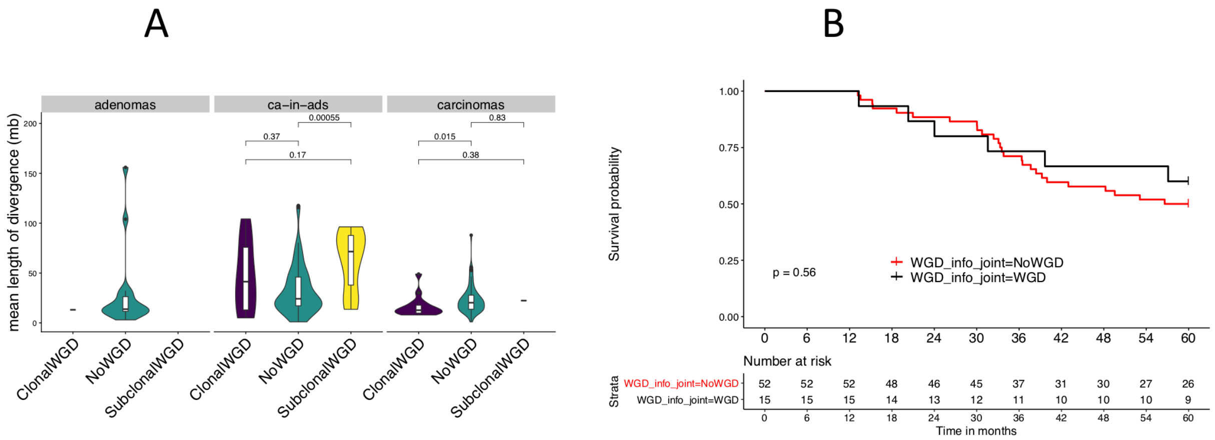


**Supplemental Figure 5: Divergence is highest in subclonal and least in clonal genome doubled cases**. (A) The length of divergent segments was calculated within clonal, subclonal and non-genome doubled samples in CRAs, early CRCs (ca-in-ads) and CRCs. The mean divergent length of segments was found to be greatest in subclonal and least in clonal genome doubled samples. (B) Early CRCs that were non-genome doubled or which had clonal genome doubling had a low proportion of subclonal CNAs. (C) A Kaplan Meier of CRCs comparing the survival of patients with CRC that were either genome doubled or not genome doubled showed no significance difference in survival between the two groups.

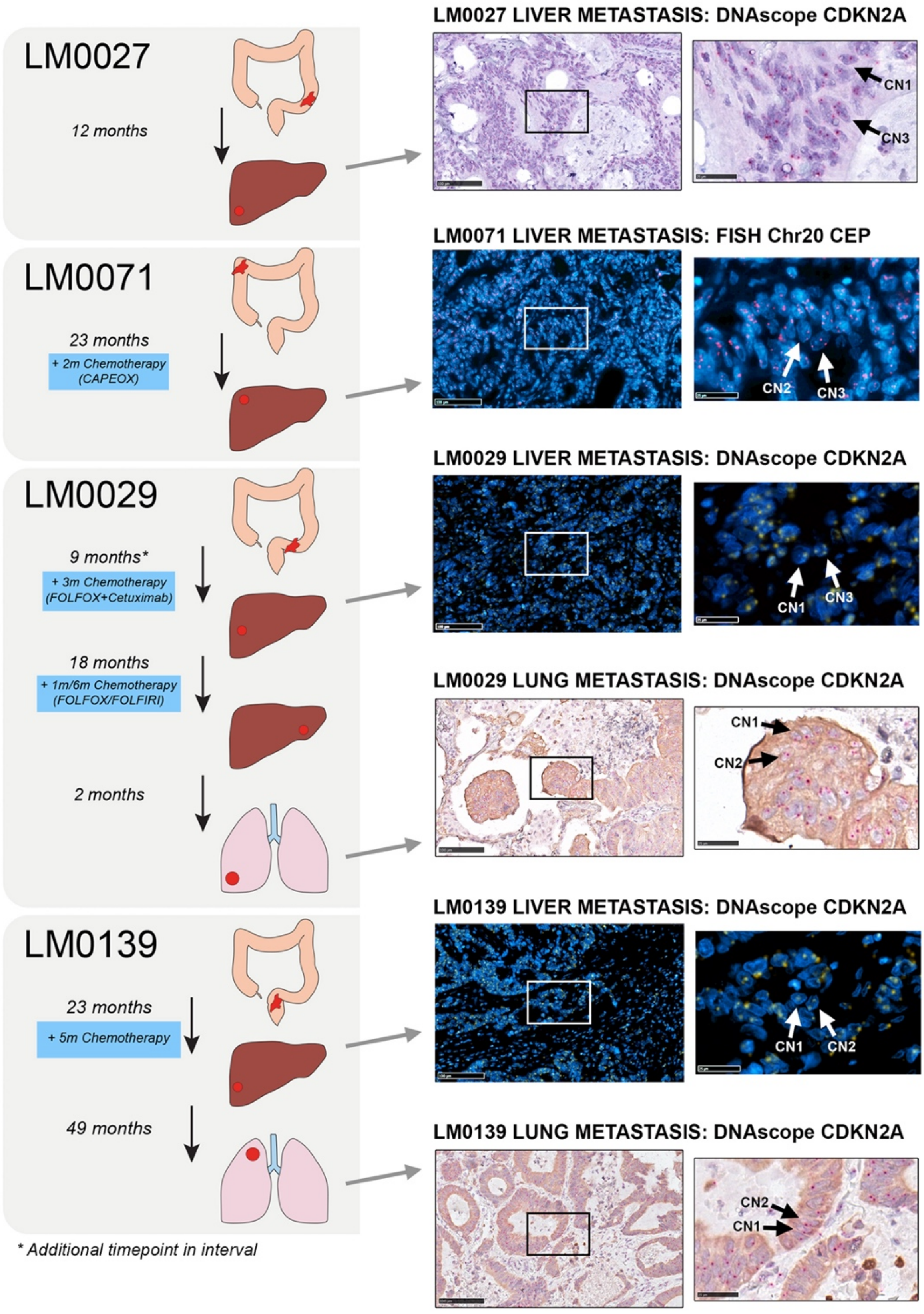


**Supplemental Figure 6:** ***In situ* analysis using FISH and DNAscope shows copy number instability at the single cell level.** DNAscope probes against CDKN2A or FISH Chr20 chromosome enumeration probe (CEP) were applied to FFPE sections from 4 metastatic CRC cases, and nuclei were counterstained with DAPI (fluorescent images) or haematoxylin (brightfield images). Small punctate dots indicate probe binding, and cells with changes in copy number (CN) at the locus of interest are indicated with arrows. Representative images corresponding to the four patients described in main figure 3 are shown.

**Supplemental Figure 7: Cross-cohort study shows CNA subclonality, CNA burden and genome doubling change with progression.** (A) The proportion of subclonal CNAs versus percentage genome altered stratified by lesion type with adjusted p (FDR) values reported in comparisons. Significantly different correlations between lesion types were detected between lesion types. CNA subclonality decreased with PGA in adenomas and CRCs but increased in metastases and early cancers (B) The proportions of samples that were clonal, subclonal and non-genome doubled across our early-primary-metastatic cohort within adenomas, early CRCs, CRCs, and metastatic CRC cohorts. Genome doubling increased with progression of disease, from a single adenoma sample that is genome doubled, to the highest proportion of samples as clonal genome doubled within the metastatic cohort.

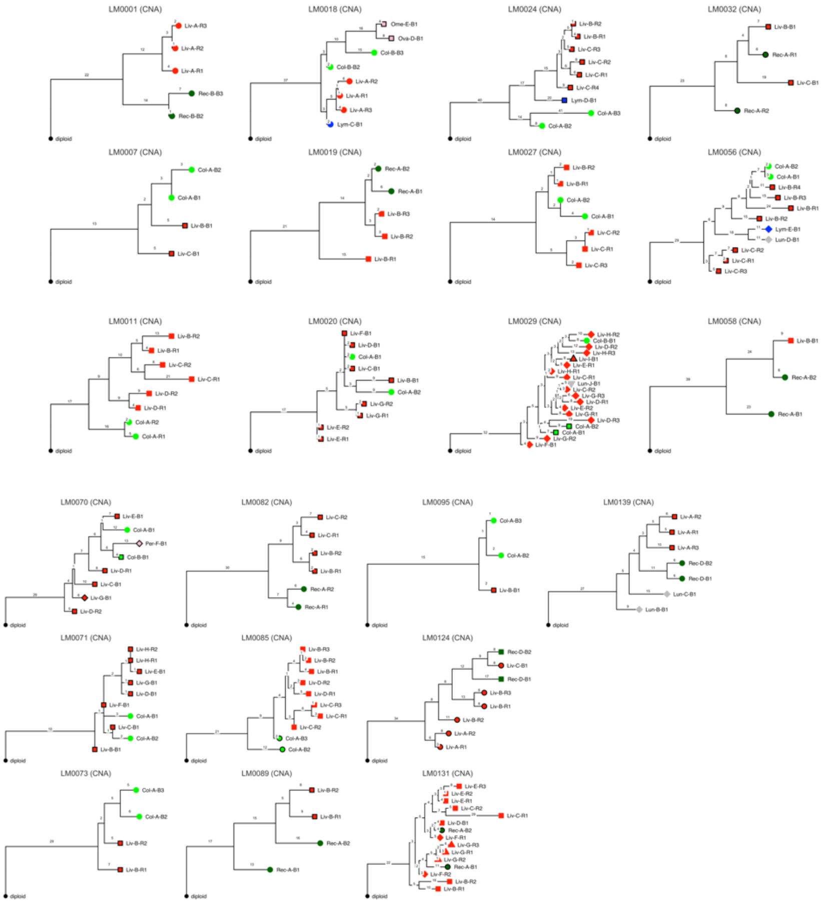


**Supplemental Figure 8: Multi-site longitudinal karyotype phylogenetic assessment of individual CRC patients during tumour progression and drug treatment *(patients LM001 – LM0029).*** Phylogenetic trees derived from karyotypes of individual metastatic CRC patients**.** Tree annotated with the number of copy number events and rooted at a diploid normal. Individual leaves of the tree annotated by organ location: Col denotes colon, Rec denotes rectum, Liv denotes liver, Lym denotes lymph node, Ova denotes ovary, Ome denotes omentum, Per denotes Peritoneum and Lun denotes lung.

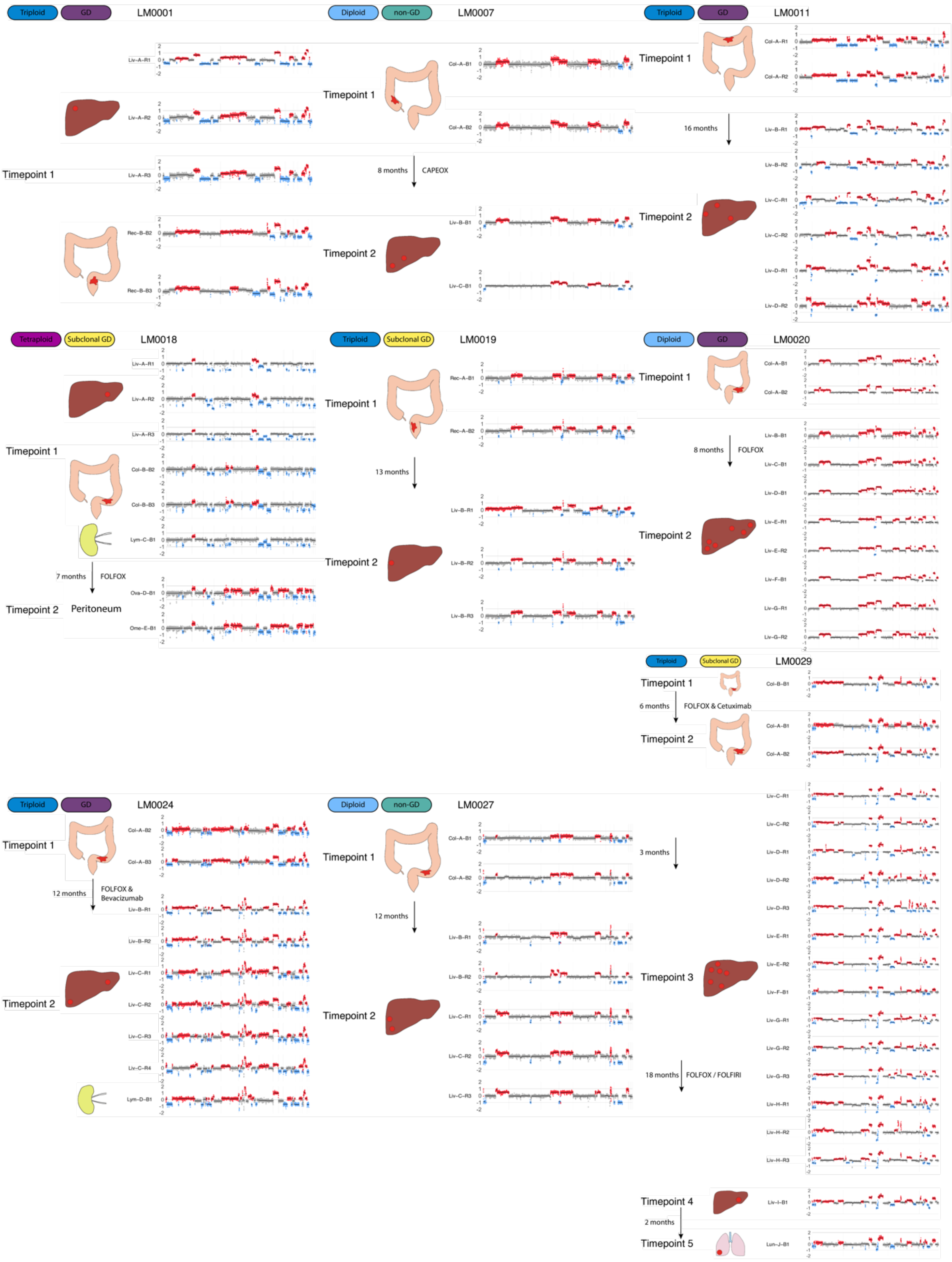


**Supplemental Figure 9: Multi-site longitudinal karyotype assessment of individual CRC patients during tumour progression and drug treatment *(patients LM001 – LM0029).*** Timelines for each patient are annotated with the timepoint at which they were sampled, which organ was sampled and the approximate location in which the samples were taken from. Intervals are annotated with the elapsing period in months and include information on the treatment received in the interval (absence of an annotation indicates no treatment). Copy number alteration plots represent gains in red (greater than baseline) and losses in blue (less than baseline) with baseline copy number (ploidy of sample) in grey. Log2ratios per genomic bin is represented on the y-axis and bins are ordered by chromosome and genomic location on the x-axis. For each patient there is an annotation of the ploidy called for the patient based on a ploidy search performed for absolute copy number calling as well as the classification of the patient's genome duplication status (GD) as determined separately by a GD classifier. Organs can be recognised from linked samples; Col denotes colon, Rec denotes rectum, Liv denotes liver, Lym denotes lymph node, Ova denotes ovary, Ome denotes omentum, Per denotes Peritoneum and Lun denotes lung.

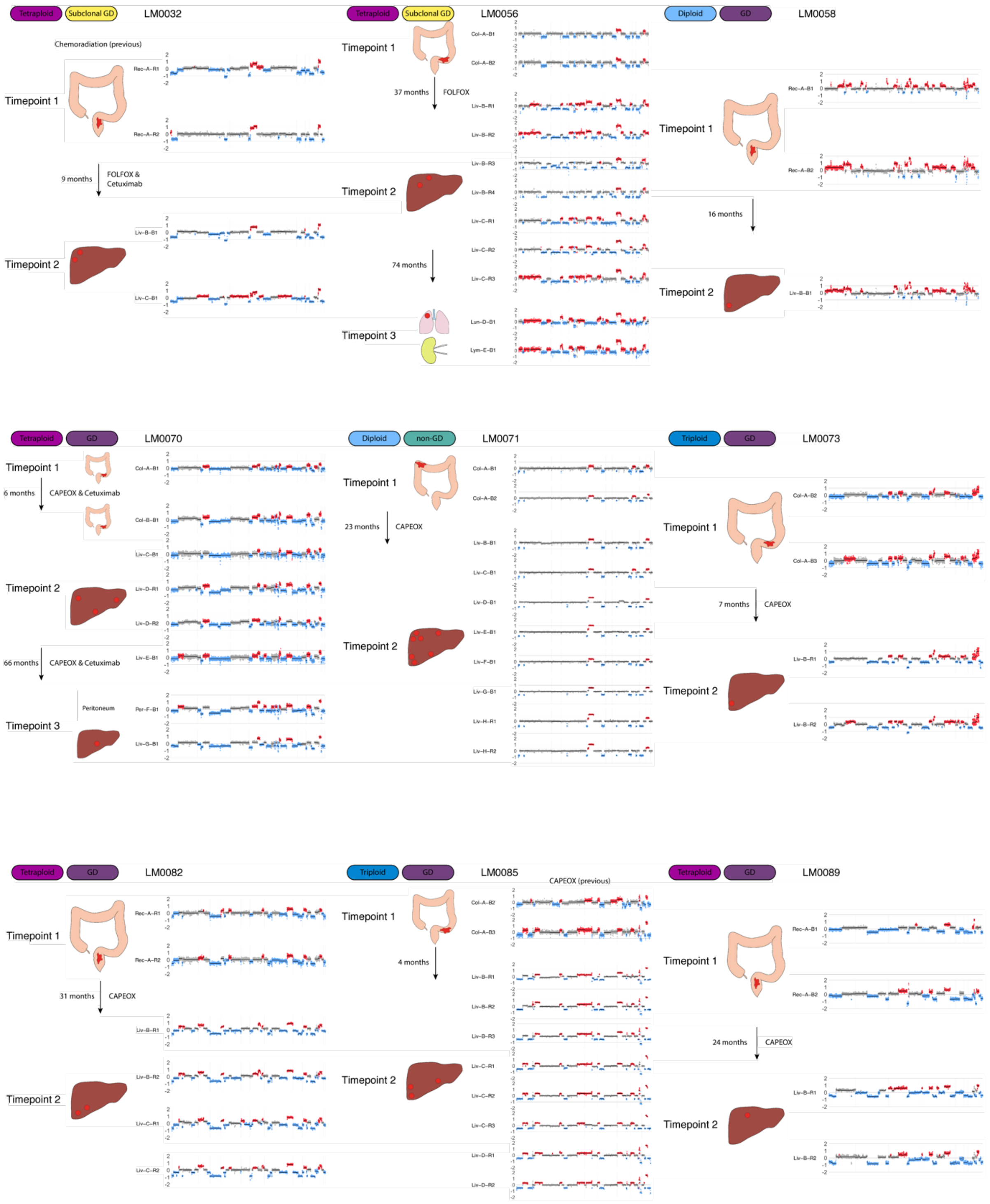


**Supplemental Figure 10: Multi-site longitudinal karyotype assessment of individual CRC patients during tumour progression and drug treatment *(patients LM0032 – LM0089).*** Timeline schematics are present as in Supplemental Figure 10. Of note, LM0032 received chemoradiation prior to the first timepoint and LM0085 received CAPEOX prior to the first timepoint.

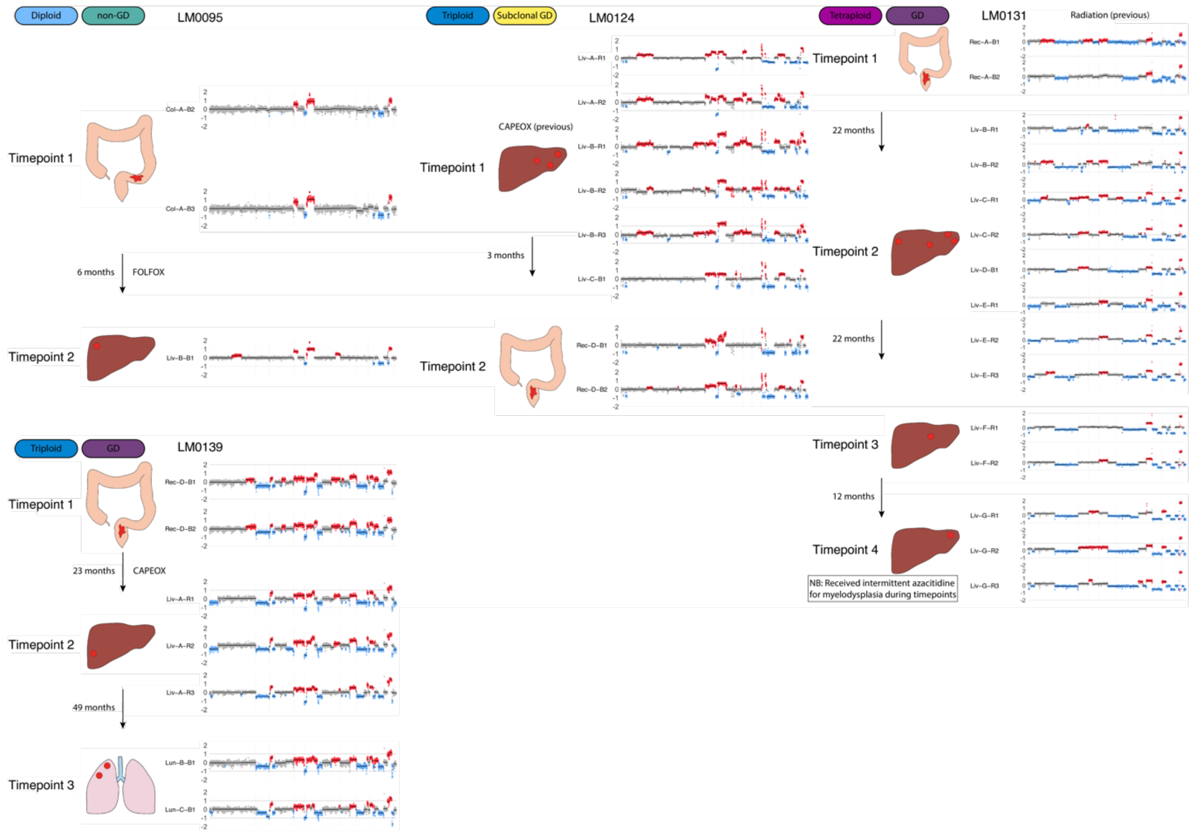


**Supplemental Figure 11: Multi-site longitudinal karyotype assessment of individual CRC patients during tumour progression and drug treatment *(patients LM0095 – LM0139).*** Timeline schematics are present as in Supplemental Figure 10. Of note, LM0131 received radiation prior to timepoint 1 and intermittently received azacytidine for myelodysplasia during the timepoints. LM0124 received CAPEOX prior to timepoint 1.

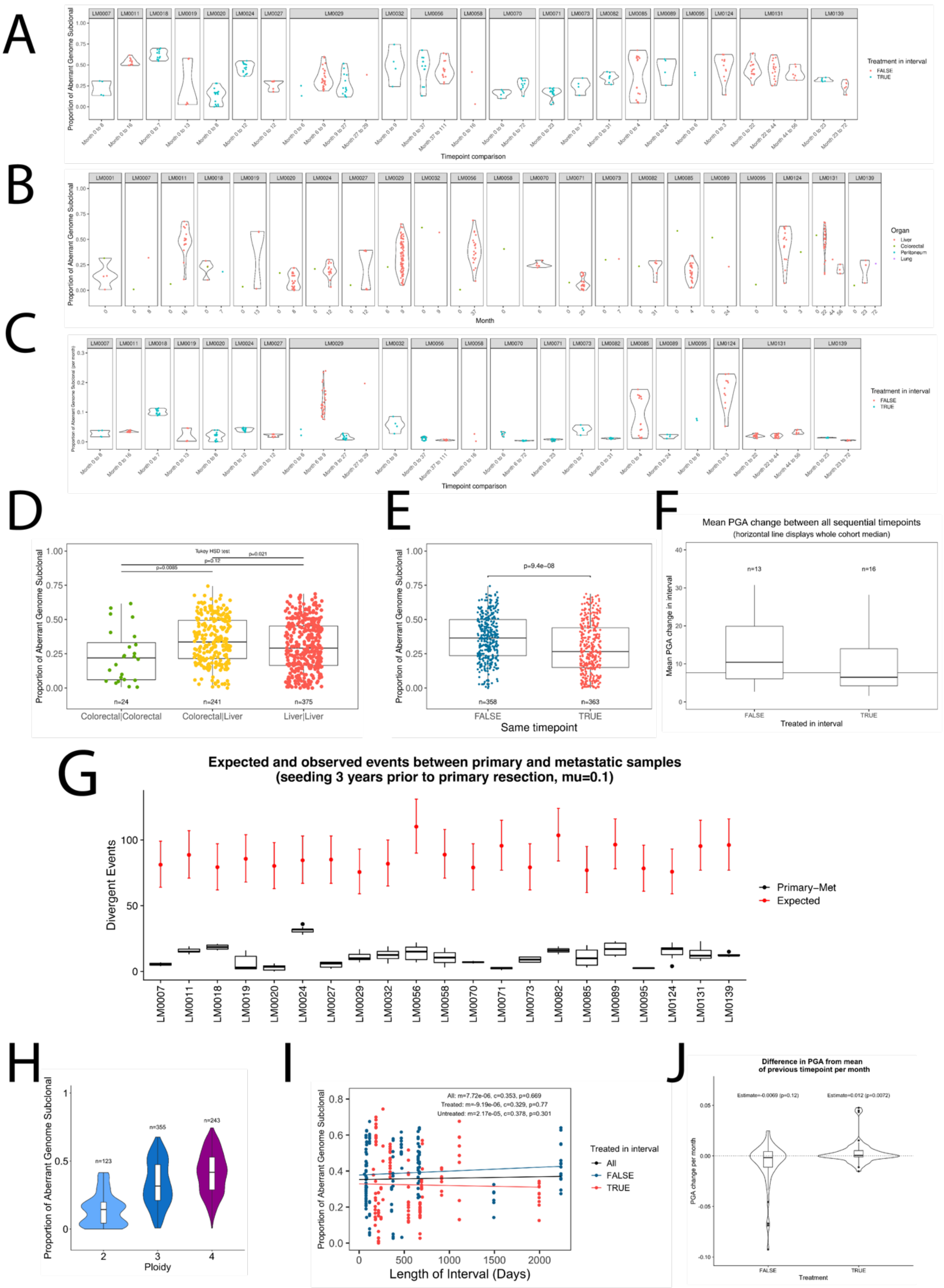


**Supplemental Figure 12: Patterns of aneuploidy evolution during metastatic spread and treatment.** (A) Proportion of aberrant genome subclonal for sample pairs in sequential timepoint comparisons for all patients in the longitudinal metastasis cohort including annotation on if treatment was received in the interval. (B) Proportion of aberrant genome subclonal for sample pairs at the same timepoint and organ type. (C) Rate of change in proportion of aberrant genome subclonal per month for sequential timepoints. (D) Comparison of proportion of aberrant genome subclonal for within the colon or rectum, within the liver and between colon or rectum and liver, diversity is significantly higher between organs (Tukey HSD test). (E) Proportion of aberrant genome subclonal for comparison in the same time point and between time points. Diversity is higher between timepoints (mixed effects linear model). (F) Mean absolute PGA change in all sequential intervals categorised by treatment status, horizontal line represents the cohort median. (G) Number of pairwise events between primary tumour samples and first sampled metastasis samples (Primary-Met) are less than the expected range of events (red) across six years of cell divisions (three years of separate divergence) plus the time difference between the sampling of the primary and the metastasis given the expectation of an event once every ten cell divisions and a cell division every 3 days according to a Poisson distribution. (H) Proportion of aberrant genome subclonal pairwise comparison for samples according to their patient tumour ploidy. (I) Comparison of proportion of aberrant genome subclonal and time elapsed between sampling points shows the relationship to be flat for all data and both treated and untreated samples in the previous interval (linear models). (J) Mean PGA change per month for treated and untreated intervals was not significantly different from zero in untreated intervals but slightly greater than zero in treated intervals (0.012, mixed effects linear model).

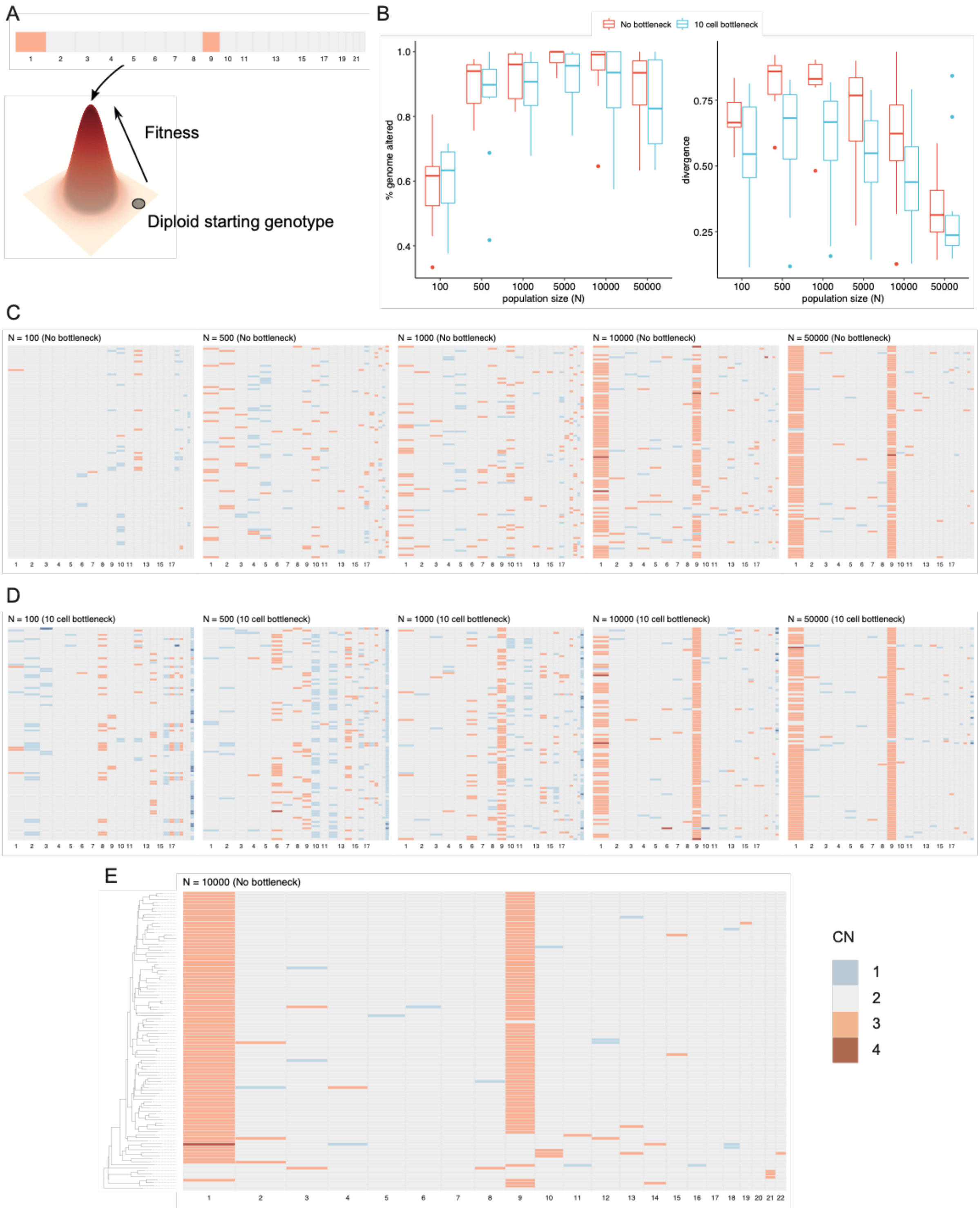


**Supplementary Figure 13: Simulations demonstrate stabilising selection suppresses karyotypic diversity when tumours are close to fitness peak. (A)** Tumours were simulated with glands starting with a diploid karyotype and the approximated C319 karyotype (gain of chromosome 1 and 9) considered as optimum. **(B)** Karyotype divergence (measured by percentage genome altered or average pairwise divergence) initially increased as the population grew before decreasing once most glands had a CNA pattern locating them at the fitness peak (10 simulations obtained with CNA rate 0.05, selective strength 1, initial death rate 0.1, final population size 50000, and 100 sampled glands). This was also true when a 10-cell bottleneck was introduced to model the seeding of a metastasis. The box plots show the median (centre), 1st (lower hinge), and 3rd (upper hinge) quartiles of the data; the whiskers extend to 1.5× of the interquartile range (distance between the 1st and 3rd quartiles); data beyond the interquartile range are plotted individually. Heatmaps of sampled glands in one simulation at increased population size demonstrate that glands gradually arrive at fitness peak, either without a bottleneck **(C)** or with a bottleneck **(D)**. **(E)** The phylogenetic tree of sampled glands in another simulation suggests that multiple lineages converge on the optimal karyotype rather than the selective sweep of a single lineage.

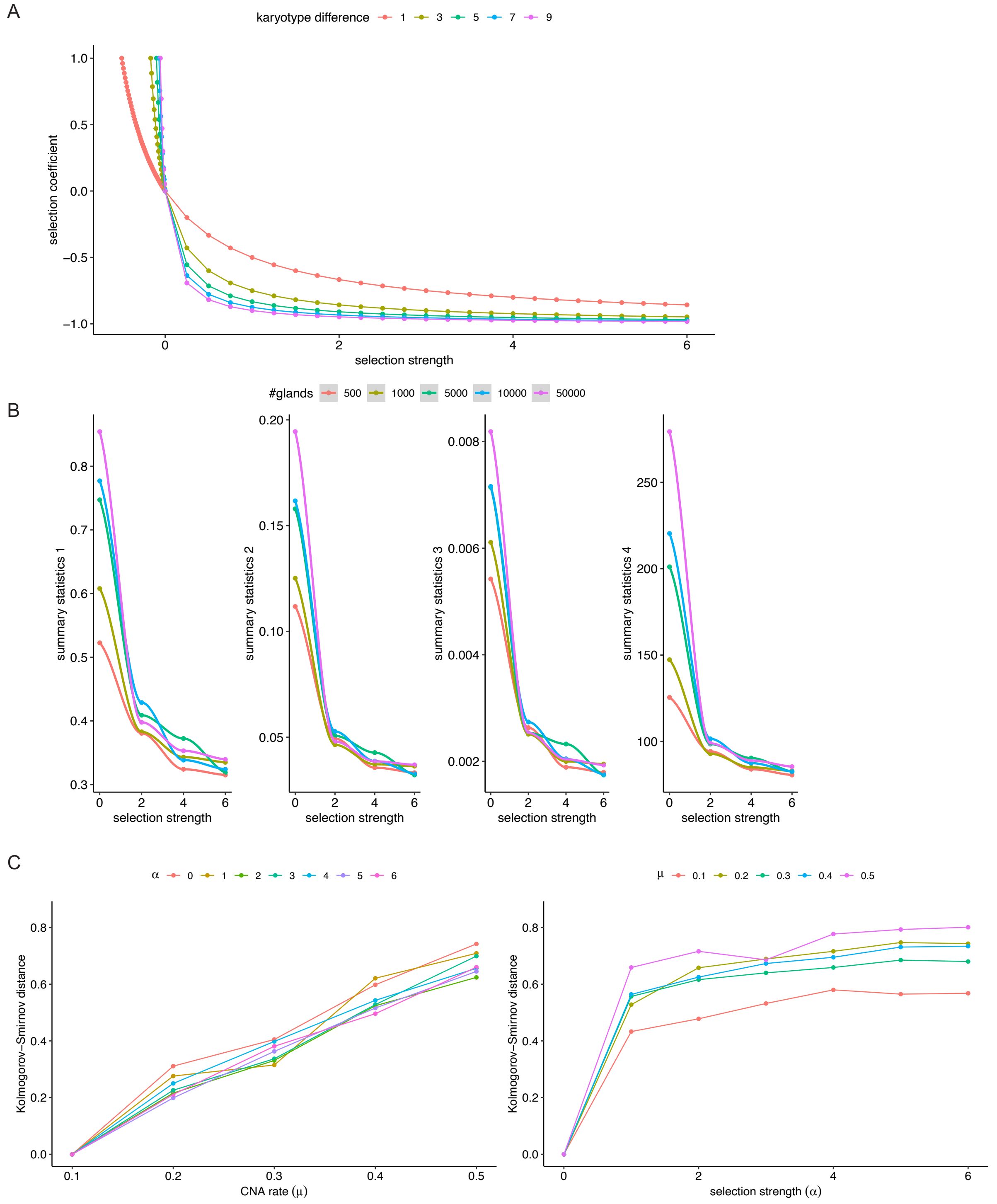


**Supplemental Figure 13: The non-identifiability of the mutation rate (μ) and the strength of stabilising selection (α) in the stochastic branching process model. (A)** The relationship among selection strength $\alpha$, selection coefficient $s$, and karyotype difference $d(G_i, G_o)$. **(B)** Changes of summary statistics when the simulation stops at different number of glands under different values of selection strength $\alpha$ (based on data of C282, $\mu = 0.2$). Each point is the mean value over 1000 simulations. **(C)** Sensitivity analysis on the power of inferring $\mu$ and $\alpha$. Y-axis shows the Kolmogorov-Smirnov distances between the distributions of the number of mutations in 1000 simulated glands under different parameter settings and the distribution at $\mu = 0.1$ and $\alpha = 0$.
