## Supplementary Methods for "Negative selection may cause grossly altered but broadly stable karyotypes in metastatic colorectal cancer"

### Patient sample acquisition and processing

*1. Archival samples.*

Various cohorts of archival specimens were used in this study:

a) Oxford: CRAs, CRCs and ca-in-ad specimens used in Figure 1 were obtained from the pathology archives of the John Radcliffe hospital under ethical approval from Oxfordshire REC A (MREC 10/H0604/72).
b) Utrecht: 26 early cancer lesions used in Figure 1 were obtained from the University Medical centre in Utrecht. These samples were extracted from an earlier cohort-nested matched case–control study on pedunculated T1 colorectal cancers conducted in the T1 CRC registration cohort, initiated by the Dutch T1 CRC Working Group, a collaboration of Dutch hospitals, including patients with T1 CRC diagnosed between January 1, 2000 and December 31, 2014, identified using the Netherlands Cancer Registry. The earlier study was approved by the Medical Ethics Review Committee of the University Medical Center Utrecht (approval for data collection and histologic review, reference number: 15-487 and 15-716), and performed in accordance with the Helsinki Declaration.
c) Barts: Metastatic CRC samples used in Figure 3 were obtained from the Barts Cancer Tissue Bank (www.cancertissuebank.org; REC Ref: 14/LO/2031; Tissue request reference number 2015/2/QM/TG/CaCOL). After screening the records of 200 patients in the CTB database, 22 patients were chosen who had all undergone surgical resection of the primary CRC and also had liver metastases resected. Subsequently, some of these patients also underwent resection of extra-hepatic metastases (e.g. lung) or recurrent liver metastases. All relevant clinical information associated with these patients was collected by performing a thorough search of the NHS electronic patient records and the CTB database.

*2. Fresh-frozen samples*

Fresh-frozen CRC samples used in Figure 2 were collected from UCL Hospitals under ethical approval 11/LO/1613 and via the UCLH Biobank (15/YH/0311). All samples were collected from patients who had given informed consent. Cancers were obtained from surgical resections and processed within the same day. A maximum of four different spatial regions (biopsies) were sampled from each tumour, and these were further separated into smaller tissue pieces and subsequently cryopreserved (medium: MEM, 10% DMSO, 5% fetal calf serum, 25µM HEPES buffer). Individual glands were pulled from thawed tissue pieces using a stereoscopic microscope (Zeiss Stemi SVII microscope) and transferred and stored in buffer ATL (Qiagen, Hilden, Germany) until DNA extraction.

### Tissue processing, DNA extraction and quality control

*1. Archival samples*

For cases used in Figure 1: FFPE blocks were sectioned onto glass slides at 10µm thickness and following dewaxing, regions of interest were scraped into tissue lysis buffer using a pathologist annotated H&E as a guide.

For cases used in Figure 3: FFPE blocks were sectioned at 8µm thickness onto PALM (Zeiss, Oberkochen, Germany) slides A serial H&E slide was reviewed by one of the collaborating pathologists to confirm tumour content and guide subsequent isolation of cancer tissue. A combination of techniques was used to obtain tumour tissue from the slides: when tumour cell content was high material was scraped into tissue lysis buffer (most primary CRCs had high tumour cell content). When tumour cellularity was intermediate, the PALM MicroBeam laser capture microdissection (LCMD) system (Zeiss) was used to isolate regions enriched for tumour cells, and these regions were gently lifted off the slide and placed in lysis buffer, using a sterile needle, as necessary. Finally, if the tumour cellularity was very poor, LCMD of individual tumour cells was performed.

DNA was extracted using the High Pure FFPET DNA isolation kit (Roche), and quantified using the Invitrogen Qubit Fluorometer before genome-wide copy number analysis using whole genome sequencing or SNP array (details below).

*2. Fresh frozen samples*
Individual tumour glands were lysed in buffer ATL supplemented with proteinase K (Qiagen) during overnight incubation at 56°C. Genomic DNA was extracted using the QiaAmp DNA Micro Kit (Qiagen) following the manufacturer's protocol and further concentrated using Agencourt Ampure XP Beads (Beckman Coulter Inc., Brea, California, USA). Subsequently, DNA concentrations were fluorometrically determined with the Qubit dsDNA HS Assay Kit (Thermo Fisher). Additionally, peripheral blood buffy coat DNA was prepared using the same strategy and used for germline comparison in downstream genomic analyses.

During the study we became aware that isolated tissue we believed to be tumour glands during the microdissection often had very low or no tumour content when measured by genome sequencing, and so may have been normal colon crypts, or other glandular shaped tissue (perhaps stromal in origin) from the tumour. In order to preselect tumour glands for subsequent next generation sequencing (NGS), TaqMan SNP Genotyping (Applied Biosystems) assays were designed for the specific *APC* or *BRAF* mutations previously identified by bulk exome sequencing of the tumour (see below for details). The Taqman assays quantified the allelic fraction of somatic mutations in the *APC* or *BRAF* gene, and so served as a sensitive, quantitative measure of tumour cell content. All putative tumour gland DNAs from a single patient were collectively subjected to quantitative PCR using the Quantstudio 7 system (Applied Biosystems) and matched DNA from bulk tumour tissue was used as positive control. Only glands with detectable mutant allele signal were taken forward to NGS library preparation and sequencing.

**Library preparation and DNA Sequencing**
*1. Sequencing of DNA derived from archival material*
Prior to library preparation, the input DNA was subjected to repair using the NEBNext® FFPE DNA Repair Mix (New England Biolabs, Ipswich, Massachusetts, USA). Library preparation for all samples was done using the NEBNext® Ultra™ II FS DNA Library Prep Kit for Illumina (New England Biolabs). A fragmentation time of 5 minutes was used for the samples. The size selection protocol using Agencourt AMPure® XP beads (Beckman Coulter) was applied only for samples with DNA input ≥100ng going into the FFPE DNA repair step.

All libraries were indexed with unique indexing primers. Two sets of indices were used for the samples: NEBNext® Multiplex Oligos for Illumina® Dual Index Primers Set I and NEBNext® Multiplex Oligos for Illumina® 96 Unique Dual Index Primer Pairs (New England Biolabs). PCR enrichment of adaptor-ligated DNA was done using the above-mentioned indices and the NEBNext® Ultra II Q5 Master Mix (New England Biolabs). The standard protocol, as recommended by the manufacturer, was used for the entire library preparation with one variation across all samples- half volumes were used throughout the process.

The DNA libraries generated were first quantified using the Qubit® dsDNA HS (high sensitivity) Assay kit and the Qubit® 4 Fluorometer (Invitrogen) before running it on the Agilent Tapestation system using an Agilent High Sensitivity D5000 ScreenTape (Agilent Technologies, Santa Clara, California, USA). Subsequently, equimolar pooling and sequencing was done in preparation for shallow whole genome sequencing (sWGS). The pooled libraries were then subjected to sequencing (target genomic coverage 0.1x) either on a NextSeq 550 system or a NovaSeq 6000 system (Illumina).

*2. Sequencing of individual tumour glands*
For the construction of sWGS libraries, DNA obtained from tumour glands was processed using the NEBNext Ultra II DNA Library Prep Kit (New England Biolabs) according to manufacturer's instructions.

WES libraries were prepared from bulk tissue pieces or individual glands with the Agilent SureSelect XT low input reagents in combination with the SureSelect Human All Exon V6 baits (Agilent) and unique dual indexing strategy. WES libraries were prepared from bulk tissue pieces using the Agilent SureSelect whole exome reagents with the SureSelect Human All Exon V5 baits by Source Biosciences (Nottingham, UK) or by Nextera rapid capture whole exome preparation (Illumina, San Diego, California, USA), again according to manufacturer's instructions.

The concentrations of all libraries were quantified using Qubit® dsDNA HS Assay kit and the Qubit 4 Fluorometer (Invitrogen). Library fragment size profiling was performed using the Agilent Tapestation system using Agilent High Sensitivity D1000 ScreenTapes (Agilent Technologies) and subsequently all libraries were pooled equimolarly.

sWGS libraries were subjected to shallow sequencing on a Illumina NextSeq 550 system using mid- or high-output kits (target genomic coverage: 0.1x) while WES libraries were subjected to deep sequencing on a NovaSeq 6000 system using a S2 PE100 flow cell (target exonic coverage: 300x) or a HiSeq 2500 platform (target exonic coverage 50X).

**SNP arrays on DNA derived from archival tissue**
The quality of FFPE DNA extractions was verified using the Infinium HD FFPE QC kit (Illumina), then 100-250ng of each sample was treated using the Infinium HD FFPE Restore kit (Illumina) and processed using the Infinium HTS Assay, Manual Protocol (Illumina). Samples were then run on the HumanOmniExpress-24 v1.1 BeadChip using the iScan system at the QMUL Genome Centre.

**Fluorescence *in situ* hybridisation (FISH)**
Sections of FFPE tissue at 5-micron thickness were stained as previously described[1] using SureFISH probes against Chr8 CEP (Agilent cat. no. G101034R-8) or Chr18 CEP (Agilent cat no G101074R-8).

**Organoid culture and passaging**
3994-117 patient-derived organoid (PDO) was established from a liver metastasis of a colorectal tumour and it has been previously characterised as described in (doi: 10.1126/science.aao2774). PDOs were cultured embedded in Growth Factor Reduced (GFR) Basement Membrane Matrix (Corning), and Advanced DMEM/F12 media (Thermo Fisher Scientific), supplemented with 1X B27 and 1X N2 supplements (Thermo Fisher Scientific), 0.01% BSA (Roche), 2mM L-Glutamine (Thermo Fisher Scientific) and 100 units/ml penicillin streptomycin (Thermo Fisher Scientific). Additionally, 12 different growth factors were used

to maintain PDOs culture: 50 ng/ml EGF, 100 ng/ml Noggin, 500 ng/ml R-Spondin 1, 10 ng/ml FGF-basic, 10 ng/ml FGF-10 (all from PeproTech), 10 nM Gastrin, 10 µM Y-27632, 4 mM Nicotinamide, 5 µM SB202190 (all from Sigma-Aldrich), 100 ng/ml Wnt-3A (R&D Systems), 1 µM Prostaglandin E2 and 0.5 µM A83-01 (Tocris Bioscience).

Passaging of PDOs was performed using TrypLE 1X diluted in 1mM PBS-EDTA (Thermo Fisher Scientific). In short, after media removal, PDOs in matrigel were harvested by pipetting with 1ml of TryplE1X and they were incubated for 20 min at 37oC, with mechanical homogenisation every 5 minutes. Then, PDOs were centrifuged at 1,200 rpm for 5 min at 4oC and washed with HBSS (Thermo Fisher Scientific). Counting and viability measurements were done using 0.4% Trypan Blue staining solution (Thermo Fisher Scientific) and the Countess 3 Automated Cell Counter (Thermo Fisher Scientific). Expected cells were pelleted again and re-seeded in matrigel.

**Monolayer cell culture**

The colorectal cancer cell line SW620 was obtained from the American Type Culture Collection (ATCC CCL-227™) and was cultured in high glucose Dulbecco's modified Eagle's medium (DMEM) with 10% Foetal Bovine Serum (FBS) and 2% Penicillin-Streptomycin. Cells were grown in T-175 vented flasks at 37°C, 5% CO2 and 95% relative humidity. Cell line identities were confirmed using STR analysis at the beginning of the experiment and were tested regularly for mycoplasma infection.

**Single cell WGS using Direct Library Preparation + (DLP+) method.**

For interrogation of SW620 single cell whole genomes using the DLP+ method, the recently published protocol by Laks et al. (Cell 2019) was adapted and optimised for a 384-well "one-pot" plate-based workflow.

First, 1ul each of 384 unique i5 indexing primer at a concentration of 4uM were dispensed into empty 384-well plates and air-dried. 1ul lysis buffer, consisting of 86.2% DirectPCR Lysis Reagent (Cell) (Viagen, cat# 301-C), 8.6% protease (Qiagen, cat# 19157), 5.2% glycerol (Sigma, cat# G5516-1L) was added to each well and a single SW620 cell was sorted into each well using the CellenOne F1.4 system (Cellenion). Following centrifugation of the plate at 3000G for 5 min and overnight incubation at +4°C cell lysis was performed by incubating the plate at 50°C for 1h and 70°C for 15min. Tagmentation of genomic DNA was carried out by adding 1.8ul tagmentation mix, consisting of 1.775ul tagmentation buffer (20mM Tris-HCl pH 8, 10mM $MgCl_2$, 20% dimethylformamide), 0.0165ul Tween, 0.00875ul Tn5 loaded (Diagenode, cat# C01070012-30), and incubating at 55°C for 10min. Subsequently, the reactions were neutralised by adding 1ul neutralisation mix (0.5ul Qiagen protease, 0.01ul Tween and 0.049ul ddH2O) and incubating the plate at 50°C for 15min and 70°C for 10min. Finally, PCR amplification was performed after adding 3.8ul master mix (3.78ul 2x NEB Ultra II Q5 master mix (NEB, cat# M0544L), 1mM i7 indexing primer (500nM final, IDT)) with the following cycling parameters: Gap-filling at 72°C for 5min, initial denaturation at 98°C for 30s followed by 10 cycles of 98°C for 30s and 65°C for 75s and final elongation at 65°C for 5 min. PCR products were pooled and purified using a Zymo DCC5 spin column and then subjected to Exonuclease I digestion (NEB cat# M0568) and 1x Ampure bead purification. Final libraries were quantified using Agilent HS D1000 screentapes on an Agilent Tapestation 4200 system and sequenced on the Illumina Novaseq 6000 system using Novaseq S2 PE50 flow cells.

**Single cell WGS using 10X Chromium Single Cell CNV**

Prior to starting the 10X workflow the cell viability was confirmed to be > 90% using the Countess™ II Automated Cell (Thermo Fisher Scientific), targeting an estimated number of captured cells ranging between 1,500-3,000. The single cell suspension was loaded on a Chromium Single Cell 3′ Chip C, followed by a Chip D (10X Genomics). Single-cell gel bead-in-emulsion were generated using the Chromium Single Cell DNA kit and the Chromium Controller.

A single-cell DNA-seq library was prepared, and the final library quality was confirmed using the Tapestation High Sensitivity Screentape assay D1000 (Agilent) and a High Sensitivity Qubit dsDNA Kit (Life Technologies). Samples were normalised, pooled, and sequenced in an Illumina NovaSeq 6000 according to standard 10X Genomics recommendations at a median depth of at least 750K read pairs per cell. The sequencing depth enables over 2Mb CNV detection per cell.

**Bioinformatic processing**

*SNP array processing*

Log-R values were calculated from the HumanOmniExpress-24 v1.1 BeadChip. For a subset of the samples matched normals were used to create a panel of normals (PON). Following this, the PON was used to calculate the log-R ratio for each sample. For 500Kb bins across the genome, the mean log-R ratio was calculated and assigned to a single position within the bin to decrease noise within the sample and to enable a direct comparison between the sWGS data (see below). The *winsorize* and *pcf* functions from the copynumber package[2] were then used to remove outliers and to segment the data respectively. For each sample a density plot of the log-R ratios for the segments produced a number of normally distributed peaks, the greatest of which was assumed diploid normal. All values were diploid centered to this point. For those samples which were potentially triploid or tetraploid, the samples were diploid centered to the 'correct' diploid peak for all multi-region samples from the same patient. These new log-R ratio values were then analysed using the CGHcall package[3], which adopts the DNAcopy[4] method for segmentation. Each sample was preprocessed, normalized, segmented, normalized post segmentation, and gains and losses were called.

*Sequencing quality control*

Per base quality, GC content, and adapter contamination, of the raw read sequences in each fastq file were assessed using FastQC (versions 0.11.5 and 0.11.8)[5]. Read clipping in a subset of samples that were found to have adapter contamination was performed using the Skewer tool (versions 0.2.1 and 0.2.2). A minimum read length of 35bp after trimming was required. Reads were aligned with BWA (versions 0.7.15 and 0.7.5 using default parameters) to UCSC human genome reference version 19. Genome build GRCh38 was used for the time course metastasis dataset.

To check the quality of the alignments a combination of Picard[6] tools (*CollectWgsMetrics, CollectInsertSizeMetrics, CollectAlignmentSummaryMetrics,* all versions 2.18 or 2.18.11) was used. Duplicate reads were marked using Picard *MarkDuplicates*. Of note, we ensured that the percentage of reads removed was not excessive (<35%) and that the target coverage and depth were obtained. Finally, on WES data, bamutil *ClipOverlap* version 1.0.14 was run to clip overlapping reads where insert sizes were small.

*Copy number alteration (CNA) calling in shallow whole genome sequencing data*

On sWGS, the R package QDNAseq[7] was used to call CNAs. The QDNAseq package bins reads from the input bam file into a user-specified series of bins, apply filtering (loess residual and blacklisting), correct the bins for GC content and mappability, then normalises read counts per bin using the 'median' method and smooths outlier counts.

QDNAseq was also used to segment per-bin read counts into 'segments' of equal copy number. The 'sqrt' function was used to transform the data, bin counts normalized with the 'normalizeSegmentedBins' function.

To facilitate integration of SNP array data samples with the sWGS, losses and gains for both data types were called using the CGHcall R package *calls* function. For data included in Figures 1 where sWGS and SNP array data were compared, 1Mb bins were constructed for both sWGS and SNP array data, and the data segmented. Segments <=2Mb in length were deemed noise and removed.

For data in Figure 3 500kb bins were used with no segment-size filtering. In Figure 3, the resulting $\log_2$ ratio values in each bin as well as the segment mean of each bin were normalised according to the global median $\log_2$ ratio by subtracting this value from all $\log_2$ ratios.

*Analysis of exome sequencing data: calling small nucleotide variants (SNVs) and insertion and deletions (indels)*
A two-pass approach was used to identify SNVs and indels. First, to obtain a set of candidate variant 'proposals' for each gland and bulk sample each sample was run through GATK Mutect2 (GATK version 4.0.1)[8]. The non-default parameter "--dont-use-soft-clipped-bases" was added to ensure the bases removed from Skewer and *ClipOverlap* were not reintroduced into the analysis. All resultant vcfs from each set were then merged into a single file representing the union of all variant proposals. A second variant caller, Platypus (version 0.8)[9], was then used to assess the presence of each variant proposal in each gland and bulk sample. Platypus was designed to call haplotypes within genomes, making it suitable for use on the near-clonal glands used in this study. Variants marked with the "allele bias" filter flag were not excluded. The quality control parameters were set as follows: minMapQual=20, minReads=3, maxVariants=100, trimOverlapping=0. To ensure equal calling across the samples the following criteria for each variant was used: minimum depth=10X, minimum variant allele frequency 5%. Note, if a sample failed QC (below) it was excluded from this filtering step.

*Analysis of exome sequencing data: assessing SNV pathogenicity*
The platypus SNVs and indels (both somatic and germline) were annotated using AnnoVar (v 20190409), SIFT (v6.2.1), POLYPHEN (v2) the scores from which were obtained via VEP 92.5. Indels were assessed with VEST (within CRAVAT v4.3). This enabled putative driver mutations to be identified using the following criteria. If a variant was marked as pathogenic with P<=0.05 by one of the above tools, or was a "STOPGAIN" and in a known hotspot (identified through the ICGC database), and was in a tier 1 COSMIC driver epithelial cancer census gene (see: https://cancer.sanger.ac.uk/cosmic/census?genome=37) then it was considered a potential driver.

*Analysis of exome sequencing data: calling CNAs*
The Sequenza R package[10] (version 3.0) was used to call allele-specific copy number alterations. Initially a first-pass analysis on all samples was performed where a broad range of potential cellularity (0.05 to 1) and ploidy (0.8 to 5) values were considered. Following manual inspection of the model fits and breakpoints of each sample, Sequenza was re-run with

narrower priors covering the likely true cellularity and ploidy values. The second run also made use of a table of candidate breakpoints compiled from the aggregate across samples from that cancer of the first run. To construct the candidate table, we used a breakpoint merging routine (custom script) that merged first-round breakpoints that differed by less than a megabase in location.

*Calling absolute copy number where paired exome sequencing data existed*
Each CRC presented in Figure 2 had both sWGS and exome-sequenced glands available. Absolute copy number information derived from Sequenza was used to infer absolute copy number of the sWGS data. Near-clonal CNAs (present in essentially every gland) were assumed to have the same copy number as inferred in the exome sequenced gland(s). The copy number of other segments was then inferred using these known copy-numbers as a reference point.

*Calling absolute copy number from sWGS data in the absence of exome sequencing data.*

In Figure 3, we followed the approach laid out by ASCAT[11] however, as we only had log2 ratio values available due to the low coverage of the data, we sought to leverage multiple sampling to search for ploidy solutions. Here we search across a range of ploidies (1.5-4), allowing solutions to be found with a minimum purity of 0.5 for laser microdissected "region" samples and 0.2 for "bulk" samples. We recorded the per sample fit for each ploidy value. We then recorded the pairwise Euclidean distances of the integer copy number values of each sample divided by the ploidy, filtering for bins with a log2ratio greater than 1 to avoid distances being exaggerated by amplifications. To determine the ploidy of the patient we compared the mean pairwise Euclidean distances of ploidy normalised copy number values to the mean per sample solution for each ploidy value after linearly scaling each measure to the maximum and minimum values, (i.e. minimum equals 0, maximum equal 1). Ploidy solutions were then ranked according to their Euclidean distance from both values being zero. This was performed to ensure the chosen ploidy value produces a good fit both within samples and across samples.

After manual curation the second-best fitting option was chosen for LM0032 and LM0082. Additionally, Liv-C-R1 in LM0131 was determined to be less than 50% pure and was refitted accordingly. The ploidies for fitting copy number profiles of samples Rec-B-B2 (LM0001) and Liv-C-R2 (LM0131) were increased by 0.4 and Liv-G-R1-3 (LM0131) and Liv-B-R1 (LM0089) were decreased by 0.4 to accommodate outlier ploidies in these samples that produced poor fits.

**Sample quality control**
To ensure only samples with adequate tumour content and quality were included in downstream analysis, a sample QC step was introduced.

For exome sequencing, this was: (a) Sequenza-inferred tumour content (see above) minimum 70%, (b) SNV counts >70 and (c) clonal mutations with VAF >0.2.

For the SNP array analysis, the variance of the logR values of bins within putatively diploid segments was considered. Samples that had variance that fell in the top 5% of the population of samples were removed. In addition, samples were removed when the logR values of bins in putatively diploid segments were not normally distributed around the mean (Shapiro-Wilks test). A further 6 samples were removed due to low cellularity.

**Phasing of whole exome and sWGS copy numbers**

An approach similar to published methods[12,13] was used to phase SNVs. In the cases presented in Figure 2, information from whole exome and sWGS was available from the same tumour. Information was combined from these two data types to phase alleles. First the set of SNPs that were detected in all exome sequenced glands was identified. These SNPs were subsetted to leave only those in chromosome copy change regions, and the distribution of the B-allele frequencies (BAFs) was plotted. SNPs phased to either the major or minor chromosomes (i.e. maternal and paternal chromosome) could be observed by being at high vs low BAF respectively. For instance, in a trisomy region the allele imbalances resulted in SNPs shifted to occupy the 0.33 (minor) and 0.67 (major) ranges when purity was 1.

Where coverage permitted, these phased SNPs were then recaptured in each of the sWGS samples. The total number of bases containing either the A or B allele of the SNPs were then used in a statistical analysis that aims to build evidence for the same BAF shift as observed in the exome gland exomes. We used binomial statistics to test whether a significant departure from 0.5 frequency exists in the ratio of the A and B alleles. Note that in addition to verifying the copy calls produced from the QDNAseq method, this phasing routine effective allows us to call copy neutral LOH events in sWGS data as well.

Phased SNPs were then considered in the sWGS data. For each CNA segment, the number of detected major-allele and minor-allele SNPs was reported, and we note that most SNPs were undetected due to low coverage. To test for the expected allelic imbalance, the segmental SNP counts were then assessed with a Fisher's Exact Test with the null expectation based upon the inferred allele-specific copy number.

**Phylogenetic trees construction from SNVs**

PAUP* software[14] was used to build phylogenetic trees from SNV data (Figure 2). Variants were represented as a binary matrix, where the rows corresponded to a particular sample or the normal sample and the columns to a specific variant and binary encoding (0/1) indicated absence or presence of a variant.

A nexus file was constructed for each set to specify the parsimony search parameters as following: (i) the outgroup function was used to "root" the phylogenies in respect to the normal sample; (ii) the *hsearch* function was used to heuristically search 10,000,000 trees; we retained 1,000 of the shortest trees; (iii) the bootstrap routine was used to sub-sampling the tree construction and data 10,000 times. This involved randomly selecting a set of mutations from the binary matrix (with replacement). The percentage of each branch instance was reported in a log file and used to annotate the trees; (iv) the *alltrees* function was used in sample sets with less than 10 samples. This 'brute-force' run makes it possible to acquire the definitely shortest tree(s) from the total tree-space. The resulting .tre files were inspected and converted to .pdf format using FigTree[15]. The homoplasy indexes for the trees were calculated and output as part of the PAUP* log file.

To obtain a p-value for tree balance the Colless's test function in the R apTreeShape package[16] was used. The statistic represent comparison against a "Yule" growth process, meaning the p-value represents the likelihood of imbalance compared to the null expectation provided by this model.

**Assessment of CNA divergence**

Divergence of CNAs across samples within each patient was quantified by computing the pairwise divergence between samples. Specifically, this was the proportion of altered bins (copy number not equal to the base line ploidy in either or both samples) that had different copy number in each sample. When multiple samples were available from each patient, the distribution of all pairwise comparisons was computed and statistical tests on subsets of comparisons performed as described in the main text. This measure of divergence was comparable to others. The following measures were compared for similarity and presented in Fig S2:

1. **Fraction of different bins** (FracDiffBins), defined by counting the number of bins with different CN status in the two samples (gained, lost, baseline) and dividing by the total number of bins across the genome.
2. **Fraction of different altered bins** (FracDiffAltBins), defined by counting the number of bins with different CN status (gained, lost, baseline) and dividing by the total number of bins that are not baseline in at least one of the two samples in a pair. This is presented in the main manuscript, where it is called 'divergence'.
3. **Fraction of different bins weighted by gene content** (GeneDoseFDB) is the same as (1) however each bin is weighted by the number of genes it contains (including the total number of bins) to explore if gene rich/poor areas are increasing/decreasing diversity measurements.
4. **Fraction of different altered bin weighted by gene content** (GeneDoseFDAB) weights each bin by its relative content of genes (weight = number of genes in the bin), and uses the weights to compute diversity as per metric (3).
5. **Genetic distance** is the sum of the difference between integer copy numbers in each bin as opposed to a binary same/different criteria for baseline, gains and losses. The total sum is divided by the number of bins across the genome.
6. **Breakpoint divergence** measures the amount of non-overlapping breakpoints as a proportion of the total breakpoints present in a pair of samples.

All diversity statistics were highly correlated with one another (Fig S2). Importantly, the correlation between statistics that normalised for the proportion of the altered genome and those which did not (e.g. FracDiffBins v FracDiffAltBins) was very high (Pearson correlation coefficient = 0.742) showing that our measurement of low CNA diversity in cancers relative to adenomas was not an artefact of the choice of statistic.

Linear mixed-effects models that control for inter-patient differences were calculated using the lmerTest package[17] in R.

*Genome Duplication Classifier*

Using ICGC segment calls we binned the genome into 4Mb bins to mimic data expected in low coverage WGS. Here we took the median calls for each bin when comparing to overlapping segments. For each sample we calculated the number of segments after binning by detecting changes in call status in adjacent bins across the genome. We calculated PGA by measuring the fraction of bins across the genome that did not equal the median CN of the bins of the sample (baseline copy number). Taking the called GD status per sample from the ICGC resource we trained a support vector machine using the e1071 package[18] in R (no scaling, a linear kernel and a cost of constraints violation of 5). The classifier to these data calls GD as present in a sample if:

$$0 < 0.000355NS + PGA - 0.432 \qquad (1)$$

(Where NS denotes the number of segments, PGA the percentage genome altered [fraction of genome with non-baseline copy number]). On PCAWG data, this classifier calls 83.8% of GD samples correctly, and 93.2% of non-GD correctly (Fig S4).

We determined an 'ambiguous' call zone ±0.5 of the beta0. To classify patients with multiple samples patients were considered clonally non-GD if at least one sample was 'confidently' called non-GD and no samples were call 'confidently' called GD and vice versa. Subclonal GD was called when a patient has both 'confident' non-GD and GD calls. Ambiguous calls per sample were then corrected according to the patient assessment, in subclonal GD patients ambiguous calls were corrected to the confident call that had the highest frequency in the patient and if this was tied we corrected to ambiguous calls to GD.

*Phylogenetic analysis of sWGS data*

Copy number data across all samples for a given patient were split up based on breakpoints detected in at least one sample (defined as a copy number difference between two consecutive bins) to generate a comparable set of regions. Regions containing less than 4 bins were removed for phylogenetic analysis. Phylogenetic trees were created using MEDICC2 in the total copy number mode[REF]. Trees were plotted using ggtree[REF]. Reconstructed MEDICC2 unobserved copy number states were used to calculate clonal and tip frequencies. Clonal changes represented gains and losses from the root and tip changes presented alterations present between the tip nodes and the nearest internal node. Gains and losses were only counted once even if they resulted in a change in copy number greater than 1 and therefore more than 1 copy number event. Clonal events were normalised to the ploidy of the patient to avoid all regions being gained in triploid and tetraploid tumours.

*Copy number quantification in DLP+ scDNA-seq*

Demultiplexed dual-index fastq files were obtained for single cells undergoing the DLP+ protocol and the data were used as input for the workflow automation pipeline designed for the DLP+ method (https://github.com/shahcompbio/single_cell_pipeline) using default settings. First, raw reads were adapter-trimmed using TrimGalore, mapped with BWA aln to the hg19 reference genome and deduplicated using picard MarkDuplicates. Subsequently, copy number calling was performed using the HMMcopy tool with reads segmented into non-overlapping 500kb genomic regions. Cells were required to have a minimum quality score of 95, a minimum of 250,000 reads and an S-phase probability of less than 0.1. Only bins considered "ideal" in all cells were used.

*Copy number quantification in 10X scDNA-seq*

As per the 10X recommended analysis, CNAs were determined using CellRanger with default settings (using GRCh38 as the reference genome). For each cell the mean event confidence weighted by segment size and number of segments was calculated and a minimum mean event confidence of $10^{1.85}$ was required per cell and a minimum of 416 segments was allowed. Calls were then binned into 1Mbp bins and bins with no overlapping segments in at least one cell were removed. Cells with more than 50% of the genome with copy number zero were removed, as were cells with a mean copy number of more than 20. A maximum of 105 segments based on final bins were allowed per cell.

*Phylogenetic analysis of scDNA-seq data*

Comparable regions were calculated for the SW620 and 3994-117 cells according to the sWGS approach. For both sets of cells, ploidy was limited to less than the mean ploidy + 1 to remove potential G2 phase cells or doublets. A minimum of 2 bins was required for regions for both populations. A random sample of 100 3994-117 cells were selected to construct the tree to make it comparable to the SW620 tree. MEDICC2 was used, trees were plotted, and clonal and tip frequencies were calculated, as previously described for sWGS.

**General statistical analyses**

All statistical comparisons were performed in the R statistical programming language[19], with specific statistical tests used described in the manuscript.

**Mathematical modelling of CNA evolutionary dynamics and statistical inference**

To model the fitness effects of chromosomal instability a computational framework called *CINulator* was developed. *CINulator* models tumour evolution as a birth death process where cells can gain or lose loci at a specified rate $\mu$. Changes in copy number at a locus may influence the birth and/or death rates. Fitness is incorporated by defining a fitness landscape where the optimum is some pre-specified genotype $G_O$, the fitness of individual cells is proportional to the distance from this optimum genotype:

$$s = \frac{s_{max}}{1 + \alpha |G_O - G_i|}$$

With fitness defined as the net growth rate - $s = b - d$. Changes in fitness may either increase the birth rate or decrease the death rate. The simulation begins with a single cell that has some genotype $G$ which can be at, or close to the optimum. In this scenario the majority of changes will bring a fitness cost. Alternatively, if the genotype of the founder cell is far away from the optimum, then the majority of changes will give a fitness benefit and the population will tend to climb the fitness landscape.

We also constructed a Bayesian inference framework, based on an Approximate Bayesian Computation method, to compare model predictions to data, and learn the model parameters that best described the observed data.

A full description of the model and inference framework is provided in the supplementary mathematical note, details of the various assessments of model dynamics, and details of the statistical approach.

**Data availability statement**

Processed data sufficient to reproduce the analysis in each figure is presented as supplementary tables, referenced throughout the manuscript. All raw sequencing data and arrays files will be available through the EGA repository, accession number: EGAS00001004219. Code used for analysis is available on github: https://github.com/BCI-EvoCa/CNA_stability and simulation code is at https://github.com/ucl-cssb/CIN_CRC .

19. R Core Team. R: The R Project for Statistical Computing. https://www.r-project.org/.
