## Supplementary note on modeling of glands for "Negative selection may cause grossly altered but broadly stable karyotypes in metastatic colorectal cancer"

### Stochastic modelling of individual glands

To test whether the lower diversity observed in copy number alterations (CNAs) from individual colorectal cancer (CRC) glands (crypts) can be better explained by negative selection or not, we utilized a Bayesian approach to do parameter inference and model selection (Supplementary Note Figure 1). We used a stochastic birth-death branching process to simulate the gland growing process under the model without and with negative selection. Then we used approximate Bayesian computation (ABC) to infer parameters of each model by comparing real data to simulated data. Finally, we used deviation information criterion (DIC) to find which model is better supported. We did extensive tests via simulations to ensure our approach is valid before applying it to the real data.

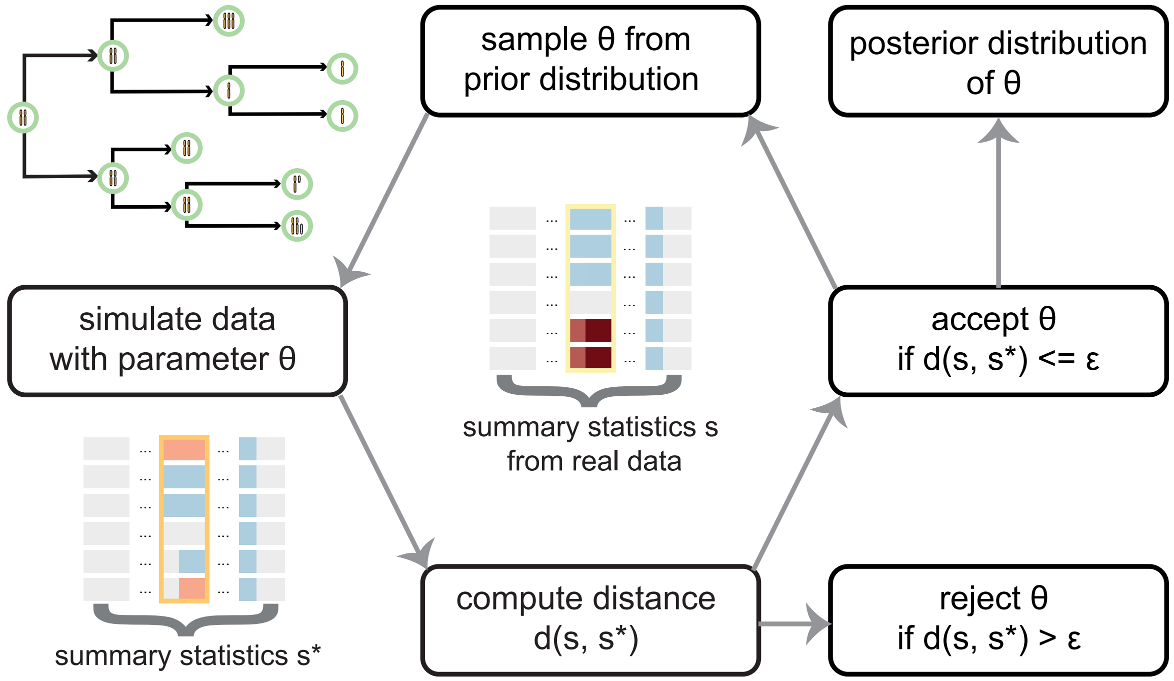

**Supplementary Note Figure 1** The framework for stochastic modelling of colorectal cancer glands.

We used the above Bayesian inference approach to infer the strength of negative selection acting within each of the 6 cancers subjected to gland-by-gland analysis (Fig 2A). We initialised simulations with a single gland that had the putative "optimal" core karyotype that was set to the modal karyotype observed in the tumour. In an initial investigation we realised that we could not separately infer the mutation rate ($\mu$) and parameter controlling the strength of negative selection ($\alpha$), as low CNA diversity could be equally well explained by a low mutation rate combined with strong negative selection, or *vice versa* (Fig S13A-C). However, our simulations showed that there is often more support for the (simpler) model without negative selection than the (more complex) model that included negative selection. Hence, the power to detect the strength of negative selection is low, and therefore our method was conservative in detecting negative selection. Simulations also showed that mutation rates tended to be accurately estimated, but accuracy decreased in the negative selection model. The posterior distributions of $\alpha$ had a wide range, but it was sufficient to distinguish low and high negative selection strengths.

We describe our methods to obtain these results in more detail in the following sections.

### 1 Simulation of individual CRC gland data

#### 1.1 Branching process to simulate evolutionary dynamics of glands

A human colon gland typically has stem cells at the base, above which are transient amplifying cells and differentiated cells [1]. The stem cell can divide in three different ways: asymmetric division which generates one stem cell and one differentiating cell; symmetric division which either generates two stem cells or two differentiating cells. If a stem cell has new mutation(s) making it outcompete other stem cells, it may expand to dominate the whole gland (monoclonal conversion). This gland may then spread to neighbouring regions through gland fission, which is the bifurcation of itself into two daughter glands. Gland fission may be complemented by gland fusion, in which two glands merge into one.

The evolutionary dynamics of glands are hence very complex, making it hard to simulate full details in an efficient way. As our aim is to determine whether negative selection is better supported by the observed CNAs, we determined to only model the basic gland growing process. Because an individual gland is often occupied by a stem cell and its decedents, we considered each gland as a homogeneous population of cells and hence the basic unit of the stochastic branching process. Following the hypothesis that new CNAs accumulated in cancer cells at fitness peak are preferentially removed by negative selection, we assumed the branching process starts from a gland with the putative "optimal" core karyotype $G_o$. The gland divides at rate $b$ and dies at rate $d$. The average number of glands at time $t$ is then $N(t) = e^{(b-d)t}$. After each division (fission), a number of CNAs accumulate in the daughter glands according to Poisson distribution at mutation rate $\mu$. The mutational process can be seen as a simplification of the generation of a mutant stem cell and the subsequent monoclonal conversion. Each CNA either increases or decreases a copy of a genomic segment in (the stem cell dominating) the gland. Under the model with negative selection, a gland $i$ with karyotype $G_i$ has fitness cost controlled by $\alpha$ and the karyotype difference $d(G_i, G_o)$. The fission (birth) rate $b_i$ and death rate $d_i$ of gland $i$ are specified by $1 + \alpha * d(G_i, G_o) = \frac{b_o - d_o}{b_i - d_i}$, where $b_o$ ($d_o$) is the birth (death) rate of the initial gland. If we denote the net growth rate of gland $i$ as $\lambda_i = b_i - d_i$, then $\lambda_i = \frac{\lambda_o}{1 + \alpha * d(G_i, G_o)}$. With larger $\alpha$ and/or larger karyotype differences, $\lambda_i$ will be smaller.

Suppose the karyotype of gland $i$ is $G_i = (c_1^i,\ c_2^i,\ \ldots,\ c_n^i)$, where $n$ is the number of bins across the genome and $c_k^i$ is the copy number at bin $k$. The number of altered bins from the optimum karyotype is $n_a = \sum_{k\in\{1,2,\ldots,n\}} I_{ko}$, where $I_{ko} = 1$ if $c_k^i! = c_k^o$ and $I_{ko} = 0$ otherwise. The karyotype difference $d(G_i, G_o)$ can then be defined in multiple ways, such as PGA (percentage genome altered, $\frac{n_a}{n}$) and $L_1$ distance between copy number vectors ($|G_o - G_i|$). Because the fitness cost depends on the product of $\alpha$ and $d(G_i, G_o)$, given a fixed fitness cost, $\alpha$ and $d(G_i, G_o)$ are inversely correlated. For example, when $d(G_i, G_o) = \frac{n_a}{n}$, the karyotype difference is generally very small because $n_a$ is much smaller than $n$ and hence a very large $\alpha$ is required to get intermediate fitness cost. On the contrary, when $d(G_i, G_o) = |G_o - G_i|$, the karyotype difference can be huge due to many small CNAs and hence even a very small $\alpha$ can indicate a large fitness cost. Moreover, these two definitions of karyotype difference take into account the size of CNAs, which introduces more stochasticity. To obtain a reasonable range of $\alpha$, we defined $d(G_i, G_o) = n_m$, where $n_m$ is the number of new mutations present in $G_i$ but not in $G_o$.

#### 1.2 Analysis of the branching process

It is difficult to analyze the model analytically, so we did an approximate analysis. Under the model without negative selection, the karyotype diversity of the gland population at an earlier time forms a lower bound on the diversity later, because more mutations are accumulated. Under the model with negative selection, the diversity is expected to be lower with any amount of selection. Assuming an infinite sites model, we notice that $G_i$ must be increasingly different from $G_o$ as new mutations arise. Then $d(G_i, G_o)$ will be increasing and $\lambda_i$ will be decreasing when $\alpha$ is fixed. Furthermore, assuming that the accrual of mutations is linear in time, then the number of new mutations in gland $i$ at time $t$ is $M_i = \mu t$ and we can write $\lambda_i \approx \frac{\lambda_o}{1+\alpha\mu t}$. This means that the estimation of $\mu$ and $\alpha$ is essentially non-identifiable and only their combination matters. Because we are more interested in model selection rather than parameter estimation, we still use this model since it is easy to understand and simulate.

To better understand the imposed selection strengths in the model, we explored the relationship between $\alpha$, karyotype difference $d(G_i, G_o)$, and standard selection coefficient $s$ (Supplementary Note Figure 2), which is defined by $s = \frac{1}{1 + \alpha * d(G_i, G_o)} - 1$, assuming $1 + s = \frac{b_i - d_i}{b_o - d_o}$. The values of selection coefficient $s$ directly indicates the fitness of a gland $i$. If $s > 0$ ($-\frac{1}{d(G_i, G_o)} < \alpha < 0$), gland $i$ has fitness advantages and will grow faster. If $-1 < s < 0$ ($\alpha > 0$), gland $i$ will be selected against and grow slower. Here, the positive selection strengths are very sensitive to changes of $\alpha$. Since our model is specifically designed to detect negative selection, we limit $\alpha > 0$ in our inferences. When $\alpha$ increases, the relative change of $s$ becomes smaller, especially with larger karyotype differences. This suggests that the relative differences of the fitness costs of glands at a certain generation during the exponential growth are very small when mutation rate and $\alpha$ are large. In the branching process model, one gland is randomly picked from the current population to grow at a time. When all glands have similar selective advantages at the same time, it is less likely for a gland to expand and provide a stronger signal of selection in the final population. We also ran some simulations with increasing death rate of a gland under selection. However, more generations are required to reach $N$ glands, which generates relatively more mutations in the end and further decreases the power to detect selection. Therefore, our model may not have enough power to distinguish different values of $\alpha$.

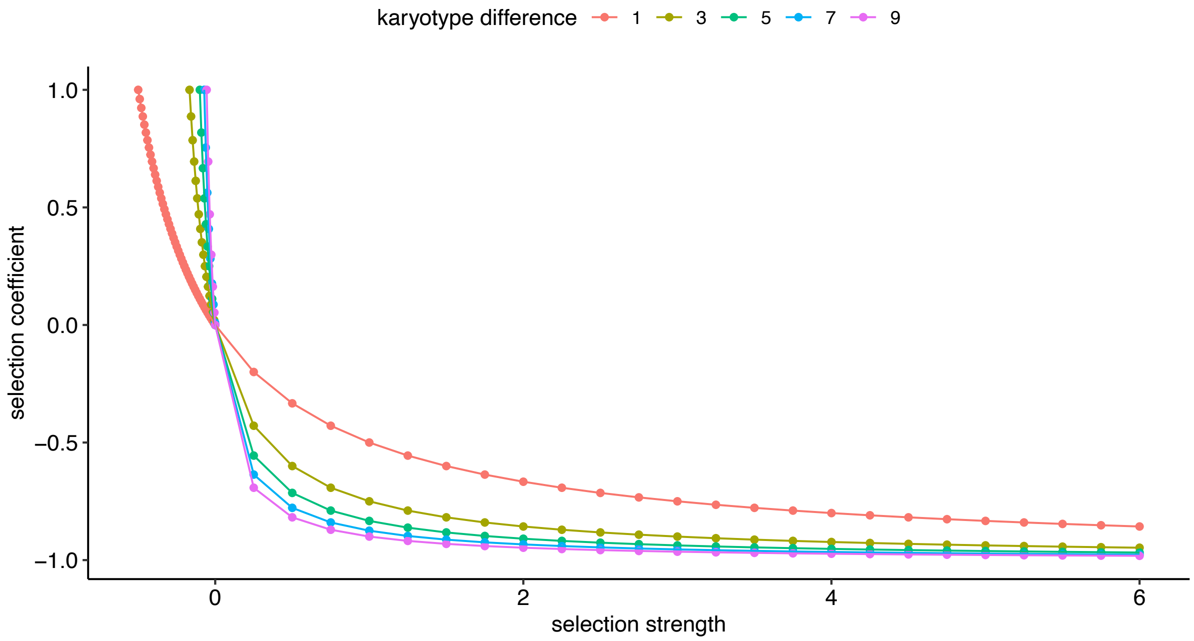

**Supplementary Note Figure 2** The relationship among selection strength $\alpha$, selection coefficient $s$, and karyotype difference $d(G_i, G_o)$ in the stochastic branching process model.

#### 1.3 Implementation of the branching process

We implemented the branching process with rejection-kinetic Monte Carlo algorithm [2] in C++. The program is available at https://github.com/ucl-cssb/CIN_CRC.

We ran the branching process until $N$ glands. It is computationally unfeasible to simulate millions of glands in a real tumour. As a balance between biological meaning and computational efficiency, we simulated the gland growing process until $N = 1000$. We discuss how the choice of $N$ affect inference in real data in Section 4.2. To consider the spatial relationships among glands which were sampled from different sides of a patient, we sampled from two sides of the lineage tree to get the same number of glands as measured in real data. We fixed $b_o = 1$ and $d_o$ = 0, since only relative growth rate matters here. When new mutations arise in a gland, giving rise to negative selection, the birth rate of this gland will be decreased.

To get more realistic simulated data, we incorporated CNAs detected in real data in the simulation. The real data here include all the exome sequencing and shallow whole genome sequencing samples for six patients C267, C274, C277, C282, C288, and C319. We excluded two exome samples in C274 (CE2_P6_22 and CE2_P6_23) which have much more unique CNAs. We considered the representative genome of a gland as a set of equal-size bins (500 Kbp) to be consistent with copy number calling in real data. The size of the last bin in each chromosome is determined by the chromosome boundary. We only considered 22 autosomes and hence there were 5776 bins in total. Since just a few copy neutral loss of heterozygosity event were detected, we only simulated total integer copy numbers.

We assumed that the clonal CNAs more frequently present in each patient represent the "optimum" core karyotype, so we computed the modal copy number for each bin across all the samples for each patient and used the obtained copy number vector as the core karyotype in the simulation (Supplementary Note Figure 3). The modal karyotype provides a baseline for copy number changes in simulated CNAs, which makes it easier to see the differences between simulated and real CNAs. Since what matters more in the simulation is the relative copy number changes, a different choice of the core karyotype will not affect the subsequent inferences too much. To reduce the effect of potential noise in calling short CNAs and retain as much information as possible, we considered CNAs of at least eight bins (4 Mbp). We mention the choice of the number of bins in Section 4.1. The size of each CNA (in the unit of bins) was sampled from the sizes of observed CNAs in each patient (Supplementary Note Figure 4). The location of each CNA was sampled based on the number of unique copy number changes in each bin across all samples in each patient. We assumed the same probability of copy number gain and loss.

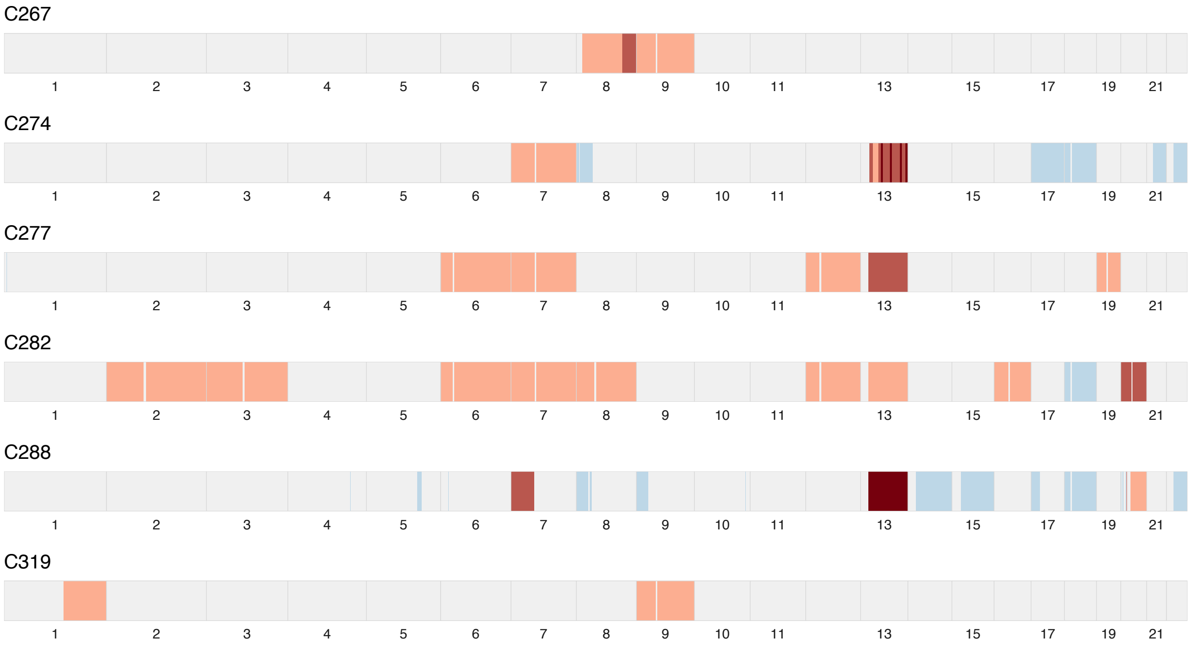

**Supplementary Note Figure 3** The core (mode) karyotype of each patient, which is the initial state of the simulated tumours. The colour coding of copy number: light blue: 1; light grey: 2; light red: 3; red: 4; dark red: 5.

The numbers of glands sampled from each side of the gland lineage tree were set according to the number of samples taken for each patient (Supplementary Note Table 1). For patients with samples taken at more than two sides (C282 and C319), the adjacent glands were combined.

To track the relationships between sampled glands, we allowed the output of their complete lineage which was obtained by recording the ID of each gland generated in the process. We may then extract the sampled glands from the complete lineage tree to get the phylogenetic relationships among the samples.

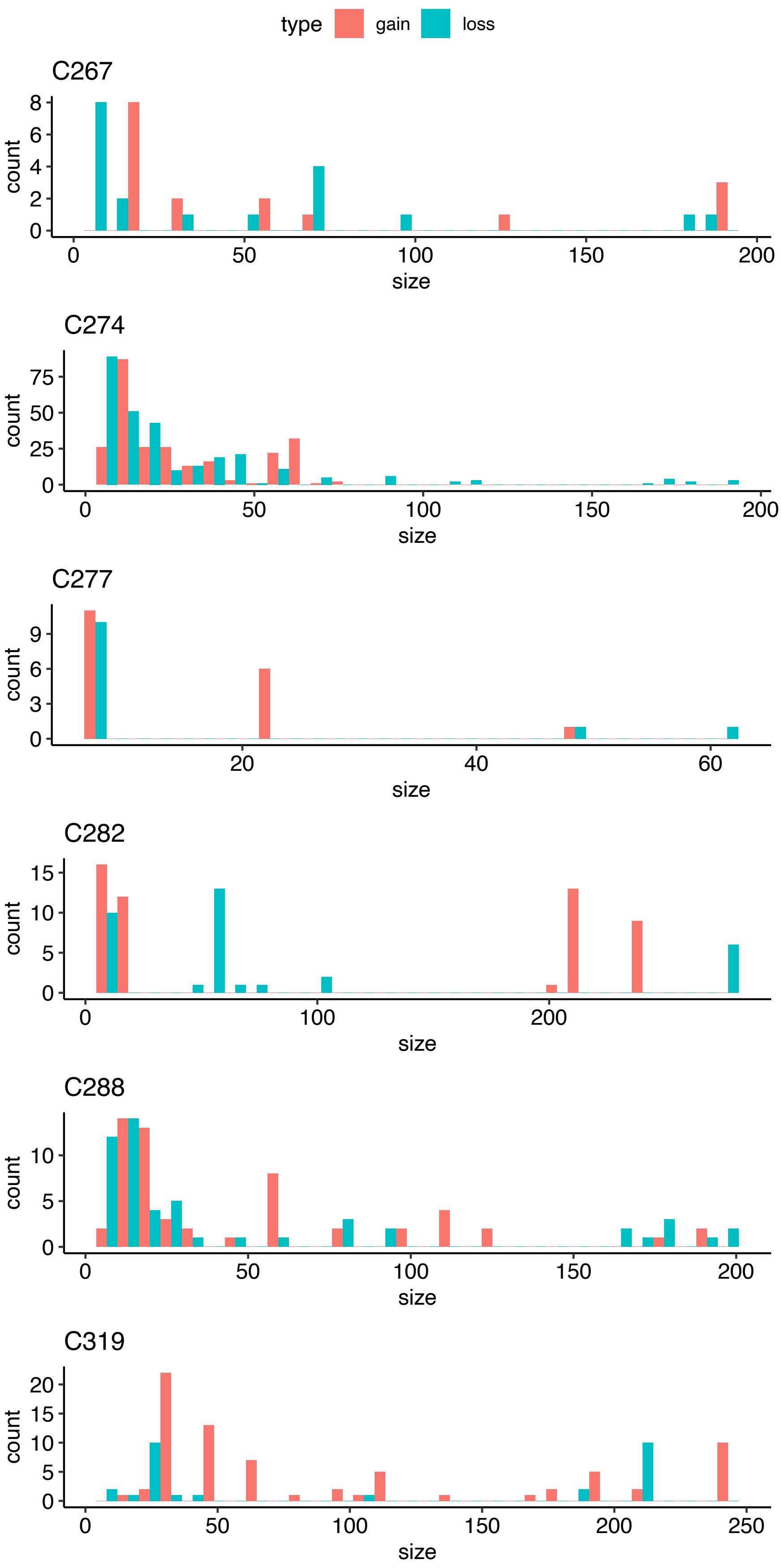

**Supplementary Note Figure 4** The distributions of copy number gain/loss sizes (in term of bins of size 500 Kbp) in each patient. Only alterations of at least eight bins are included.

**Supplementary Note Table 1** The number of glands sampled from each patient.

| Patient | #total glands | #glands at side 1 | #glands at side 2 |
|---|---|---|---|
| C267 | 43 | 32 | 11 |
| C274 | 56 | 37 | 19 |
| C277 | 39 | 20 | 19 |
| C282 | 51 | 34 (2 + 32) | 17 |
| C288 | 39 | 33 | 6 |
| C319 | 77 | 26 | 51 (42 + 3 + 3 + 1 + 2) |

#### 1.4 Simulation starting from diploid karyotype

To show how negative selection suppresses karyotype diversity when the tumour genome is close to fitness peak, we implemented another simulation approach which starts from diploid karyotype and evolves towards "optimal" core karyotype. Because the karyotypes may become too divergent to reach fitness peak (optimum) in a considerable amount of time if focal CNAs are incorporated, we only considered chromosome-level CNAs in this simulation mode. Since the chromosome-level copy number vector is much smaller compared with the vector obtained from bin-level CNAs, we measured the distance between karyotypes by $L_1$ distance when updating the fitness of daughter glands. To be consistent with the simulation starting from core karyotype, we also maintained a bin-level copy number vector by mapping the chromosome-level CNAs to the corresponding bins. We assigned a weight $w$ (2 by default) to the chromosomes with CNAs in the core karyotype. If $w > 1$, those chromosomes are more likely to undergo CNAs, which may reach optimum more quickly. To further accelerate the speed to reach optimum, we changed the fitness of a gland by solely increasing its death rate which starts from a non-zero value and keeping the birth rate to be 1. We introduced a bottleneck to model the seeding of a metastasis. Namely, we simulated the population until it reaches $m$ (1000 by default) glands and then randomly selected $n$ glands to start a new population.

Using this simulation approach, we explored the evolutionary dynamics when a cancer was initiated by a gland with a suboptimal karyotype by simulating chromosome-level CNAs (Fig S12A). In this case, the cancer population experienced directional selection towards the optimal karyotype (Fig S12B-D), and notably CNA diversity of the population increased transiently before the optimal karyotype was found by the population. The transient increase in CNA diversity was explained by the coexistence of multiple 'independent' clones with distinct CNA karyotypes that all had comparable fitness (equidistant from the fitness peak on the symmetrical fitness landscape; Fig S12C-E). Later in the evolutionary process, these clones were replaced by a single population (potentially composed of multiple convergently-evolved lineages) with the same optimal CNA karyotype. This transient increase in diversity and accrual of CNAs driven by directional selection were consistent with our observations of high CNA heterogeneity followed by a bottleneck during adenoma to cancer evolution (Fig 1).

### 2 Parameter inference and model selection with ABC

ABC is a flexible and powerful approach to infer parameters in a wide range of models when the likelihood function is intractable [3]. Basically, a large number of simulated datasets are generated from certain prior distributions of parameters in a pre-specified model. The simulated dataset is compared with the real data by some distance function to see how similar they are. If their distance is smaller than a target tolerance, the simulated dataset will be accepted. All the accepted simulations compose the posterior distributions of the parameters of interest in the model. It is often hard to use the full dataset for inference, so summary statistics extracted from the data are usually used instead.

#### 2.1 Choice of summary statistics

Since it is hard to directly use the copy number data for ABC, we came up with a set of informative and low-dimensional summary statistics based on biological background and tests with simulations. These summary statistics include PGA relative to the initial karyotype ($s_1$), average pairwise divergence ($s_2$), variance of pairwise divergences ($s_3$), and number of unique breakpoints across all sampled glands ($s_4$).

For the computation of PGA, we defined that a bin is altered if its copy number is different from the corresponding copy number in the initial karyotype and then PGA is the percentage of bins that are altered. Suppose there are $m$ glands, $G_i =$

$\left(c_1^i,\ c_2^i,\ \ldots,\ c_n^i\right)$, and $G_o = \left(c_1^o,\ c_2^o,\ \ldots,\ c_n^o\right)$, then $s_1 = \frac{\sum_{k\in\{1,2,..n\}} I_{ko}}{n}$ where $I_{ko} = 1$ if $\exists c_k^i ! = c_j^o$ $(i \in \{1,2,\ldots,m\})$ and $I_{ko} = 0$ otherwise.

We defined pairwise divergence between two glands as the proportion of different bins that are altered (or proportion of aberrant genome that are subclonal). For its computation, we compared each pair of glands to obtain the number of bins with different copy numbers and then divided it by the number of sharing bins which have copy numbers not equal to the ploidy (2 for all the patients in consideration). Assuming $G_i = \left(c_1^i,\ c_2^i,\ \ldots,\ c_n^i\right)$ and $G_j = \left(c_1^j,\ c_2^j,\ \ldots,\ c_n^j\right)$, the pairwise divergence between $G_i$ and $G_j$ is $\frac{\sum_{k\in\{1,2,..n\}} I_{kd}}{\sum_{k\in\{1,2,..n\}} I_{ka}}$, where $I_{kd} = 1$ if $c_k^i ! = c_k^j$ and $I_{kd} = 0$ otherwise; $I_{ka} = 1$ if either $c_k^i ! = 2$ or $c_k^j ! = 2$ and $I_{ka} = 0$ otherwise. Let $D$ be the set of all pairwise divergences, then $s_2 = mean(D)$ and $s_3 = var(D)$. For short, we use divergence to denote average pairwise divergence too.

We also computed the number of unique mutations across all sampled glands and average pairwise different number of mutations in simulated data. The former is similar to the number of segregating sites in the context of single nucleotide variants, whereas the latter is similar to Tajima's $\pi$ (average number of pairwise differences). Their distributions are consistent with the other summary statistics (Supplementary Note Figure 5).

We checked how the summary statistics change with different numbers of glands per tumour, $N$, under different selection strengths assuming $\mu = 0.2$ (Supplementary Note Figure 6). As a larger $N$ indicates more mutations are simulated, the summary statistics values increase with $N$. Simulated tumours with more glands and larger fitness cost may have summary statistics similar to those with less glands and smaller fitness cost.

#### 2.2 DIC for model selection

After inferring the posteriors of parameters with ABC under each model, we did model selection by computing DIC which measures both goodness of fit to data and model complexity [4]. DIC can be easily computed from posterior predictive distributions of parameters. The expression for DIC we used is: $DIC = 2\bar{D} - D(\bar{\theta})$, where

$$D(\bar{\theta}) = -2\,log(\frac{1}{n}(\sum_{j=1}^{n} K(s - s_j^*)),$$

$$\bar{D} = -\frac{2}{m}\sum_{i=1}^{m} log(\frac{1}{n}(\sum_{j=1}^{n} K(s - s_j^*)),$$

$$K(s - s_j^*) = \frac{1}{\sqrt{2\pi}}\, e^{-\frac{(s-s^*)^2}{2}}.$$

Here, $s^*$ is the simulated summary statistics and $\bar{\theta}$ is the posterior mean of $\theta$. We computed DIC with $m = 100$ and $n = 100$.

The range of DIC is affected by the distance between simulated and real summary statistics and the target tolerance $\epsilon$. A smaller DIC value typically indicates the corresponding model is better supported by the data. But it is hard to interpret the absolute values of DIC in practice. When the target tolerance is higher, DIC is typically larger. If the distance between simulated and real summary statistics has a wider range, DIC is also larger. Therefore, when the target tolerances are similar for inferences under two models, the relative differences (even with small values) of DIC values seem reasonable in determining which model is better.

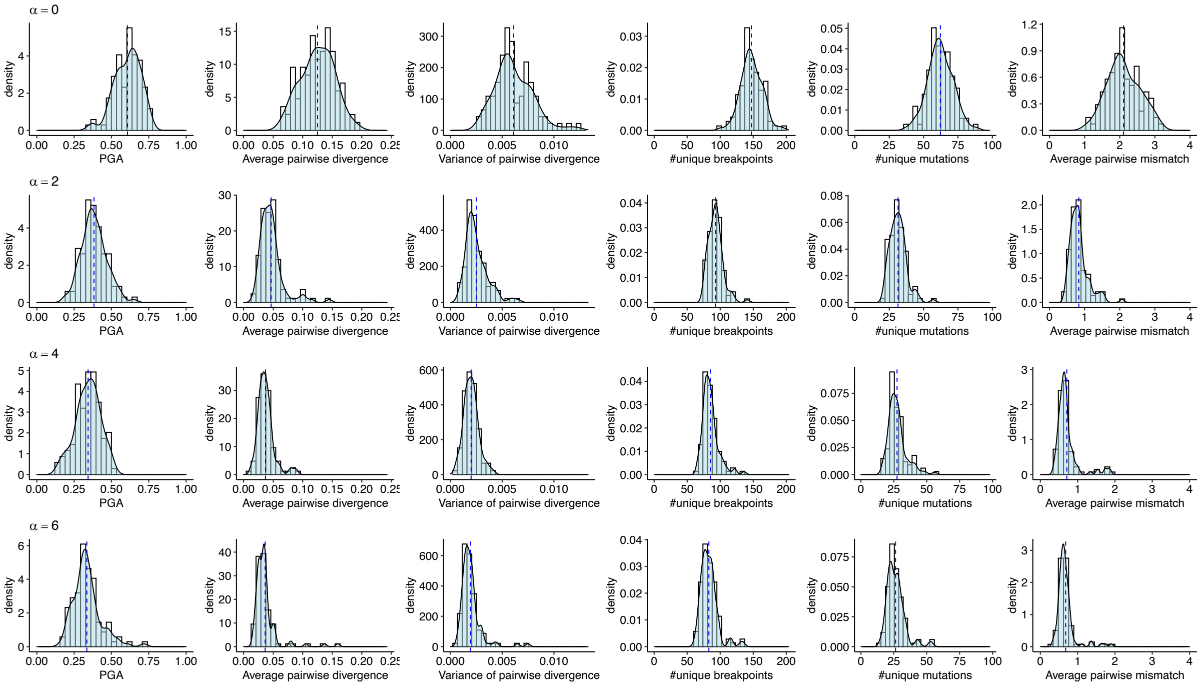

**Supplementary Note Figure 5** The distributions of summary statistics computed from 100 simulations based on data of C282 with $\mu$ being 0.2 and $\alpha$ being 0, 2, 4, and 6 respectively.

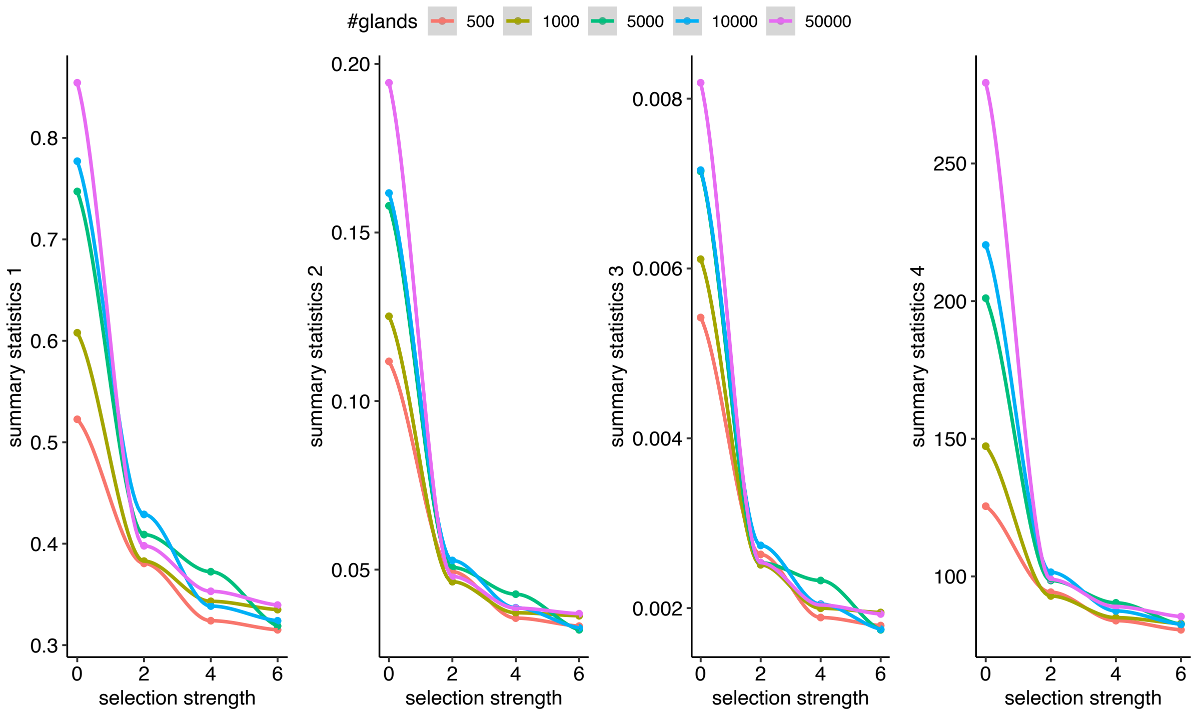

**Supplementary Note Figure 6** Changes of summary statistics when the simulation stopped at different number of glands under different selection strengths (based on data of C282, $\mu = 0.2$). Each point is the mean value over 1000 simulations.

### 3 Validations on simulated data

#### 3.1 Power analysis to select best summary statistics

To validate our approach, we checked how its power in detecting correct model varies with different mutation rates and selection strengths on simulated data. Due to computational efficiency, we used the ABC rejection algorithm to do parameter inference. We generated 100,000 simulations beforehand according to pre-specified priors under each model based on data of patient C282. To get the posterior distributions, we took the parameter values of the top 1% of the 100,000 simulations ordered by Euclidean distance to the observed summary statistics under each model. We computed DICs one time for each simulated target dataset under both models ($m = 100, n = 100$) and counted the number of times where one model is better supported than the other. To be consistent, we assumed the priors are $\mu \sim Unif(0, 0.5)$ and $\alpha \sim Unif(0, 6)$ for all the inference on simulated data.

To find the best set of summary statistics for inference, we did model selection on 120 target datasets (20 replicates for each combination of $\mu$ and $\alpha$) generated with $\mu$ being 0.1, 0.2, 0.3, 0.4 and $\alpha$ being 0, 2, 4 respectively. We tested different combinations and weightings of the four selected summary statistics ($s_1$: PGA, $s_2$: average pairwise divergence, $s_3$: variance of pairwise divergence, $s_4$: number of unique breakpoints). To avoid missing potential information on the distribution of pairwise divergence, we included both mean and variance of pairwise divergence. We used either number of unique breakpoints ($s_4$) or PGA ($s_1$), since they measure similar information. We computed average fraction of false negative predictions (when the true model is not detected) for inferences on target datasets with different sets of parameters (Supplementary Note Figure 7).

The results suggest that set 1 ($s_1$, $s_2$, $s_3$, without scaling by standard deviation of prior predictive distribution), set 6 ($s_2$, $s_3$, $\log(s_4)$, without scaling by standard deviation of prior predictive distribution), and set 8 ($s_2$, $s_3$, $\frac{s_4}{500}$, without scaling by standard deviation of prior predictive distribution) can detect the model without negative selection when it is true in most cases whereas the power to detect the model with negative selection is weak. Using set 4 ($s_1$, $\frac{s_2}{0.5}$, $\frac{s_3}{0.02}$, without scaling by standard deviation of prior predictive distribution) has a slightly weaker power in detect the model without negative selection  but a much stronger power to detect the model with negative selection (Supplementary Note Figure 8). To be more conservative in detecting negative selection, we chose to use set 1 for further inferences (Supplementary Note Figure 8). The ranges of the $s_1$, $s_2$, and $s_3$ are different, which naturally assigns different weights to them.

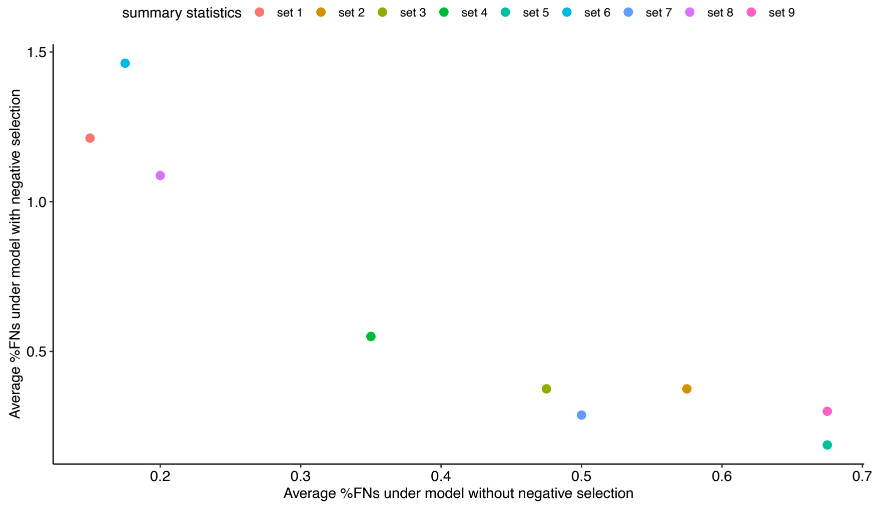

**Supplementary Note Figure 7** The fraction of false negative (FN) model selection results with different sets of summary statistics, including "s1s2s3sd0", "s1s2s3sd1", "s2s3s4sd1", "s1s2s3sd0_10.50.02", "s1s2s3sd1_10.50.02", "s2s3s4sd0_11log", "s2s3s4sd1_11log", "s2s3s4sd0_11500", and "s2s3s4sd1_11500". The names are indicated by sID1sID2sID3sdX, where each summary statistics was scaled by the standard deviations of its prior predictive distribution. The suffix after "_" indicates additional weighting factors for each summary statistics before scaling by standard deviation, where "1" means no change, "0.5" means scaling by 0.5, "0.02" means scaling by 0.02, "log" means scaling by log, and "500" means scaling by 500.

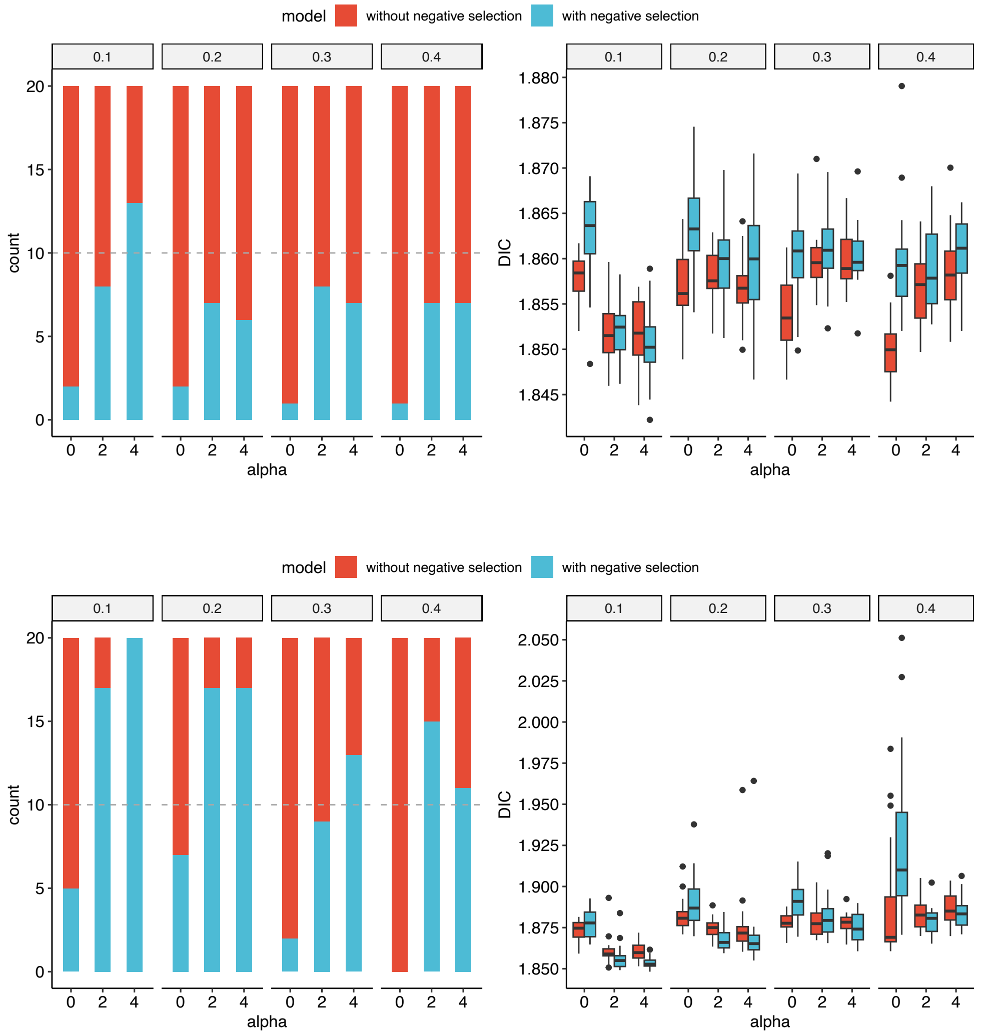

**Supplementary Note Figure 8** Results of power analysis on model selection using set 1 (top) and set 4 (bottom). Left: The bar plot of the number of times each model is supported. Right: The boxplot of DIC values under two models. The plots are grouped by mutation rate.

#### 3.2 Validation of selected summary statistics

We also validated whether the selected summary statistics can accurately recover the true parameters with the ABC rejection algorithm (Supplementary Note Figure 9). We simulated data with different values of $\mu$ and $\alpha$ as true data based on C282. To get the posterior distributions, we took the parameter values of the top 1% of 100,000 simulations ordered by Euclidean distance to the observed summary statistics of simulated “true” data. We estimated mutation rate $\mu$ under the model without negative selection with varying true $\mu$ from 0.05 to 0.45 with a step of 0.05. The posterior mean of estimated $\mu$ is close to real value in all the cases. Due to the correlation of $\alpha$ and karyotype difference, it is harder to infer the exact value of $\alpha$. We estimated $\alpha$ when $\mu$ is fixed to be 0.2 under the model with negative selection. The posterior distributions of $\alpha$ have a wider range, but it is sufficient to distinguish low and high selection strengths.

The basic ABC rejection algorithm has low acceptance rate when the prior distribution is very different from the posterior distribution, so we did further validations with the more efficient ABC SMC algorithm [3]. In the ABC SMC algorithm, a set of sampled parameter values are propagated through a series of intermediate distributions (populations) with decreasing tolerances until the target tolerance $\epsilon_T$ is reached ($\epsilon_1 > \epsilon_2 > \cdots \epsilon_T \geq 0$). Each sampled parameter value (particle) also gets a weight based on its importance. We used the function ABCSMC in the Julia package ApproxBayes (https://github.com/marcjwilliams1/ApproxBayes.jl) to run the algorithm. We took 500 particles from each population to estimate the posterior distributions and set the number of maximum simulations to $10^6$ (which is equivalent to an acceptance ratio of < 0.005). We set most parameters in the algorithm to be default: the first tolerance value is $10^4$; the 0.3th quantile of $\epsilon_i$ is the value of $\epsilon_{i+1}$; and the program stops when $\epsilon_{i+1}$ is within 0.05 of $\epsilon_i$. We mainly varied the prior distributions of parameters and target tolerance.

We simulated data with mutation rate varying from 0.1 to 0.4 with a step of 0.1 under both models based on the karyotypes of C319. We fixed $\alpha = 1$ under the model with negative selection. Then we used ABC SMC (target tolerance 0.005) to infer $\mu$ under both models with prior $\mu \sim Unif(0, 0.5)$ and $\alpha$ under the model with negative selection with prior $\alpha \sim Unif(0,6)$. The results suggest that the mutation rates can be accurately estimated under both models (
Supplementary Note Figure ***10***), although the posteriors under the model with negative selection have a wider range. As expected, the estimated $\mu$ and $\alpha$ are correlated.

#### 3.3 Sensitivity analysis on parameter estimation

To know how well the parameters ($\mu$ and $\alpha$) can be estimated, we did a sensitivity analysis (Supplementary Note Figure 11). We compared the distributions of the number of mutations in the simulated $N = 1000$ glands under different parameter settings and computed their Kolmogorov-Smirnov distances to the distribution at $\mu = 0.1$ and $\alpha = 0$. The distance increases more slowly when $\alpha > 1$, making it hard to distinguish different selection strengths from the data. Since our primary aim is to evaluate which model is better supported by the data, the lower accuracy on the inference of $\alpha$ is acceptable.

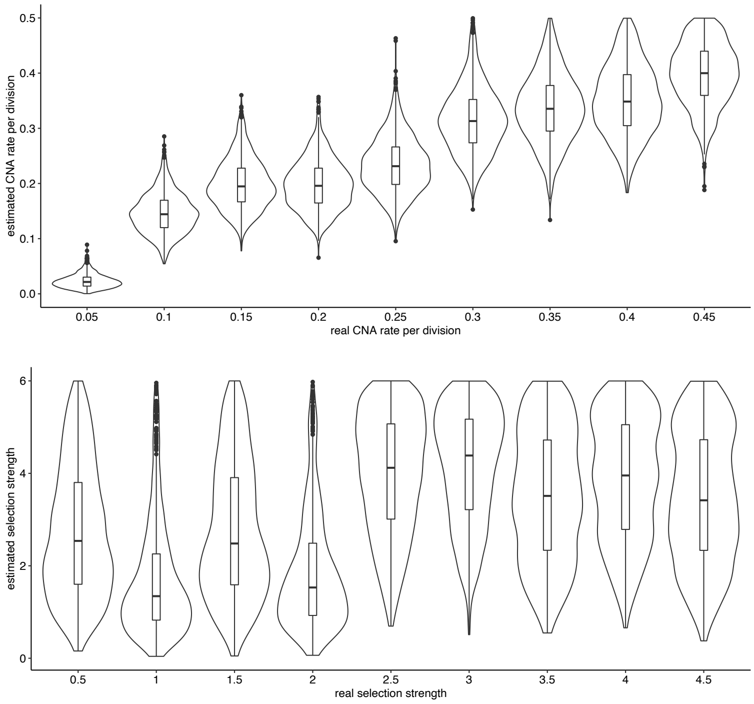

**Supplementary Note Figure 9** The posterior distributions obtained by ABC rejection algorithm when varying true parameters in a range. Top: estimation of $\mu$ under the model without negative selection; Middle: estimation of $\alpha$ assuming $\mu = 0.2$ under the model with negative selection.

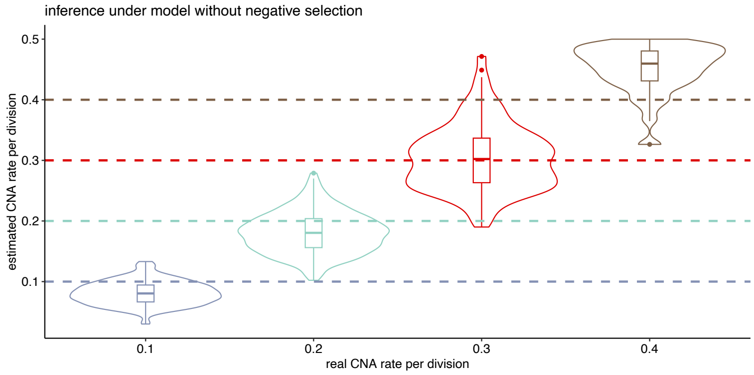

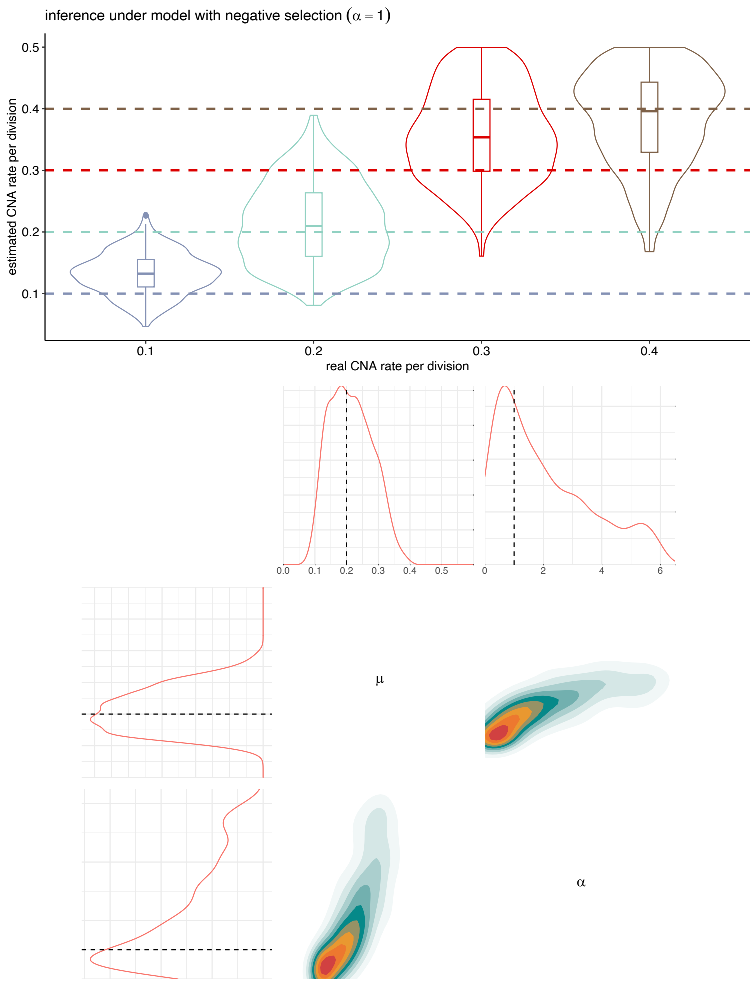

**Supplementary Note Figure 10** The posterior distributions obtained by ABC SMC algorithm when varying true parameters in a range. Top: estimation of $\mu$ under the model without negative selection; Middle: estimation of $\mu$ assuming $\alpha = 1$ under the model with negative selection; Bottom: The correlation of $\mu$ and $\alpha$ under the model with negative selection when $\alpha = 1$. Dashed line: true values used for generating the data.

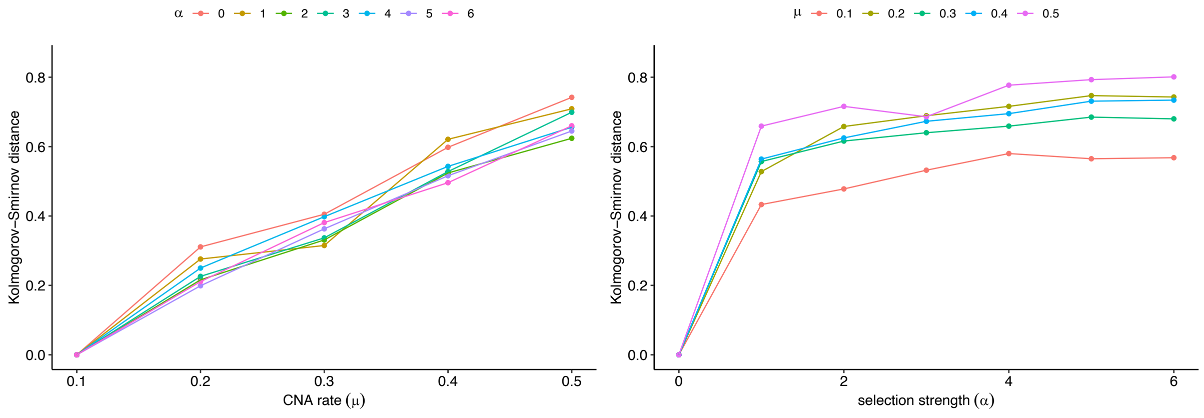

**Supplementary Note Figure 11** Sensitivity analysis on the power of inferring $\mu$ and $\alpha$.

### 4 Results on real data

#### 4.1 Summary statistics

We first computed the summary statistics from real data (Supplementary Note Figure 12). To see how different choices of threshold in excluding focal CNAs affect the values of summary statistics, we did computation by excluding CNAs of at least $n$ bins, where $n$ is from 4 to 10. The choice of $n$ = 8 should exclude enough noise and avoid missing information for some patients (e.g. C277 and C282).

#### 4.2 Choice of $N$

The distribution of the number of divisions for a single gland of birth rate $b$ reaching size $N$ at time $t$ follows a Poisson distribution with mean $2bt$ [2]. This means that on average about 14 fissions occur before the simulation stops. Assuming that one million glands exist in the real tumour, there are about twice as many fissions under the same mutation rate as simulated data and hence more mutations are accumulated. Therefore, the mutation rate tends to be overestimated with simulating 1000 glands. Suppose the mutation rate is $\mu_g$ and the estimated mutation rate is $\mu_g{}'$, then $\mu_g' = \beta\mu_g$, where $\beta$ should be around 2 ($\frac{log(1e6)}{log(1e3)}$).

To check the scale of overestimation in fitting real data with a larger number of glands, we simulated 100,000 glands with mutation rate 0.2 under the model without negative selection based on the karyotypes of different patients. Then we used ABC SMC to infer mutation rate $\mu$ under the model without negative selection with prior $\mu \sim Unif(0, 0.5)$ and target tolerance 0.01. As expected, the results suggest the estimated mutation rates are about twice the real mutation rate (Supplementary Note Figure 13).

We also simulated 100,000 glands with $\mu = 0.2$ and $\alpha = 2$ under the model with negative selection based on the data of patient C282. Negative selection causes less diversity in the target data. To explain the low diversity, mutation rates accepted in ABC tend to be very small. It is not meaningful to accept these very low mutation rates, because simulating a smaller number of glands should overestimate the true mutation rate. To resolve this issue, we imposed a lower limit on the prior of $\mu$ ($\mu_l$) and set the higher limit to 0.5. We did inferences with ABC SMC using different values of $\mu_l$, prior of $\alpha \sim Unif(0{,}6)$ and target tolerance 0.01. Then we computed DIC under the model without and with negative selection (Supplementary Note Figure 14). Since $\mu$ is more likely to be underestimated under the model without negative selection, the correct model was not detected until $\mu_l$ is higher than the real mutation rate.

##### 4.3 Choice of lower limit of $\mu$

To obtain reasonable results from real data, we used known mutation rates per cell division to impose a lower limit of $\mu$. Because our model is at gland level, it is important to understand how the mutation rate at gland level $\mu_g$ correlates with that at cell level $\mu_c$. We defined an effective cell division as a division whose decedents and accompanying mutations got fixed in the gland. Suppose there are $e$ effective cell divisions before a gland fission event, then $\mu_g = \mu_c\, e$, where $\mu_g$ is the CNA rate at gland level and $\mu_c$ is the CNA rate at cell level. Here, $e$ is the number of cell divisions which generated the mutated stem cell that finally takes over the gland. Therefore, $e \geq 1$. Since later CNAs are at a low frequency and cannot be detected with bulk-sequencing, we only consider CNAs that are fixed from the most recent common ancestor of the cells in the gland, then $e = 1$ and hence $\mu_g = \mu_c$.

The mean CNA rate (sum of chromosomal and arm level CNA rate) per cell division estimated based on Live-seq data from patient tumour-derived organoid is typically around 0.2 per cell division, with the minimal rate being around 0.04 [5]. This CNA rate is likely to be an underestimate since only reliable reciprocal CNAs were considered in the inference. To ensure that $\mu_g$ estimated by ABC SMC is not too low to be unrealistic, we constrained $\mu_g$ be at least twice the minimal $\mu_c$.

##### 4.4 Model selection with ABC SMC

We compared the real summary statistics with summary statistics obtained from simulations to get a rough range of mutation rate. The results suggest the mutation rates of real data are more likely to be under 0.5 when being fitted by the simulations. Therefore, we first set $\mu \sim Unif(0.08, 0.5)$ and $\alpha \sim Unif(0, 6)$ as prior distributions for inference on real data. We set target tolerance to 0.001, but none of the runs reached this value and they all stopped when the maximum number of mutations was larger than $10^6$. In the ABC SMC program, it may take more than $10^6$ simulations to obtain the specified number (500) of particles in the last population, because the checking of condition (that the number of total simulations is smaller than the maximum number) is done after the last population is obtained (Supplementary Note Table 2). The changes of tolerances with populations are shown in Supplementary Note Figure ***15***.

The marginal density plots for parameters under the model with negative selection are shown in Supplementary Note Figure 16. As expected, we see moderate positive correlations between the posteriors for $\mu$ and $\alpha$, since a higher mutation rate can be balanced by stronger negative selection. According to DIC values (Supplementary Note Figure 17), the model with negative selection is much better supported in two out of six patients and it is hard to distinguish for two other patients. We also checked the posterior predictive distributions for inference under both models (Supplementary Note Figure 18).

We did some other model checks to validate the fitted models. First, we simulated the data with posterior means of the parameters under both models. It seems clear that the data simulated under the model supported by DIC are more similar to real data (Supplementary Note Figure 19). Second, we did the inference by fixing $\alpha = 1$ since our model cannot accurately infer $\alpha$. The model selection results with DIC show similar conclusions as when $\alpha$ is also estimated, suggesting our results are not biased by the estimation of $\alpha$, as expected (Supplementary Note Figure 20).

As 0.08 is a very conserved lower limit of $\mu$, we used a slightly larger lower limit on the prior of $\mu$ ($\mu \sim Unif(0.1, 0.5)$) and the same other parameters in ABC SMC program, the support for detection model is stronger, as expected (Supplementary Note Figure 21). The estimated parameter values are similar as those when the prior of $\mu$ is $Unif(0.08, 0.5)$.

We also applied summary statistics set 4 when running ABC SMC program on real data to see how it changes the inference (the same other parameters). With this less conservative set of summary statistics, the model with negative selection is better supported in all patients (Supplementary Note Figure 22).

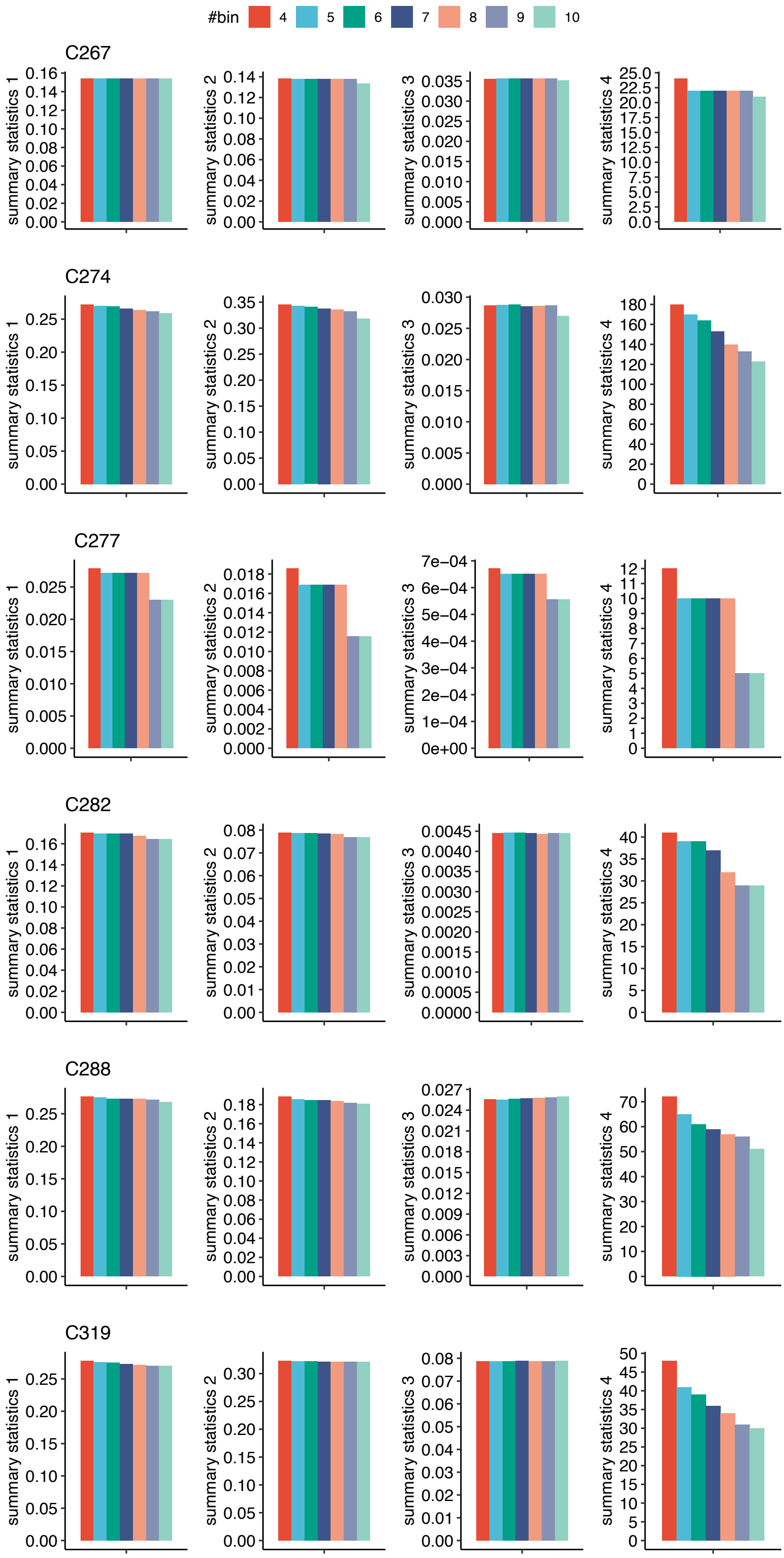

**Supplementary Note Figure 12** The summary statistics computed from real data for each patient when excluding focal CNAs of different sizes.

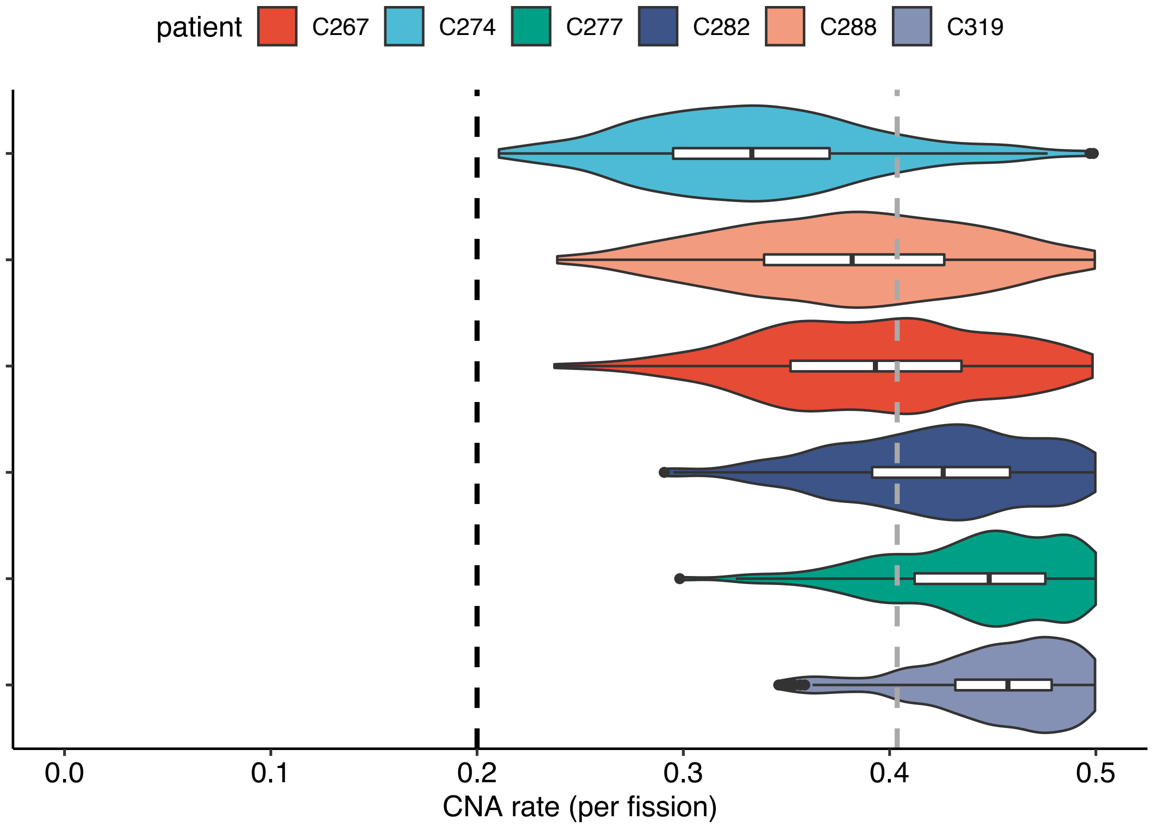

**Supplementary Note Figure 13** The posterior distributions of CNA rate inferred with ABC SMC for simulated target dataset with $10^5$ glands ($\mu = 0.2$) based on the karyotypes of different patients. Black dashed line: real $\mu$. Grey dashed line: weighted mean of estimated $\mu$.

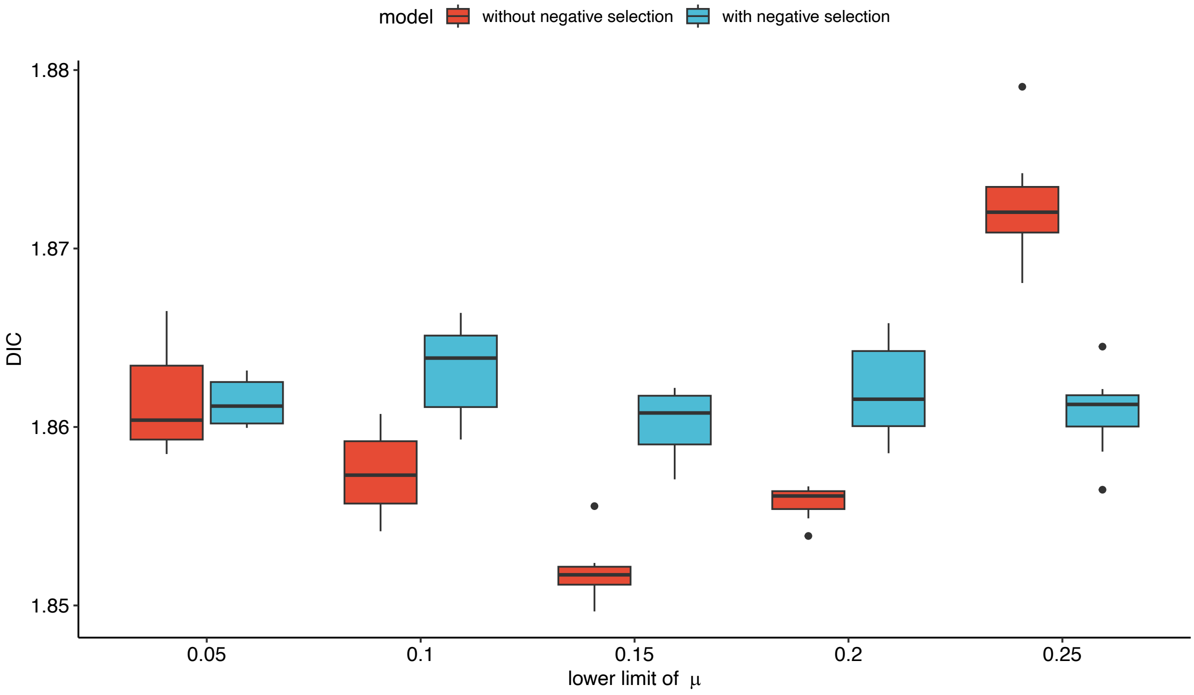

**Supplementary Note Figure 14** Model selection with DIC (10 replicates for each case) on simulated target data with the prior of $\mu \sim Unif(\mu_l, 0.5)$, where $\mu_l = \{0.05, 0.1, 0.15, 0.2, 0.25\}$.

**Supplementary Note Table 2** The number of simulations when applying ABC SMC for each patient under the model without and with negative selection when target tolerance is 0.001 and the priors are $\mu \sim Unif(0.08, 0.5)$ and $\alpha \sim Unif(0, 6)$.

| Patient | #simulations under the model without negative selection | #simulations under the model with negative selection |
|---|---|---|
| C267 | 1,323,629 | 1,754,945 |
| C274 | 2,542,014 | 1,039,805 |
| C277 | 1,008,964 | 1,798,795 |
| C282 | 2,811,294 | 1,169,039 |
| C288 | 1,240,900 | 2,282,509 |
| C319 | 2,727,334 | 2,193,467 |

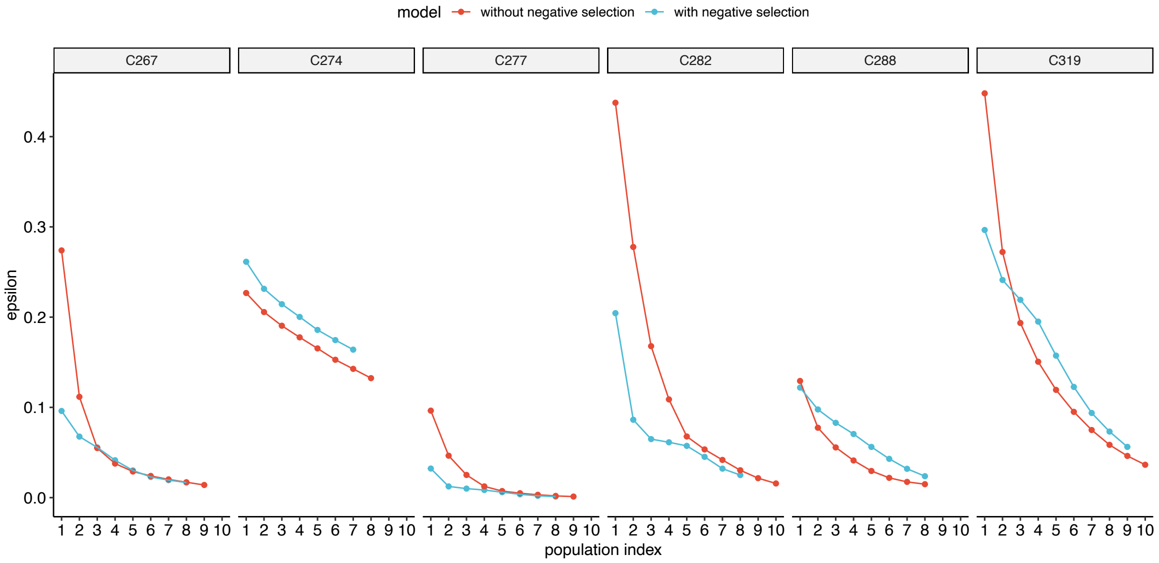

**Supplementary Note Figure 15** The changes of tolerances with populations for each patient in running of the ABC SMC algorithm under the model without and with negative selection when target tolerance is 0.001 and the priors are $\mu \sim Unif(0.08, 0.5)$ and $\alpha \sim Unif(0, 6)$.

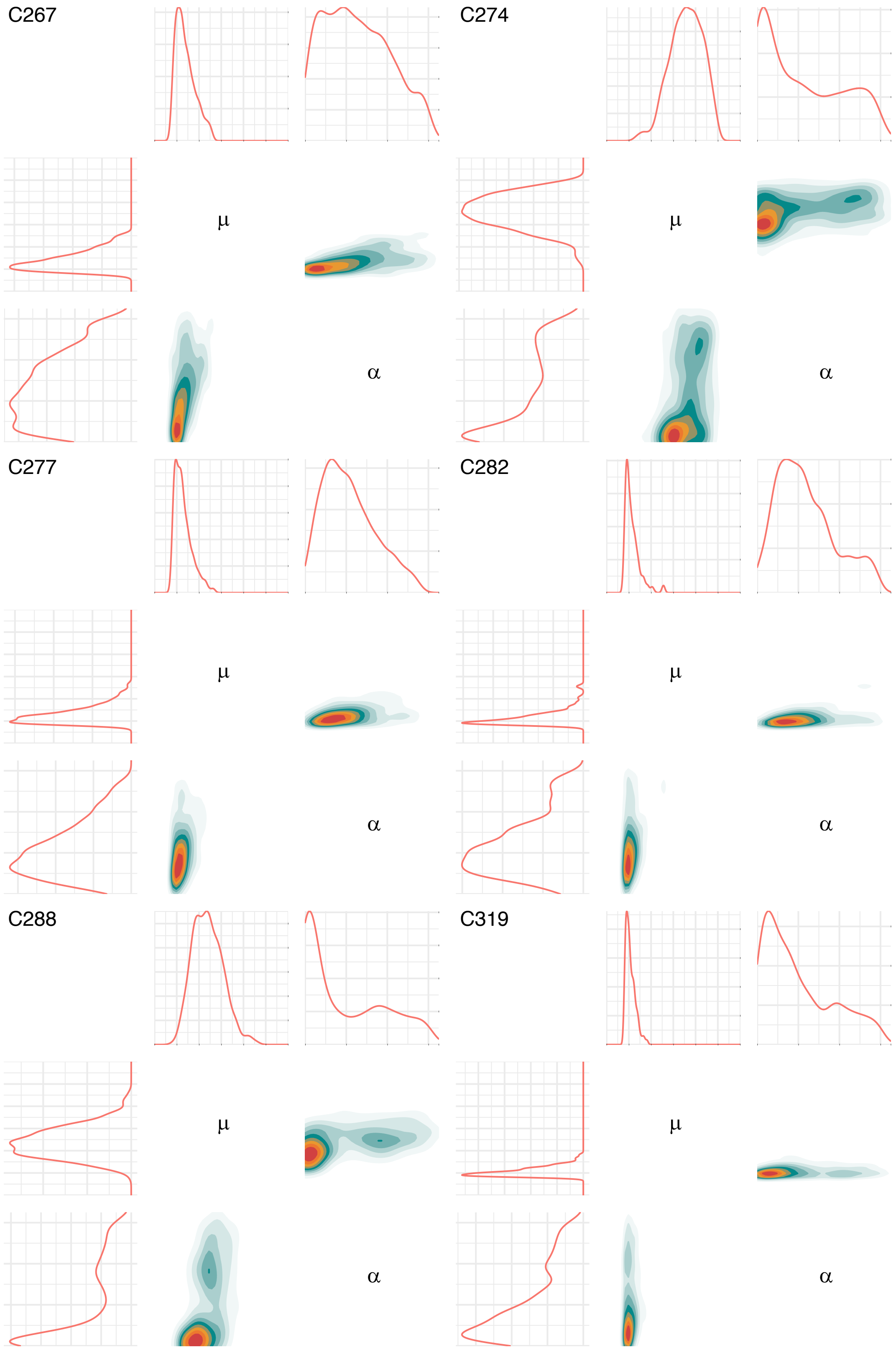

**Supplementary Note Figure 16** The marginal densities of parameters inferred under the model with negative selection when target tolerance is 0.001 and the priors are $\mu \sim Unif(0.08, 0.5)$ and $\alpha \sim Unif(0, 6)$.

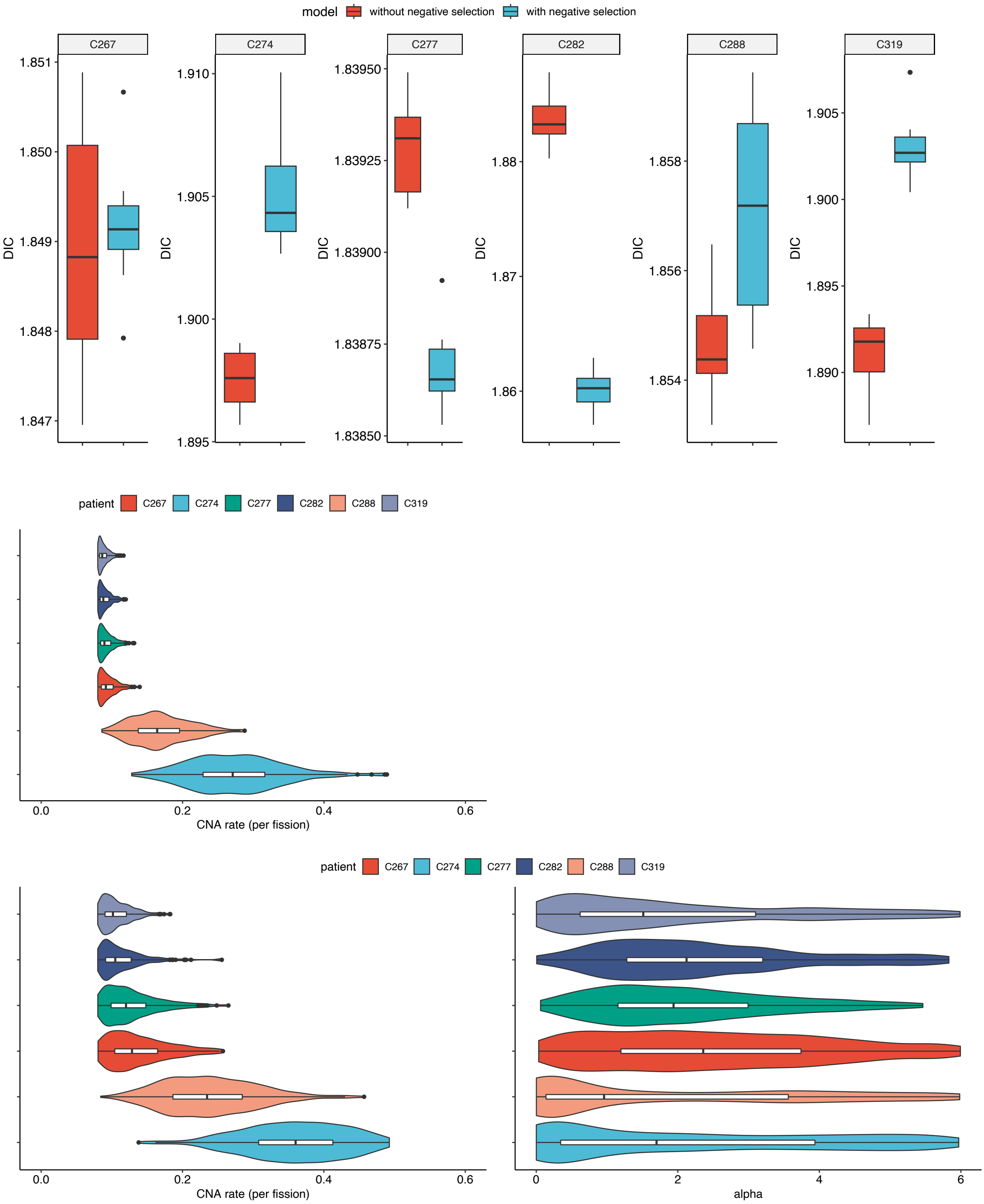

**Supplementary Note Figure 17** Model selection results with DIC (10 replicates for each case) and the posterior distributions of parameters inferred under the model without and with negative selection when target tolerance is 0.001 and the priors are $\mu \sim Unif(0.08, 0.5)$ and $\alpha \sim Unif(0, 6)$.

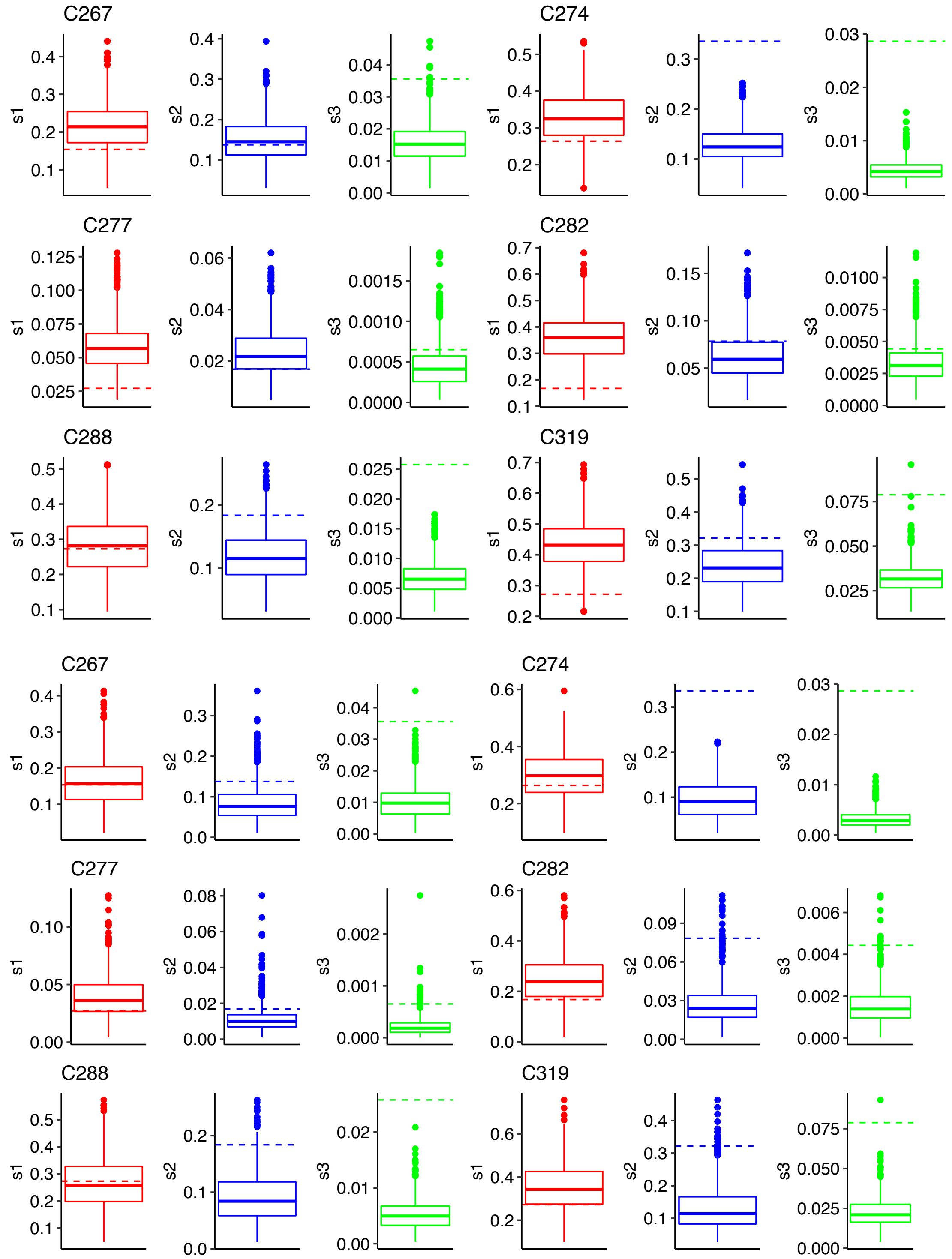

**Supplementary Note Figure 18** The posterior predictive distributions for the three summary statistics used for inference under the model without (top) and with (bottom) negative selection when target tolerance is 0.001 and the priors are $\mu \sim Unif(0.08, 0.5)$ and $\alpha \sim Unif(0, 6)$. The dashed lines indicate the observed summary statistics.

C267 (model without negative selection)

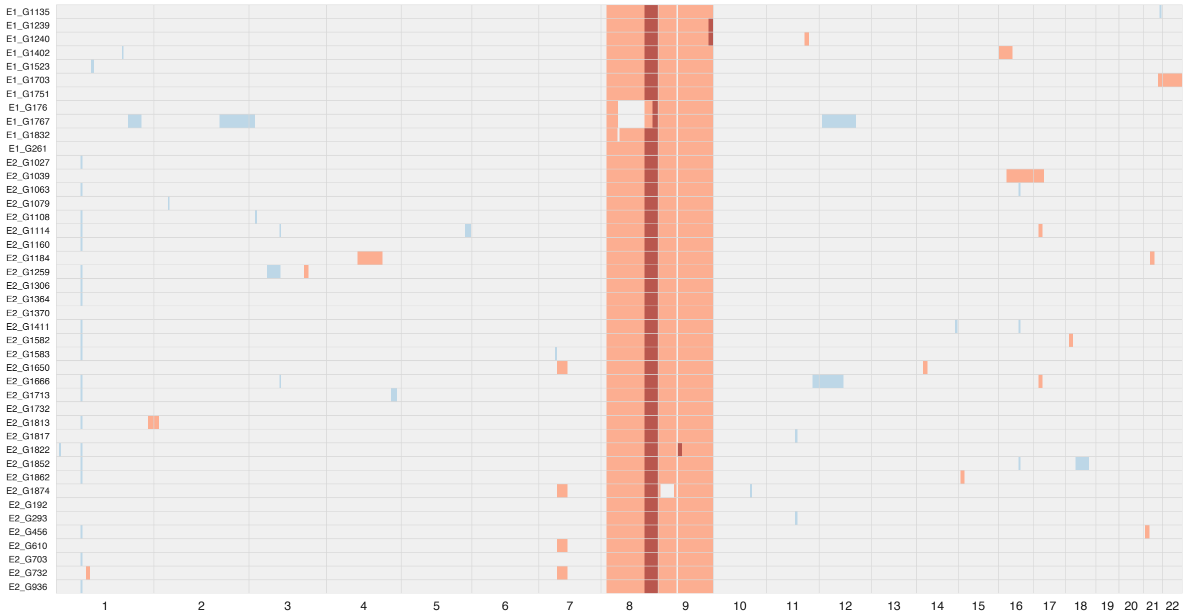

C267 (model with negative selection)

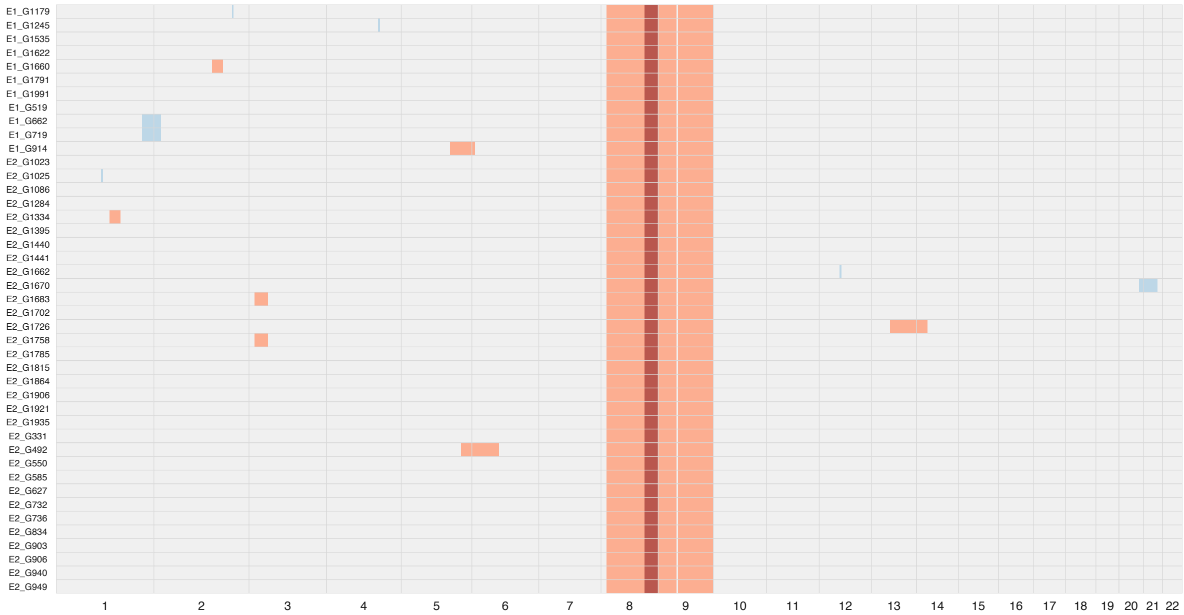

C274 (model without negative selection)

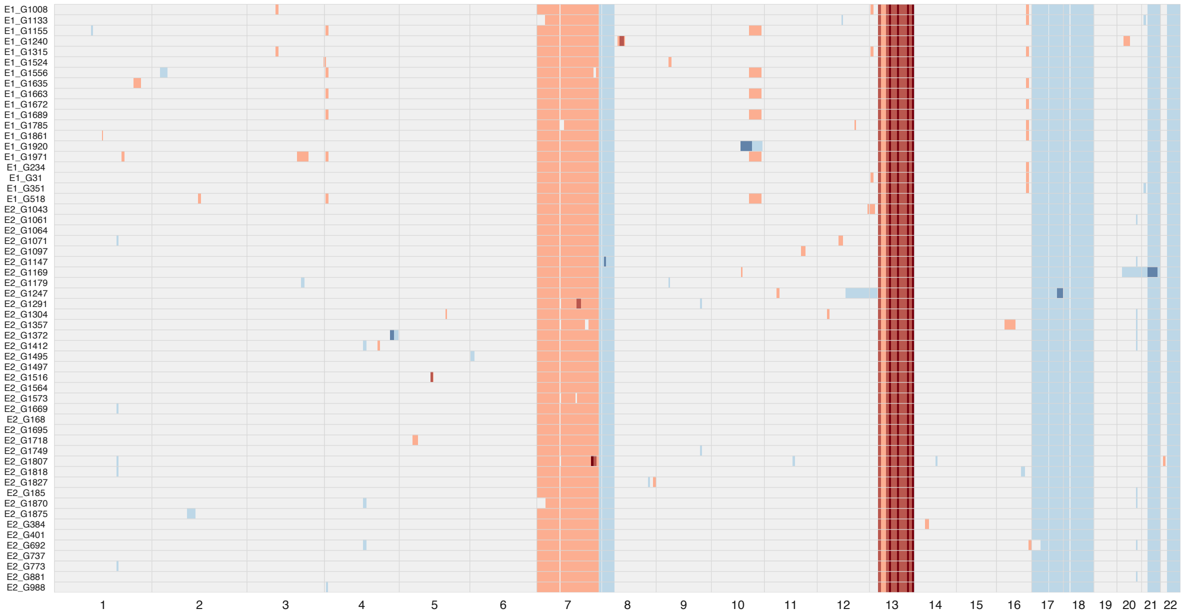

C274 (model with negative selection)

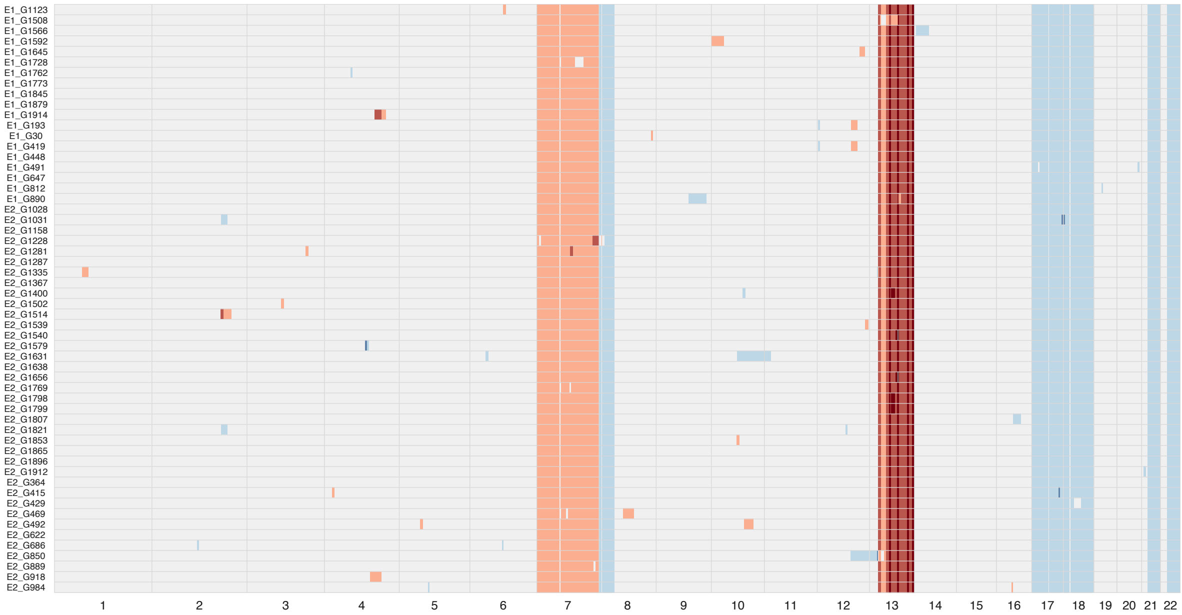

C277 (model without negative selection)

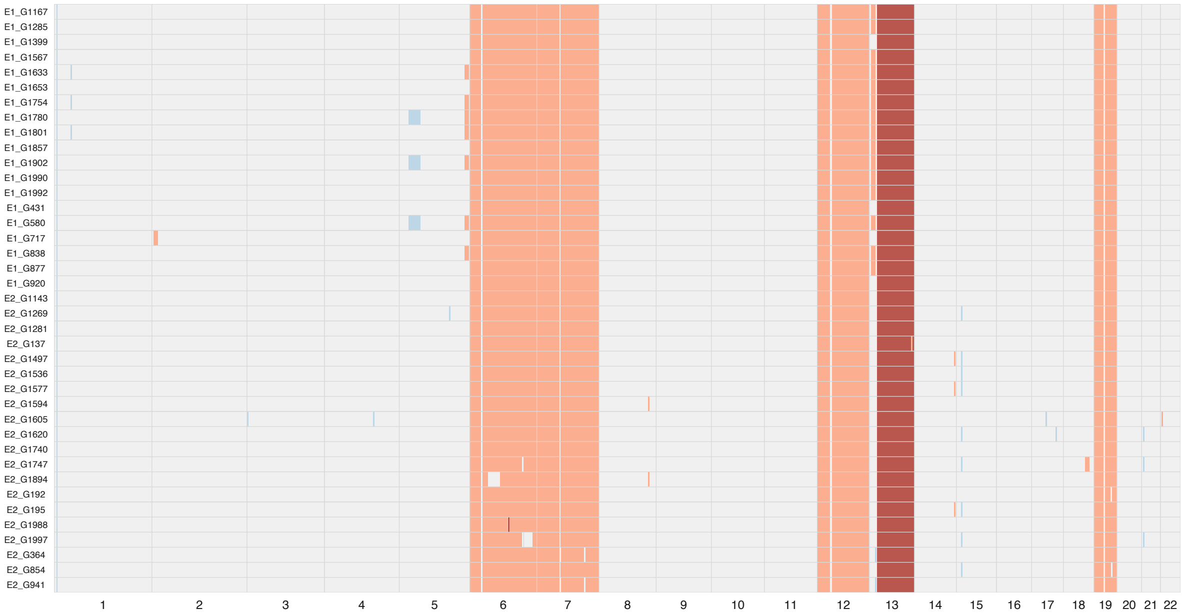

C277 (model with negative selection)

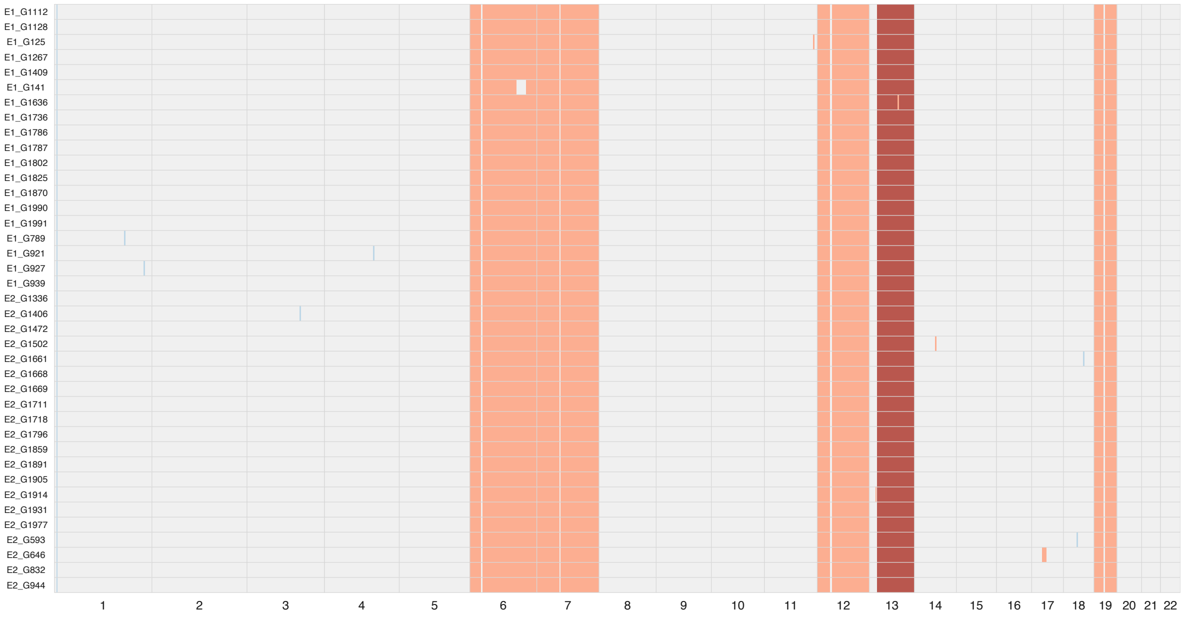

C282 (model without negative selection)

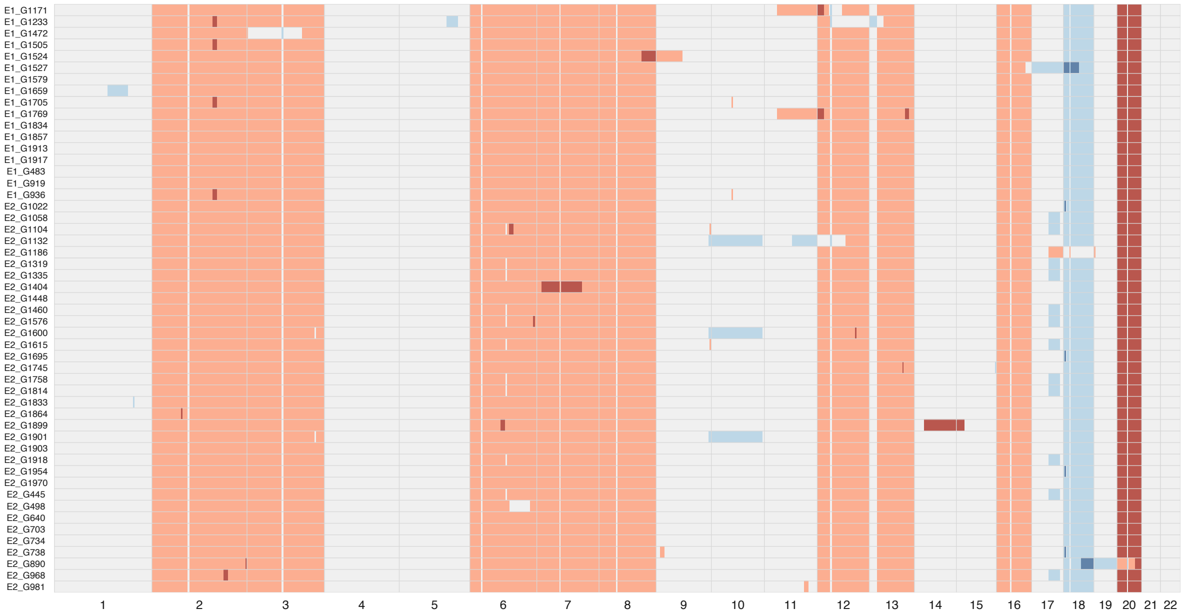

C282 (model with negative selection)

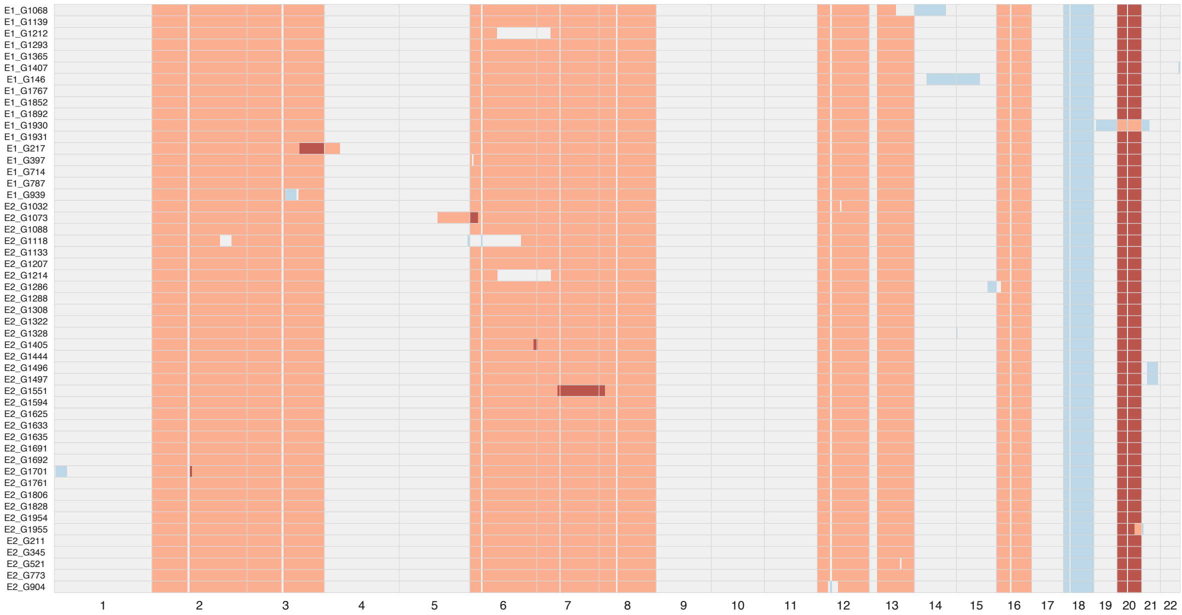

C288 (model without negative selection)

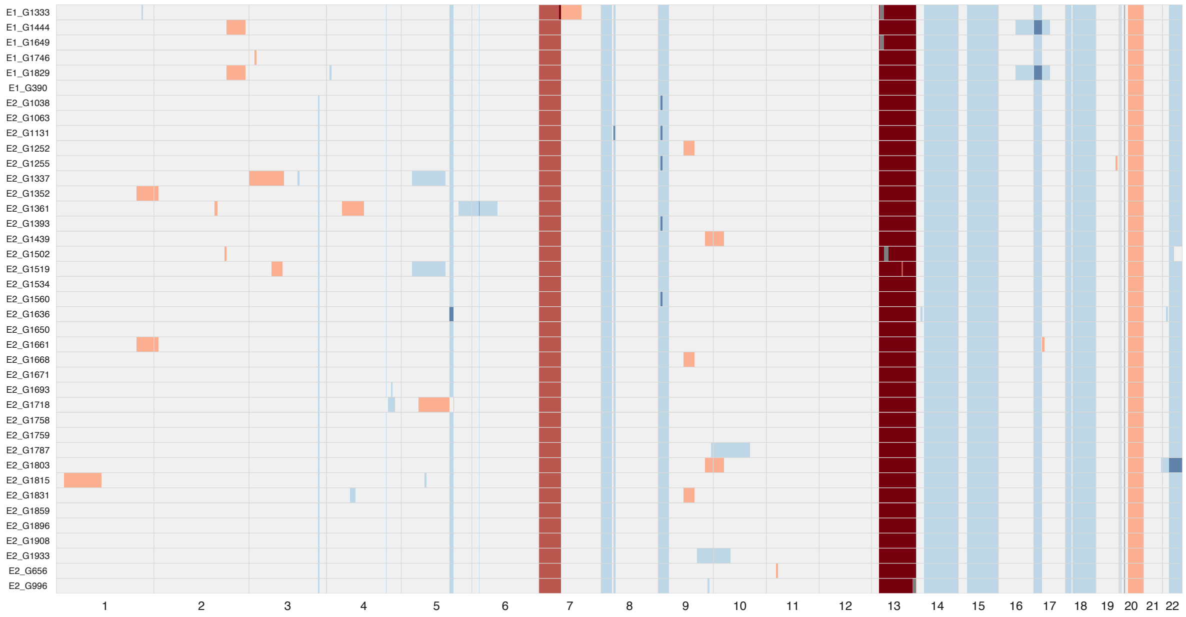

C288 (model with negative selection)

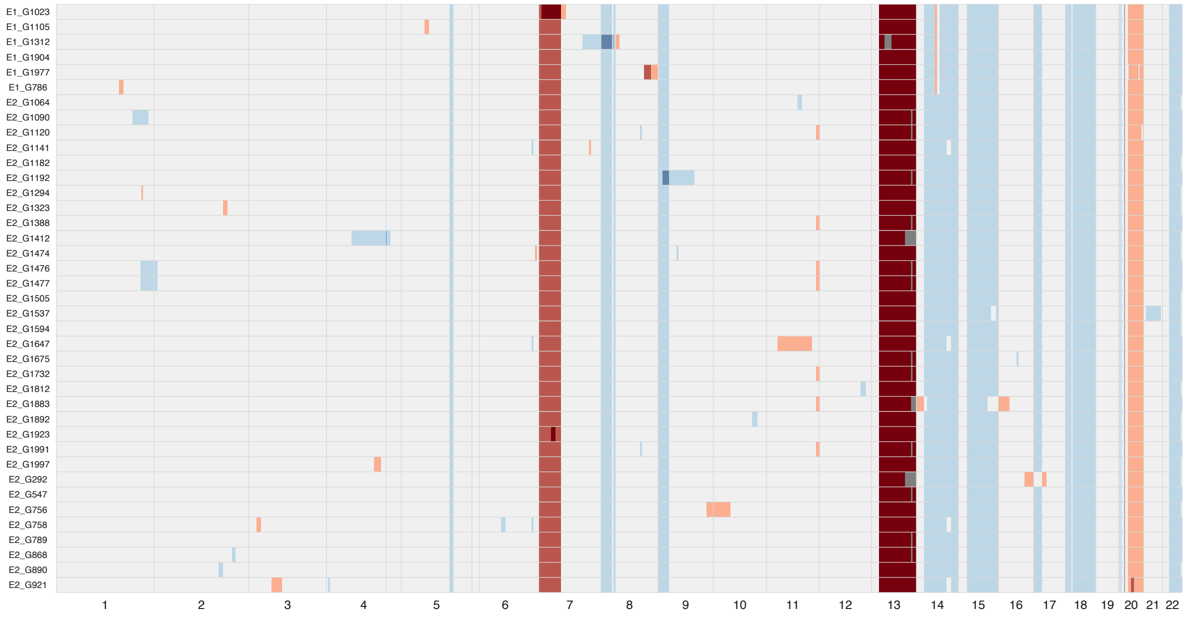

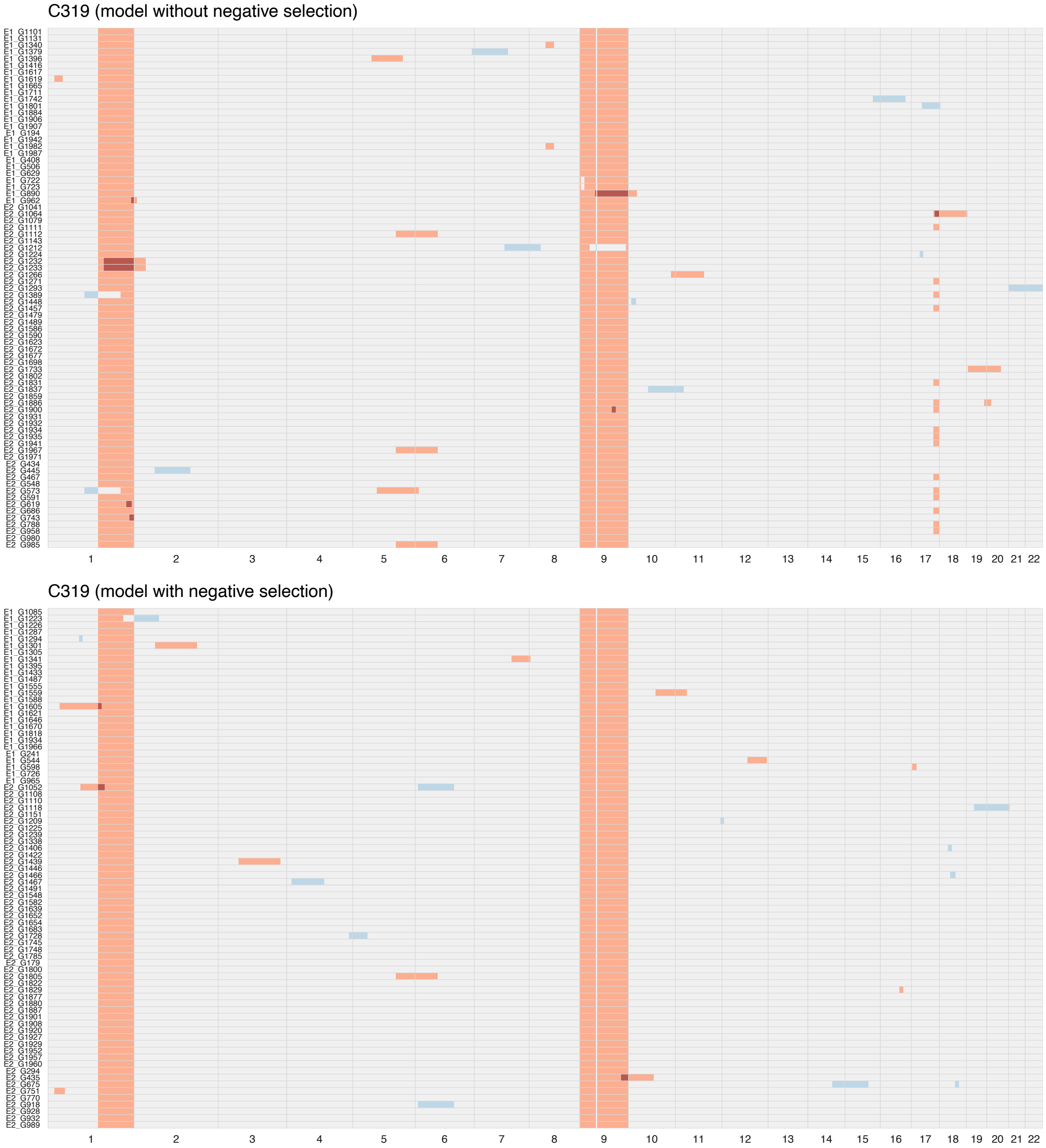

**Supplementary Note Figure 19** Simulated copy number profiles (Grey: 2N; Light red: 3N; Dark red: 4N; Light blue: 1N) for six patients with posterior means of parameters estimated under the model without and with negative selection when target tolerance is 0.001 and the priors are $\mu \sim Unif(0.08, 0.5)$ and $\alpha \sim Unif(0, 6)$. The labels on the left denote the IDs of simulated glands, where E represents the side that a gland is sampled from.

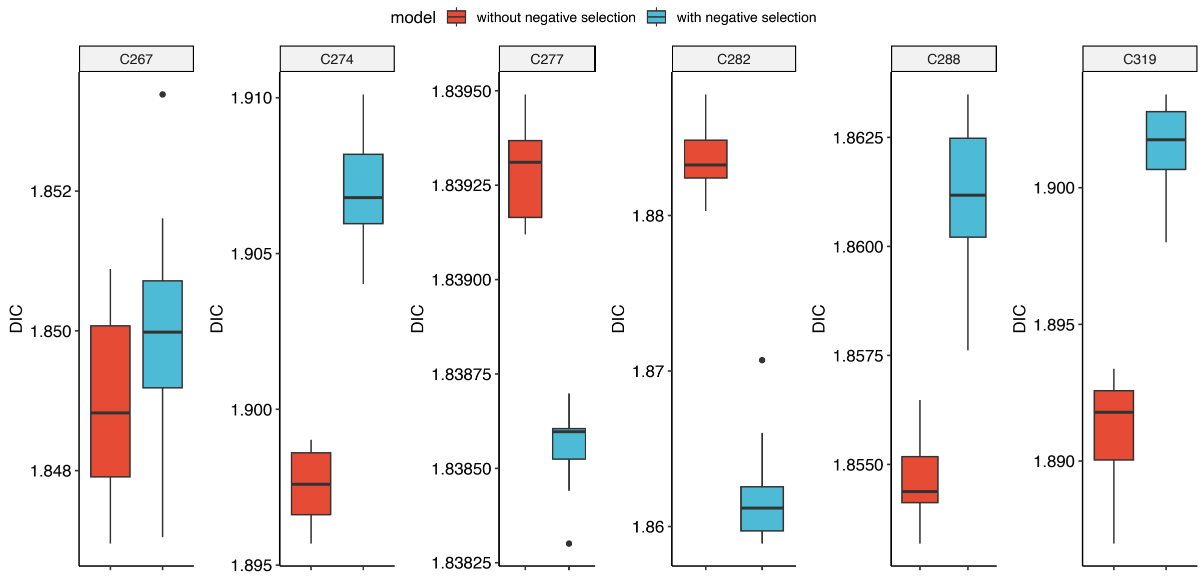

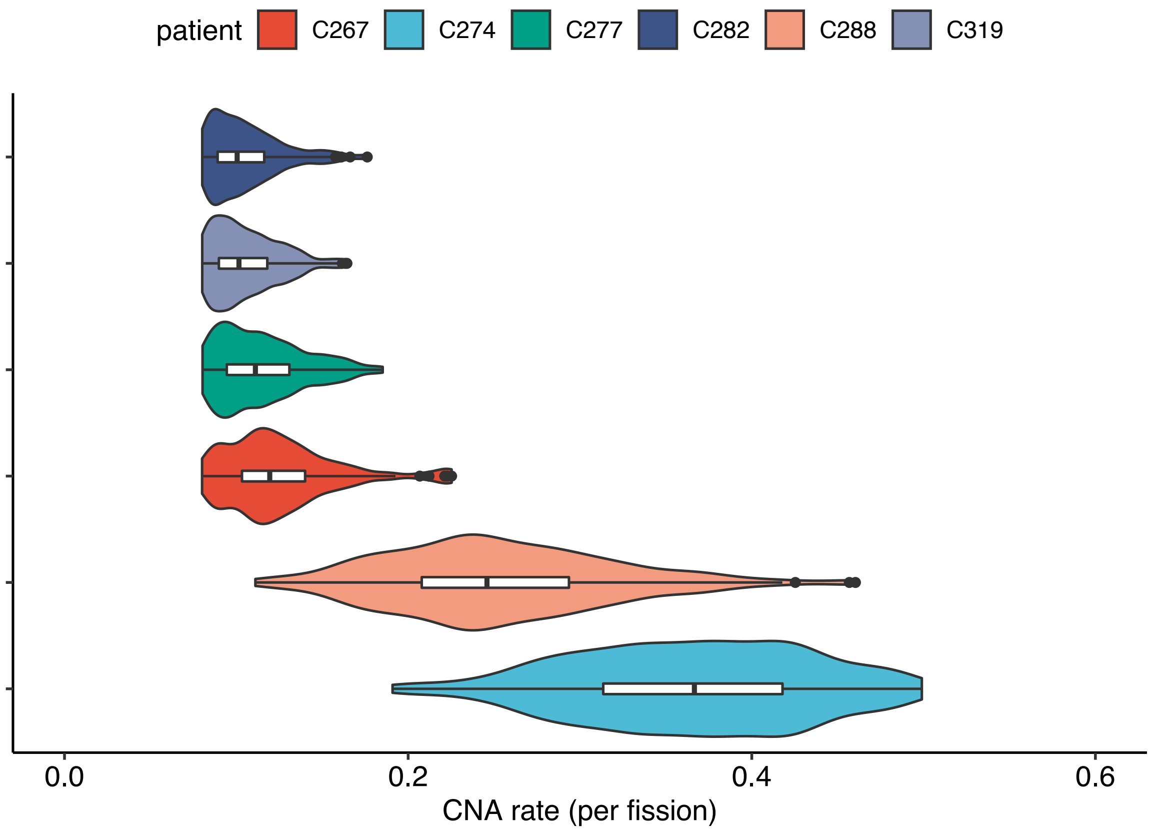

**Supplementary Note Figure 20** Model selection results with DIC (10 replicates for each case) and the posterior distributions of CNA rate when $\alpha = 1$ under the model with negative selection, target tolerance is 0.001 and the prior is $\mu \sim Unif(0.08, 0.5)$.

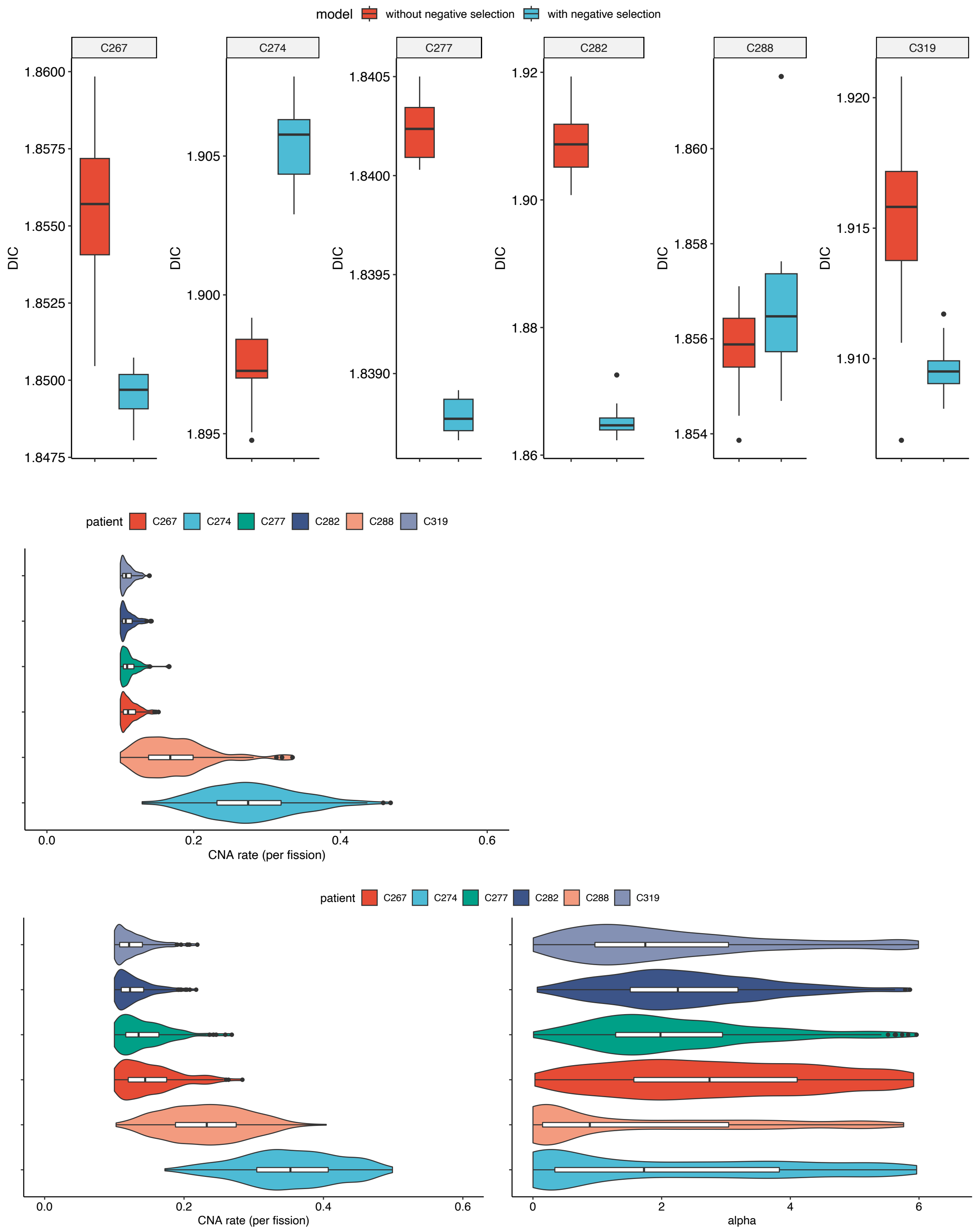

**Supplementary Note Figure 21** Model selection results with DIC (10 replicates for each case) and the posterior distributions of parameters inferred under the model without and with negative selection when target tolerance is 0.001 and the priors are $\mu \sim Unif(0.1, 0.5)$ and $\alpha \sim Unif(0, 6)$.

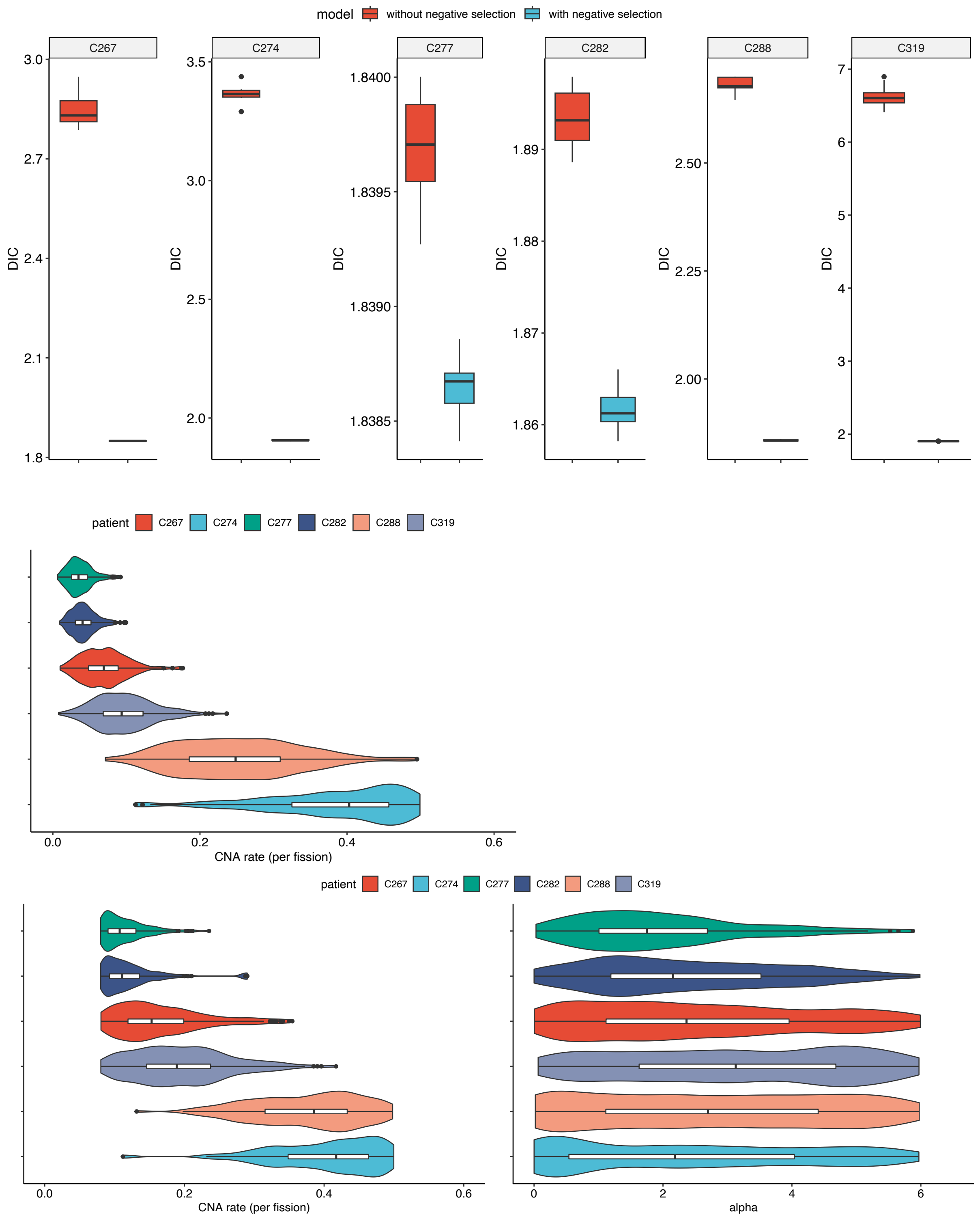

**Supplementary Note Figure 22** Model selection results with DIC (10 replicates for each case) and the posterior distributions of parameters inferred under the model without and with negative selection when summary statistics set 4 is used, target tolerance is 0.001, and the priors are $\mu \sim Unif(0.08, 0.5)$ and $\alpha \sim Unif(0, 6)$.
